# Identification and characterization of novel filament-forming proteins in cyanobacteria

**DOI:** 10.1101/674176

**Authors:** Benjamin L. Springstein, Christian Woehle, Julia Weissenbach, Andreas O. Helbig, Tal Dagan, Karina Stucken

**Affiliations:** Institute of General Microbiology, Christian-Albrechts-Universität zu Kiel, Kiel, Germany; Institute for Experimental Medicine, Christian-Albrechts-Universität zu Kiel, Kiel, Germany; Department of Food Engineering, University of La Serena, La Serena, Chile

## Abstract

Filament-forming proteins in bacteria function in stabilization and localization of proteinaceous complexes and replicons; hence they are instrumental for myriad cellular processes such as cell division and growth. Here we present two novel filament-forming proteins in cyanobacteria. Surveying cyanobacterial genomes for coiled-coil-rich proteins (CCRPs) that are predicted as putative filament-forming proteins, we observed a higher proportion of CCRPs in filamentous cyanobacteria in comparison to unicellular cyanobacteria. Using our predictions, we identified nine protein families with putative intermediate filament (IF) properties. Polymerization assays revealed four proteins that formed polymers *in vitro* and three proteins that formed polymers *in vivo*. Fm7001 from *Fischerella muscicola* PCC 7414 polymerized *in vitro* and formed filaments *in vivo* in several organisms. Additionally, we identified a tetratricopeptide repeat protein - All4981 - in *Anabaena* sp. PCC 7120 that polymerized into filaments *in vitro* and *in vivo*. All4981 interacts with known cytoskeletal proteins and is indispensable for *Anabaena* viability. Although it did not form filaments *in vitro*, Syc2039 from *Synechococcus elongatus* PCC 7942 assembled into filaments *in vivo* and a Δ*syc2039* mutant was characterized by an impaired cytokinesis. Our results expand the repertoire of known prokaryotic filament-forming CCRPs and demonstrate that cyanobacterial CCRPs are involved in cell morphology, motility, cytokinesis and colony integrity.

## Introduction

Species in the phylum Cyanobacteria present a wide morphological diversity, ranging from unicellular to multicellular organisms. Unicellular cyanobacteria of the *Synechocystis* and *Synechococcus* genera are characterized by a round or rod-shaped morphology, respectively, and many strains are motile. Species of the Nostocales order are multicellular and differentiate specialized cells, known as heterocysts, which fix atmospheric nitrogen under aerobic conditions. Within the Nostocales, species of the Nostocaceae (e.g., *Anabaena*, *Nostoc*) form linear trichomes, while cells in the Hapalosiphonaceae and Chlorogloepsidaceae divide in more than one plane to form true-branching trichomes as in *Fischerella* or multiseriate trichomes (more than one filament in a row) as in *Chlorogloeopsis* (Rippka *et al.*, 1979). Notably, cells within a single trichome of a multicellular cyanobacterium can differ in size, form or cell wall composition (Rippka *et al.*, 1979). Cells in the *Anabaena* sp. PCC 7120 (hereafter *Anabaena*) trichome are linked by a shared peptidoglycan sheet and an outer membrane (Wilk *et al.*, 2011). *Anabaena* cells communicate and exchange nutrients through intercellular cell-cell connections, called septal junctions, which are thought to comprise the septal junction proteins SepJ, FraC and FraD (Herrero, Stavans and Flores, 2016; Weiss *et al.*, 2019). SepJ is essential for the multicellular phenotype in *Anabaena* (Flores *et al.*, 2007; Nayar *et al.*, 2007).

Studies of the molecular basis of cyanobacterial morphogenesis have so far focused on the function of FtsZ and MreB, the prokaryotic homologs of tubulin and actin, respectively (Wagstaff and Löwe, 2018). FtsZ functions in a multi-protein complex called the divisome, and is known as a key regulator of cell division and septal peptidoglycan (PG) biogenesis (Bi and Lutkenhaus, 1991; Wagstaff and Löwe, 2018). FtsZ has been shown to be an essential cellular protein in *Anabaena* and in the coccoid cyanobacterium *Synechocystis* sp. PCC 6803 (hereafter *Synechocystis*) (Zhang *et al.*, 1995). The FtsZ cellular concentration in *Anabaena* is tightly controlled by a so far undescribed protease (Lopes Pinto *et al.*, 2011). Apart from its function in cell division, the FtsZ-driven divisome also mediates the localization of SepJ (Ramos-León *et al.*, 2015). MreB functions in a multi-protein complex called the elongasome, where it is a key mediator of longitudinal PG biogenesis that controls the cell shape (Jones, Carballido-López and Errington, 2001; Wagstaff and Löwe, 2018). In cyanobacteria, MreB plays a role in cell shape determination in *Anabaena*, nonetheless, it is not essential for cell viability (Hu *et al.*, 2007). In contrast, in *Synechococcus* sp. PCC 7942 (hereafter *Synechococcus*) MreB is essential, where partially segregated mutants display a coccoid morphology resembling the morphology of *E. coli mreB* deletion strains (Kruse, Bork-Jensen and Gerdes, 2005; Jain, Vijayan and O’Shea, 2012).

Proteins resembling the eukaryotic intermediate filaments (IFs) have been discovered in several bacterial species and were shown to form filaments *in vitro* and *in vivo* and to impact essential cellular processes (Lin and Thanbichler, 2013). IF proteins exhibit an intrinsic nucleotide-independent *in vitro* polymerization capability that is mediated by the high frequency of coiled-coil-rich regions in their amino acid sequence (Shoeman and Traub, 1993; Fuchs and Weber, 1994; Löwe and Amos, 2009; Wagstaff and Löwe, 2018). Eukaryotic IF proteins are generally characterized by a conserved domain buildup consisting of discontinuous coiled-coil segments that form a central rod domain. This rod domain is N- and C-terminally flanked by globular head and tail domains of variable length (Fuchs and Weber, 1994; Herrmann *et al.*, 1996; Herrmann and Aebi, 2004). Crescentin is a bacterial IF-like CCRP from *Caulobacter crescentus*, which exhibits a striking domain similarity to eukaryotic IF proteins. Crescentin filaments that align at the inner cell curvature are essential for the typical crescent-like cell shape of *C. crescentus*; possibly, by locally exuding a constriction force which coordinates the MreB-driven peptidoglycan (PG) synthesis machinery (Ausmees, Kuhn and Jacobs-Wagner, 2003; Cabeen *et al.*, 2009; Charbon, Cabeen and Jacobs-Wagner, 2009). Reminiscent of eukaryotic IF proteins, Crescentin was found to assemble into filamentous structures *in vitro* in a nucleotide-independent manner (Ausmees, Kuhn and Jacobs-Wagner, 2003). However, so far no Crescentin homologs have been found in other bacteria, indicating that non-spherical or rod-shaped prokaryotic morphologies are putatively controlled by other polymerizing proteins (Bagchi *et al.*, 2008; Wickstead and Gull, 2011). Apart from Crescentin, many other coiled-coil-rich proteins (CCRPs) with IF-like functions have been identified to polymerize into filamentous structures and to perform cytoskeletal-like roles; however, none of them resembled the eukaryotic IF domain architecture (reviewed by Lin & Thanbichler, 2013). Examples are two proteins from *Streptomyces coelicolor* whose function has been studied in more detail: FilP and Scy (Bagchi *et al.*, 2008; Walshaw, Gillespie and Kelemen, 2010; Holmes *et al.*, 2013). Gradients of FilP localize at the tip of a growing hyphae and contribute to cellular stiffness (Bagchi *et al.*, 2008). Scy forms patchy clusters at the sites of novel tip-formation and, together with the scaffolding CCRP DivIVA, orchestrates the polar hyphal growth (Holmes *et al.*, 2013). Together with FilP and a cellulose-synthase, these proteins form the polarisome, which guides peptidoglycan biogenesis and hyphal tip growth in *S. coelicolor* (Flärdh *et al.*, 2012; Hempel *et al.*, 2012; Holmes *et al.*, 2013). Another example are four CCRPs in the human pathogen *Helicobacter pylori*, which were found to assemble into filaments *in vitro* and *in vivo*, with a function in determination of the helical cell shape as well as cell motility (Waidner *et al.*, 2009; Specht *et al.*, 2011). Consequently, filament-forming CCRPs with essential cellular functions have been found in numerous prokaryotes having various cellular morphologies. The presence of filament-forming CCRPs in cyanobacteria is so far understudied. Here we search for CCRPs with presumed IF-like functions in cyanobacteria using a computational prediction of CCRPs. Putative filament-forming proteins were further investigated experimentally by structural analyses and *in vitro* and *in vivo* localization assays in morphologically diverse cyanobacteria.

## Results

### Coiled-coil-rich proteins are widespread in cyanobacteria

For the computational prediction of putative filament-forming proteins, we surveyed 364 cyanobacterial genomes including 1,225,314 protein-coding sequences (CDSs) for CCRPs. All CDSs in the cyanobacterial genomes where clustered by sequence similarity into families of homologous proteins (see Methods). The frequency of CCRPs in each CDS was calculated using the COILS algorithm (Lupas, Van Dyke and Stock, 1991). The algorithm yielded a list of 28,737 CDSs with high coiled-coil content (≥80 amino acids in coiled-coil conformation; Supplementary File 1). CCRPs were predicted in 158,466 protein families covering all cyanobacterial species. To examine the overall distribution of CCRPs in cyanobacterial genomes, we investigated 1,504 families of homologous proteins that include at least three CCRP members (Fig. 1). Notably, most protein families (1,142; 76%) include CCRP and non-CCRP members, indicating that protein properties might differ among homologous proteins. The presence/absence pattern of families including CCRPs further shows that those are less abundant in picocyanobacterial genomes (SynProCya group) in comparison to the remaining specie in the phylum. Furthermore, the proportion of CCRPs in the genome is significantly higher in multicellular cyanobacteria in comparison to unicellular cyanobacteria (P=2.65×10^−46^ using Kruskal-Wallis test and Tukey test with α=0.05). This indicates that a high frequency of CCRPs is one characteristic of multicellular cyanobacteria.

**Fig. 1:**
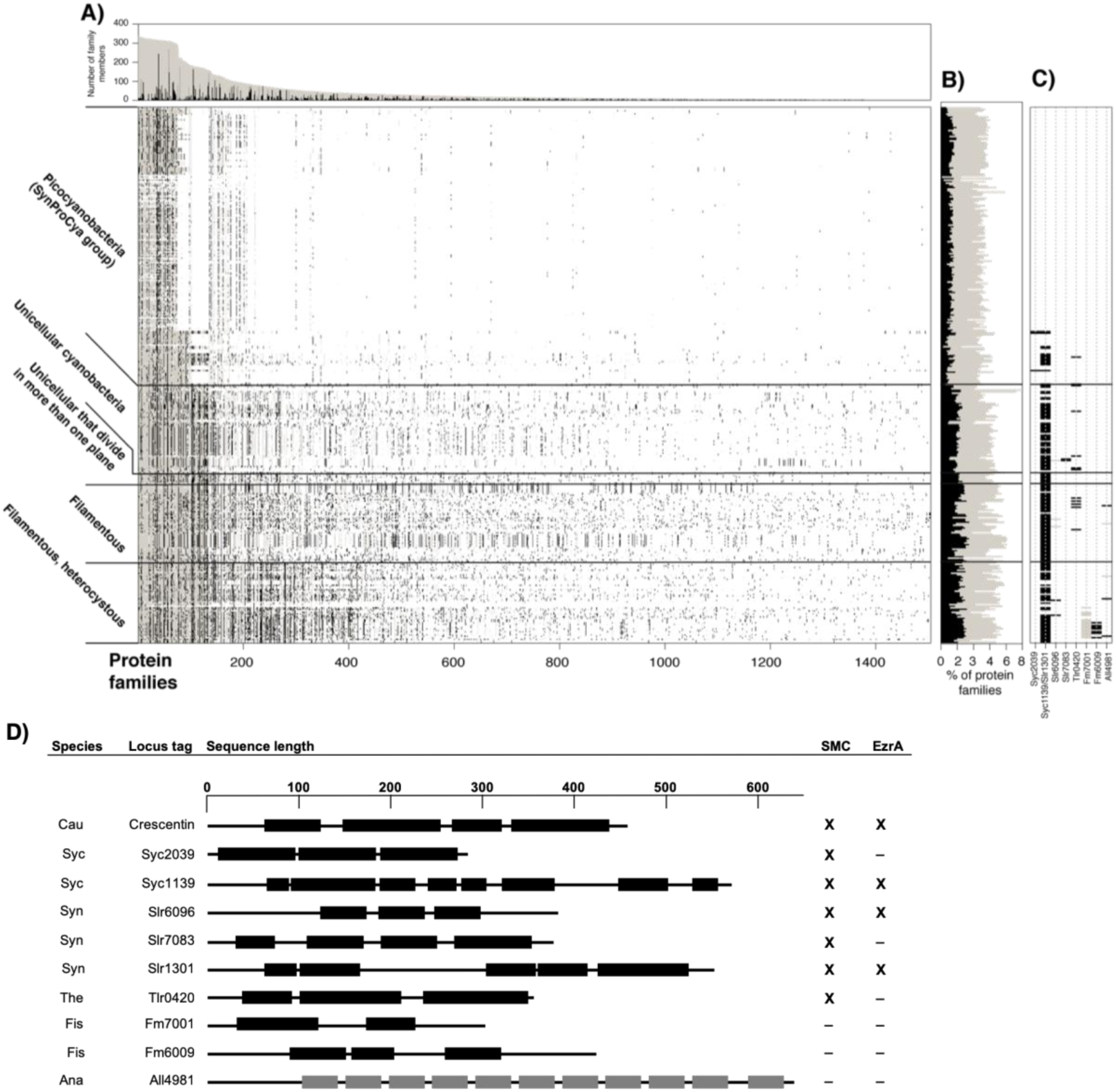
Distribution of CCRP protein families within cyanobacteria. **(A)** Lines in the presence/absence matrix designate cyanobacterial genomes; each column shows a protein family. Gray dots designate any homologous protein in the same protein family and black dots represent CCRP members. Protein families are sorted according to the number of members. Protein family size and the number of CCRP members are presented in a bar graph above. (**B**) The proportion of protein families containing CCRPs (gray) and CCRP proteins (black) in each genome. (**C**) Presence/absence pattern of CCRP candidate protein families. Only protein families with at least three members predicted to be CCRPs are shown. (**D**) Domain prediction of CCRP candidates. Scale on top is given in amino acid residues. Amino acid sequences in coiled-coil conformation are depicted by black bars with non-coiled-coil sequences represented by black lines. Tetratricopeptide repeats (TPR), also predicted by the COILS algorithm, are shown as grey bars. Proteins are given as cyanobase locus tags. Fm7001 and Fm6009 correspond to NCBI accession numbers WP_016868005.1 and WP_020476706, respectively. Abbreviations: Cau: *C. crescentus*; Syc: *Synechococcus*, Syn: *Synechocystis*; Ana: *Anabaena*; The: *Thermosynechococcus elongatus* BP-1; Fis: *Fischerella*. Cyanobacterial CCRPs had conserved domains present in prokaryotic IF-like CCRPs and eukaryotic IF proteins (Supplementary Table 1). Presence of a structural maintenance of chromosomes (SMC) domain or structural similarities to the cell division protein EzrA are marked with “**X**”, absence is indicated with “**-**“. Full list is given in Supplementary Table 1. Note: *Anabaena* CCRPs have been described elsewhere before: *Springstein et al*. (2019), bioRxiv, doi: 10.1101/553073.

For the experimental validation, the complete list of CCRPs was filtered to include candidates from freshwater unicellular and filamentous cyanobacteria that are genetically accessible, including *Thermosynechococcus elongatus* BP-1 (*Thermosynechococccus*), *Synechocystis*, *Synechococcus*, *Anabaena* and *Fischerella muscicola* PCC 7414 (*Fischerella*). In addition to cytoskeleton functions, coiled-coils are common motifs of proteins involved in other cellular processes such as transcription, the extracellular matrix, chemotaxis and host–pathogen interactions (Rackham *et al.*, 2010). Consequently, the remaining CCPRs were further sorted to include proteins having similar properties to known prokaryotic IF-like CCRPs (e.g., crescentin, FilP) and are annotated as hypothetical proteins with an unknown function. Additionally, proteins lacking an unstructured N-terminal head and C-terminal tail domain, which are characteristics of prokaryotic IF-like proteins (Bagchi *et al.*, 2008), were excluded. Furthermore, proteins with an assigned function or predicted to be involved in other cellular processes were excluded (using publicly available online bioinformatic tools: NCBI Blast, NCBI CD search, PSORTb, TMHMM, InterPro, PSIPRED and I-TASSER). In the screening for protein characteristics and annotation, Crescentin, FilP and other eukaryotic IF proteins (e.g., Vimentin and Desmin) were chosen as reference for our predictions, where proteins displaying similar results were favored. An additional *Fischerella* CDS, Fm7001, was added to the list as earlier analyses suggested that it has a cell shape-determining function. The preliminary filtration resulted in a list of nine candidates, which we investigated experimentally here (Fig. 1C,D and Supplementary Table 1).

Candidate coding sequences varied in size and ranged from ca. 280 amino acids (Synpcc7942_2039, abbreviated Syc2039) to ca. 650 amino acids (All4981). The coiled-coil domain distribution was variable among the candidates in both coiled-coil domain count and length (Fig. 1D). Only Slr7083 exhibited a somewhat characteristic domain architecture of eukaryotic IF proteins, whereas the coiled-coil domain distribution in the other candidates had major differences in coiled-coil domain number and lengths. None of the predicted CCRPs exhibited a stutter-like structure in the last coiled-coil segment. Besides coiled-coil domains, the COILS algorithm also predicted tetratricopeptide repeats (TPRs) as coiled-coils, thus we also included All4981 into our analysis, even though conserved domain searches reliably predicted these domains as TPRs and not coiled-coils. Many protein candidates contained conserved domains from eukaryotic IF proteins, found in Crescentin and FilP or from the bacterial cell division protein EzrA (Supplementary Table 1). The presence of these domains may be regarded as support for our classification. Additionally, structural maintenance of chromosomes (SMC) domains were predicted in almost all chosen candidates, all eukaryotic IF proteins as well as in Crescentin and FilP (Supplementary Table 1). The MscS_TM domain from Desmin was found in Slr7083 and Tlr0420 contains a Neuromodulin_N as well as a CCDC158 domain, both present in FilP or Crescentin, respectively.

The presence of homologs across all cyanobacterial morphotypes serves as a hint for universal protein function while a restricted distribution in specific subsections or morphotypes indicates a functional specialization within the respective taxon. An example for such species-specific candidate in our list is *slr7083* that is encoded on the pSYSA toxin-antitoxin plasmid in *Synechocystis*, similarly to *parM* and *tubZ*, which mediate plasmid segregation (Larsen *et al.*, 2007; Bharat *et al.*, 2015). In contrast, the homologous proteins Synpcc7942_1139 (abbreviated Syc1139) and Slr1301 are highly conserved and have homologs among all cyanobacterial groups (Fig. 1), including CypS from *Anabaena*, which we previously identified as a filament-forming CCRP (Springstein *et al.*, 2019). As our candidate CCRPs annotated as hypothetical proteins, we initially verified the transcription of the respective genes by RT-PCR from cDNA (Supplementary Fig. 1A-D). Our results showed that *slr7083* was only weakly transcribed during mid-exponential culture growth phase and *all4981* was found to be transcribed in an operon with its upstream genes *all4982* and *all4983* (Supplementary Fig. 1B,C).

### Cyanobacterial CCRPs assemble into diverse filamentous structures *in vitro*

A major characteristic of filament-forming proteins is their ability to self-polymerize into filaments intra and extracellularly (Fuchs and Weber, 1994; Köster *et al.*, 2015). Unlike actin and tubulin, IFs are able to form filamentous structures *in vitro* in a nucleotide-independent manner without additional co-factors upon renaturation from a denaturing buffer (Köster *et al.*, 2015). To examine the self-polymerization property of the nine tested CCRPs, we purified His_6_-tagged CCRPs under denaturing conditions and subjected them to subsequent renaturation by dialysis. Here we used protein concentrations in a similar range (0.5-1 mg ml^−1^) to previously investigated proteins shown to form filaments *in vitro* (e.g., Crescentin (Ausmees, Kuhn and Jacobs-Wagner, 2003) and Scc (England *et al.*, 2005), the metabolic enzyme CtpS (Ingerson-Mahar *et al.*, 2010) and the bactofilins BacA, BacB (Kühn *et al.*, 2010) and BacM (Koch, McHugh and Hoiczyk, 2011)). When applicable, the purified proteins were labeled with NHS-Fluorescein and the formation of *in vitro* filaments was assessed by epifluorescence or bright field microscopy. Several candidates did not form discernible structures *in vitro* and were consequently excluded from further investigation (including Slr6096, Tlr0420 and Fm6009; Supplementary Fig. 2A). The remaining CCRPs assembled into highly diverse structures *in vitro* (Fig. 2). Direct dialysis of Fm7001 from a high urea-containing buffer to a physiological buffer led to protein precipitation. However, upon slow stepwise renaturation (removing 0.5 M every 2 h), Fm7001 polymerized into a flat two-dimensional sheet floating on top of the dialysate in 4,5 M urea (Supplementary Fig. 2D). We addressed the eventuality that these structures could be the product of crystalized urea, but control experiments did not reveal filaments. Polymerized Fm7001 revealed two-dimensional filamentous sheets as well as single filamentous fibers (Fig. 2). Similar structures were observed for purified Fm7001-GFP and MBP-Fm7001-His_6_ (Supplementary Fig 2E,F). A two-dimensional filamentation pattern was observed also for Slr7083, which formed single, long and straight filamentous strings that were interconnected by two-dimensional sheets, thereby producing an irregular net (Fig. 2). Similarly, All4981 assembled into an interconnected filamentous net with thin single filaments (Fig. 2). The heterologous expression of Syc2039-His_6_ in *E. coli* failed, but we successfully purified Syc2039-GFP-His_6_ from *Synechococcus* instead. The polymerization pattern of Syc2039-GFP-His_6_ revealed sphere or cell shape-like three-dimensional sheets (Fig. 2). However, we note that most of the protein precipitated upon renaturation, hence it is unlikely that Syc2039 has *in vitro* polymerizing properties. Syc1139 polymerized into similar cell shape-like three-dimensional sheets but without any detectable aggregates (Fig. 2). The resemblance between Syc2039 and Syc1139 sheets raised the possibility that the sheet-like structures observed in the Syc2039-GFP-His_6_ sample represented co-precipitated and polymerized Syc1139. In accordance with this suggestion, we identified direct interactions of Syc1139 and Syc2039 using the bacterial adenylate cyclase two-hybrid (BACTH) assays (Supplementary Fig. 3A). For Slr1301, no clear *in vitro* structures were observed (Fig. 2). Nonetheless, we included this protein in further analyses since its homolog in *Anabaena* (CypS) has been recently reported as a filament-forming protein (Springstein *et al.*, 2019). Notably, Crescentin, which we used as a positive control, polymerized into smooth and filigree filaments only in the presence of monovalent ions (i.e. NaCl; Supplementary Fig. 2B). This observation highlights the importance of suitable buffer conditions for the detection of filament-forming proteins. To further confirm our *in vitro* observations, we included the monomeric and highly soluble maltose binding protein (MBP) as well as the oligomeric proteins GroEL1.2 (from *Chlorogloeopsis fritschii* PCC 6912; (Weissenbach *et al.*, 2017)) and the UMP kinase (from *Anabaena*) as negative controls. While both, the MBP and the UMP kinase readily clumped into comparably small aggregates, GroEL1.2 formed large proteinaceous aggregates *in vitro*, likely as a result of uncoordinated multimerization (Supplementary Fig. 2C). Consequently, we conclude that the *in vitro* filaments of the cyanobacterial CCRPs we observed here are unlikely to be oligomerization artifacts. We further validated the self-binding properties of the remaining six CCRPs using the BACTH assay and found that all proteins are able to self-interact (Supplementary Fig. 3A).

**Fig. 2:**
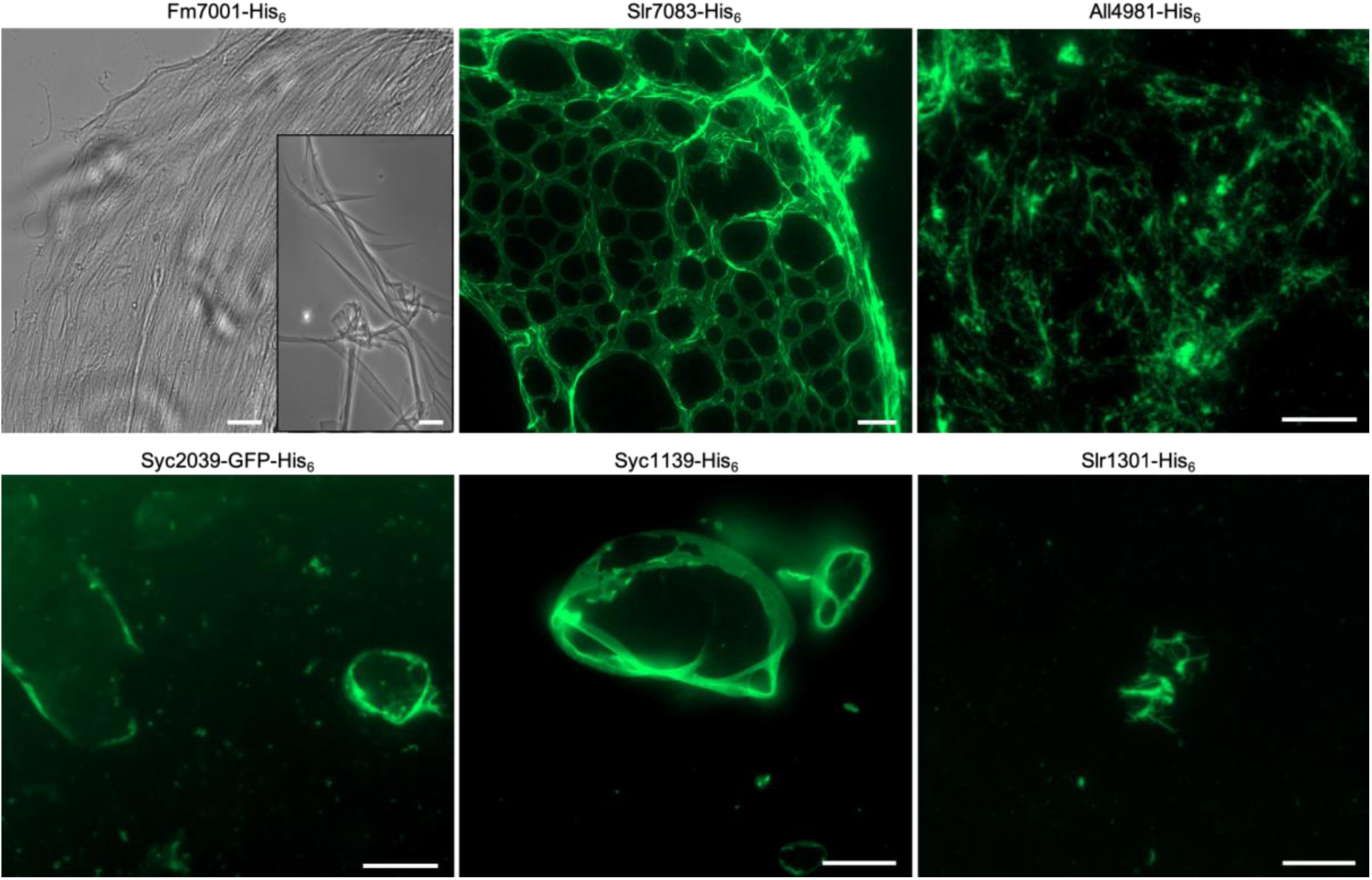
Cyanobacterial CCRPs assemble into diverse filamentous structures *in vitro*. Bright field and epifluorescence micrographs of filamentous structures formed by purified and renatured Fm7001-His_6_ (0.7 mg ml^−1^), Slr7083-His_6_ (1 mg ml^−1^), All4981-His_6_ (0.5 mg ml^−1^), Syc2039-GFP-His_6_ (0.3 mg ml^−1^), Syc1139-His_6_ (0.5 mg ml^−1^) and Slr1301-His_6_ (0.5 mg ml^−1^). Proteins were dialyzed into 2 mM Tris-HCl, 4.5 M urea pH 7.5 (Fm7001), HLB (Slr7083), PLB (All4981, Syc1139, Slr1301) or BG11 (Syc2039). Renatured proteins were either directly analyzed by bright field microscopy (Fm7001) or stained with an excess of NHS-Fluorescein and analyzed by epifluorescence microscopy. The NHS-Fluorescein dye binds primary amines and is thus incompatible with urea, which is why Fm7001 filaments were visualized by bright field microscopy. Scale bars: 10 µm or (Fm7001 inlay and Slr7083) 20 µm.

### Putative filament-forming proteins form filaments *in vivo*

To investigate whether the genetic background influences the filamentation properties of the candidate proteins, we expressed GFP or YFP translational fusion constructs of the putative filament-forming CCRPs in multiple hosts: 1) *E. coli*, 2) their native cyanobacterium and 3) in cyanobacteria of a different morphotype or subsection. Gene expression was driven by inducible or constitutive promoters commonly used in cyanobacteria. These included P_cpc560_ (for *Synechocystis*) (Zhou *et al.*, 2014), P_trc_ (for *E. coli*, *Synechocystis* and *Synechococcus*) (Huang *et al.*, 2010) or P_petE_ (for *Anabaena* and *Fischerella*) (Buikema and Haselkorn, 2001). As a positive control for *in vivo* filamentation, we expressed Crescentin-GFP in *Anabaena*, which formed round and helical filaments in the cells, thereby showing that P_petE_ is suitable for studying filament-forming IF-like CCRPs in *Anabaena*. (Supplementary Fig. 4A).

### Fm7001 forms protein filaments *in vivo* independent of the host

The *in vivo* localization of Fm7001 in *Fischerella* showed different results depending on the tag orientation. Only the expression of N-terminal YFP fusions of Fm7001 resulted in filamentous structures (Fig. 3 and Supplementary Fig. 4B). In *Synechocystis*, YFP-Fm7001 formed filaments throughout the cell (Fig. 3A) while in *Anabaena* we observed septum-arising filamentous strings (Fig. 3B). In its host, *Fischerella*, YFP-Fm7001 only rarely assembled into short filamentous strings (Fig. 3C inlays). Despite of the low abundance of filaments in *Fischerella*, induction of heterologous expression of YFP-Fm7001 induced an altered cell phenotype and trichomes seemingly divided in more than one plane resulting in a multiseriate (more than one trichome in a row) phenotype characteristic of *C. fritschii*. While under non-inducing conditions (i.e. in the absence of copper), *Fischerella* cells carrying a plasmid that expresses YFP-Fm7001 from P_petE_ had a WT phenotype, an altered morphotype and multiseriate growth was observed after around 4 rounds of replication (i.e. after 7 d) under inducing conditions (Fig. 3C). We also observed that, although expressed from a non-native promoter, YFP-Fm7001 was initially localized at branching points (Fig. 3C, 19 h after induction). Those effects suggest that Fm7001 may be involved in cell shape control and in the true-branching phenotype of *Fischerella*. Our attempts to generate a *Fischerella* Δ*fm7001* mutant strain remained unsuccessful, hence the function of Fm7001 remains unknown.

**Fig. 3:**
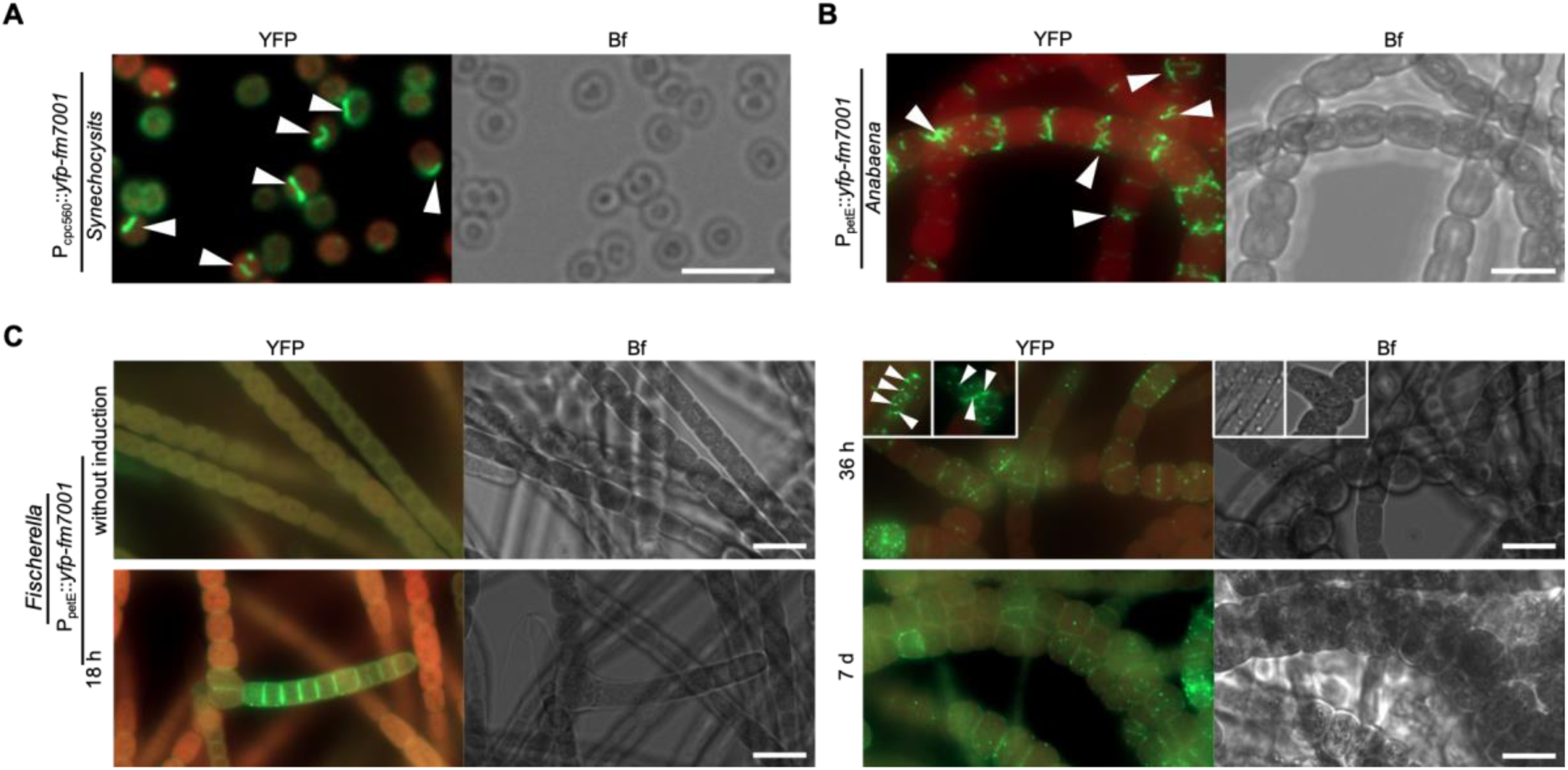
Host-independency for Fm7001 *in vivo* filamentation. Merged GFP fluorescence and chlorophyll autofluorescence (red) and bright field micrographs of (A) *Synechocystis*, **(A)** *Anabaena* or (C) *Fischerella* cells expressing YFP-Fm7001. Cells were either grown in (**A**,**B**) BG11 or (C)BG11 without copper and then induced with 0.5 µM CuSO_4_. (**C**) Micrographs were taken before induction of *yfp-fm7001* expression (without induction) and 18 h, 36 h or 7 d post induction. White triangles point to selected YFP-Fm7001 filamentous strings within the cells. Notably, unlike in *Anabaena* and *Fischerella*, Fm7001-GFP induced a swollen morphotype in *E. coli* and a subpopulation of *Synechocystis* cells (Supplementary Fig. 1E). (**B**): maximum intensity projection of a Z-stack. Scale bars: **(A**,**B**) 5 µm, (**C**) 10 µm.

### Slr7083 and Slr1301 are involved in twitching motility in *Synechocystis*

The *in vivo* localization of Slr7083-GFP in *Synechocystis* showed that it was localized to the cell periphery as well as rare focal spots and S-shaped filaments (Fig. 4A). We also attempted to localize Slr7083-GFP in the motile *Synechocystis* PCC-M substrain (hereafter PCC-M) but never obtained any successfully transformed clone, suggesting that overrepresentation of Slr7083 is deleterious for this strain. The localization of Slr1301-YFP in *Synechocystis* and PCC-M was at indistinct peripheral sites as assemblies of crescent-like shapes and rarely as S-shaped filaments (Fig. 4A and Supplementary Fig. 4C). Similar structures have been previously reported for the pilus ATPase PilB (Schuergers *et al.*, 2015). The localization of Slr7083-GFP and YFP-Slr7083 in *Anabaena* was at the cell periphery (Supplementary Fig. 4D). Furthermore, extended expression of YFP-Slr7083 in *Anabaena* altered the cellular morphology and disturbed the linear *Anabaena* trichome growth pattern (Supplementary Fig. 4E). In *E. coli*, Slr7083-GFP localized next to the cell poles (Supplementary Fig. 1E). When expressed in *E. coli*, Slr1301-GFP revealed a similar polar localization (Supplementary Fig. 1E). Additionally, reminiscent of Slr7083, the heterologous expression of Slr1301-YFP in *Anabaena* had an effect on the *Anabaena* cell-shape where it localized at the periphery and also formed single filaments or thick filamentous bundles that seemingly traversed through several cells (Supplementary Fig. 4C). To further assess the role of Slr1301 and Slr7083 in *Synechocystis* motility, we generated *Synechocystis* and PCC-M Δ*slr7083* and Δ*slr1301* mutant strains. The *Synechocystis* Δ*slr7083* and Δ*slr1301* mutants revealed no phenotypic defects compared to the WT (Fig. 4B,C). In contrast, the PCC-M Δ*slr7083* mutant is characterized by a decrease in twitching motility and a defect in cytokinesis (Fig. 4B). PCC-M Δ*slr7083* mutant cells often lacked internal chlorophyll signal entirely and failed to properly divide internal thylakoid membrane (assessed by the lack of chlorophyll autofluorescence) during cell division (Fig. 4B). Similarly, the PCC-M Δ*slr1301* mutant lost its twitching motility (Fig. 4B; confirming previous results from Bhaya, Takahashi, Shahi, & Arthur (2001)). Attempts to complement the motility defect in the PCC-M Δ*slr1301* mutant by expressing Slr1301-YFP from the conjugation plasmid pRL153 failed, possibly as a result of the comparably high expression of Slr1301-YFP from P_trc_ (we note that P_trc_ cannot be regulated by IPTG in *Synechocystis*). Additional attempts to complement the PCC-M Δ*slr7083* mutant never resulted in exconjugants. In order to further explore how Slr1301 affects motility, we analyzed co-precipitated proteins of Slr1301-YFP expressed in *Synechocystis* by mass spectrometry. This revealed multiple putative interaction partners involved in motility, including a twitching motility protein (Slr0161), two methyl-accepting chemotaxis proteins (McpA and PilJ) and the type IV pilus assembly ATPase PilB (Fig. 4D). The interaction of Slr1301 with PilB, together with their similar *in vivo* localization, prompted us to characterize the interaction of both proteins. For this purpose, we attempted to express PilB-GFP in *Synechocystis* WT, and in the Δ*slr1301* and Δ*slr7083* mutants. In *Synechocystis* WT, PilB-GFP localized to the cell periphery and often formed crescent-like formations (reminiscent of Slr1301-YFP and Slr7083-GFP; Fig. 4A), confirming previous results by Schuergers *et al.* (2015). However, we never observed any PilB-GFP expression in the *Synechocystis* or PCC-M Δ*slr7083* and Δ*slr1301* mutants. The similarity between our observations so far for Slr1301 and Slr7083 led us to test for an interaction between these two proteins. Indeed, a bacterial two-hybrid assay confirmed a direct interaction between Slr7083 and Slr1301 (Fig. 4E). Taken together, our investigation identified two *Synechocystis* CCRPs that are involved in cell motility. Slr7083 is a cell envelope-localized protein involved in cytokinesis and motility. It polymerized into filaments *in vitro* but only few filaments were identified *in vivo*. Slr1301, although failing to assemble into filaments *in vitro*, occasionally polymerized into filaments *in vivo* and was found to be an interaction partner of proteins that function in *Synechocystis* twitching motility.

**Fig. 4:**
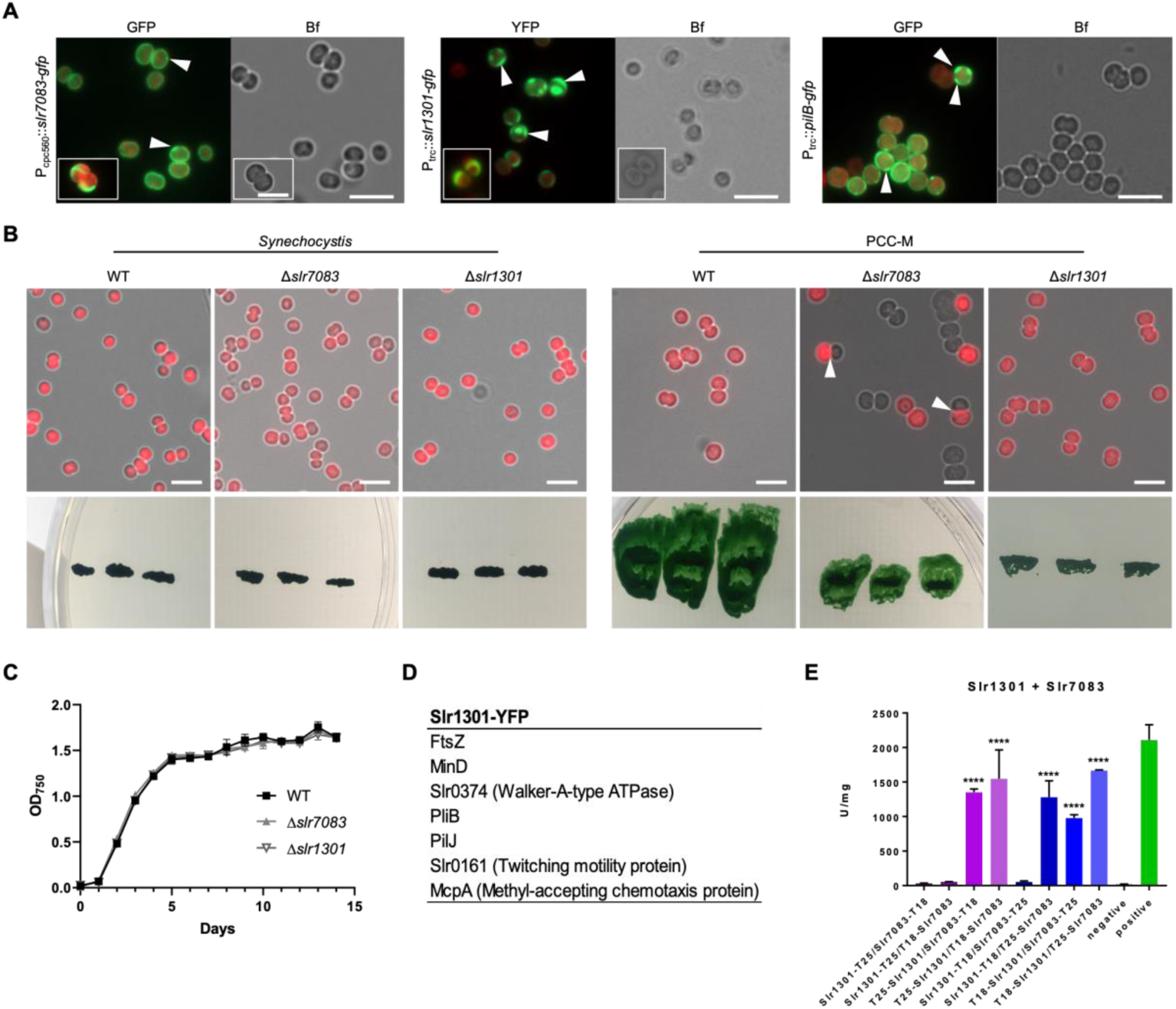
Slr7083 and Slr1301 are involved in twitching motility in *Synechocystis*. (**A**) Merged GFP fluorescence and chlorophyll autofluorescence (red) and bright field micrographs of *Synechocystis* cells expressing, Slr7083-GFP, Slr1301-YFP or PilB-GFP from P_cpc560_ (Slr7083) or P_trc_ (Slr1301, PilB). Expression of PilB-GFP in PCC-M resulted in the same localization pattern (data not shown). White triangles indicate focal spots and crescent-like formations. Scale bars: 5 μm. (**B**) Merged bright field and chlorophyll autofluorescence micrographs of motile and non-motile *Synechocystis* WT, Δ*slr7083* and Δ*slr1301* mutant cells. Below, motility tests of three single colonies from indicated cells streaked on BG11 plates and illuminated from only one direction are shown. (**C**) Growth curve of *Synechocystis* WT, Δ*slr7083* and Δ*slr1301* mutant strains grown in quadruples at standard growth conditions. OD_750_ values were recorded once a day for 15 d. Error bars show the standard deviation (n=4). (**D**) Excerpt of interacting proteins of interest from mass spectrometry analysis of anti-GFP co-immunoprecipitations of *Synechocystis* cells expressing Slr1301-YFP from P_trc_. (**E**) Beta-galactosidase assays of *E. coli* cells co-expressing indicated translational fusion constructs of all possible pair-wise combinations of Slr7083 with Slr1301 grown for 1 d at 30 °C. Quantity values are given in Miller Units per milligram LacZ of the mean results from three independent colonies. Error bars indicate standard deviations (n=3). Neg: pKNT25 plasmid carrying *slr1301* co-transformed with empty pUT18C. Pos: Zip/Zip control. Values indicated with * are significantly different from the negative control. *: P<0.05, **: P<0.01, ***: P<0.001, ****: P<0.0001 (Dunnett’s multiple comparison test and one-way ANOVA).

### All4981 is an *Anabaena* TPR protein that forms septal-arising filaments

The expression of All4981-GFP in *Anabaena* revealed numerous filaments that traversed the cell while in other cells All4981-GFP was associated with the cell septa (Fig. 5A). All4981-GFP filaments also occasionally spread in a star-like pattern into the cytosol. Additionally, in freshly ruptured All4981-GFP-expressing cells, filamentous *ex vivo* structures assembled in the medium into an interconnected network (Supplementary Fig. 4F), resembling the *in vitro* polymerization pattern of All4981 (Fig 2). We confirmed a host-independent *in vivo* polymerization capacity of All4981 by expressing All4981-GFP in *Synechocystis*, which lacks homologs to that protein (Fig. 5B). Intrigued by the septal localization, we tested for an interaction with SepJ, a septal junction protein in *Anabaena* (Flores *et al.*, 2007) and found weak, albeit significant physical interactions (Supplementary Fig. 3A). In addition, bacterial two-hybrid assays revealed that All4981 interacted with two other *Anabaena* filament-forming CCRPs, namely LfiA and LfiB (Springstein *et al.*, 2019), and strongly interacted with the cell shape-determining protein MreB (Supplementary Fig. 3A). Notably, MreB has previously been shown to form similar filamentous structures in *Anabaena*. However, in contrast to genes in the *mreBCD* operon, whose overexpression induces abnormal cell morphologies (Hu *et al.*, 2007), no direct morphogenic influence was detected for All4981 in *Anabaena*. Notably, it is likely that All4981 is an essential protein in *Anabaena* as we were not able to generate an *all4981* deletion strain. Initially, we accidently also created a YFP-All4981 fusion construct with a deletion of 240 bp between nt 735 and nt 975 of the *all4981* CDS, resulting in a deletion of the third and fourth TPR (YFP-All4981^ΔTPR3-4^) leaving the remaining ORF intact. Remarkably, this fusion protein, like All4981-GFP, formed cell-traversing filaments in *Anabaena* and sometimes assembled into a filamentous structure within the cells (Supplementary Fig. 4G). In contrast, full length YFP-All4981 localized to the septa between two neighboring cells but also revealed indistinct cytosolic localization (Supplementary Fig. 4G). Co-immunoprecipitation experiments following LC-MS/MS analytics from *Anabaena* WT expressing YFP-All4981^ΔTPR3-4^ revealed an association of YFP-All4981^ΔTPR3-4^ with ParB, MinD and MreB (Fig. 5C). Thus, All4981 might be involved in ParA/B/S-driven plasmid or chromosome segregation. The interaction with MreB agrees with the *in vivo* localization of YFP-All4981^ΔTPR3-4^ in *Anabaena* (Supplementary Fig. 4G) and the results from the bacterial two-hybrid assay (Supplementary Fig. 3A). Further interactions were found with a variety of putative S-layer and prohibitin-like proteins and with DevH, an essential protein for heterocyst glycolipid layer synthesis. Notably, we never observed All4981 expression in heterocysts, regardless of the fluorescence tag. All4981 also interacted with All4982, a protein encoded directly upstream of *all4981*, but not with All4983, which is encoded upstream of *all4982* (Supplementary Fig. 1C). This observation, together with the common transcript of *all4981* and *all4982* (Supplementary Fig. 1C) argues for a common function of both proteins. Thus, we attempted to localize All4982 with an eCFP tag in *Anabaena* but could not observe a coherent localization pattern. Overall, our results demonstrate that All4981 is connected to other *Anabaena* filament-forming CCRPs, the MreB cytoskeleton, the septal junctions and the protective S-layer. Additionally, All4981 polymerizes *in vitro*, *in vivo* and *ex vivo*, is likely essential for *Anabaena* and is thus accordingly classified as a novel cyanobacterial filament-forming TPR-repeat protein.

**Fig. 5:**
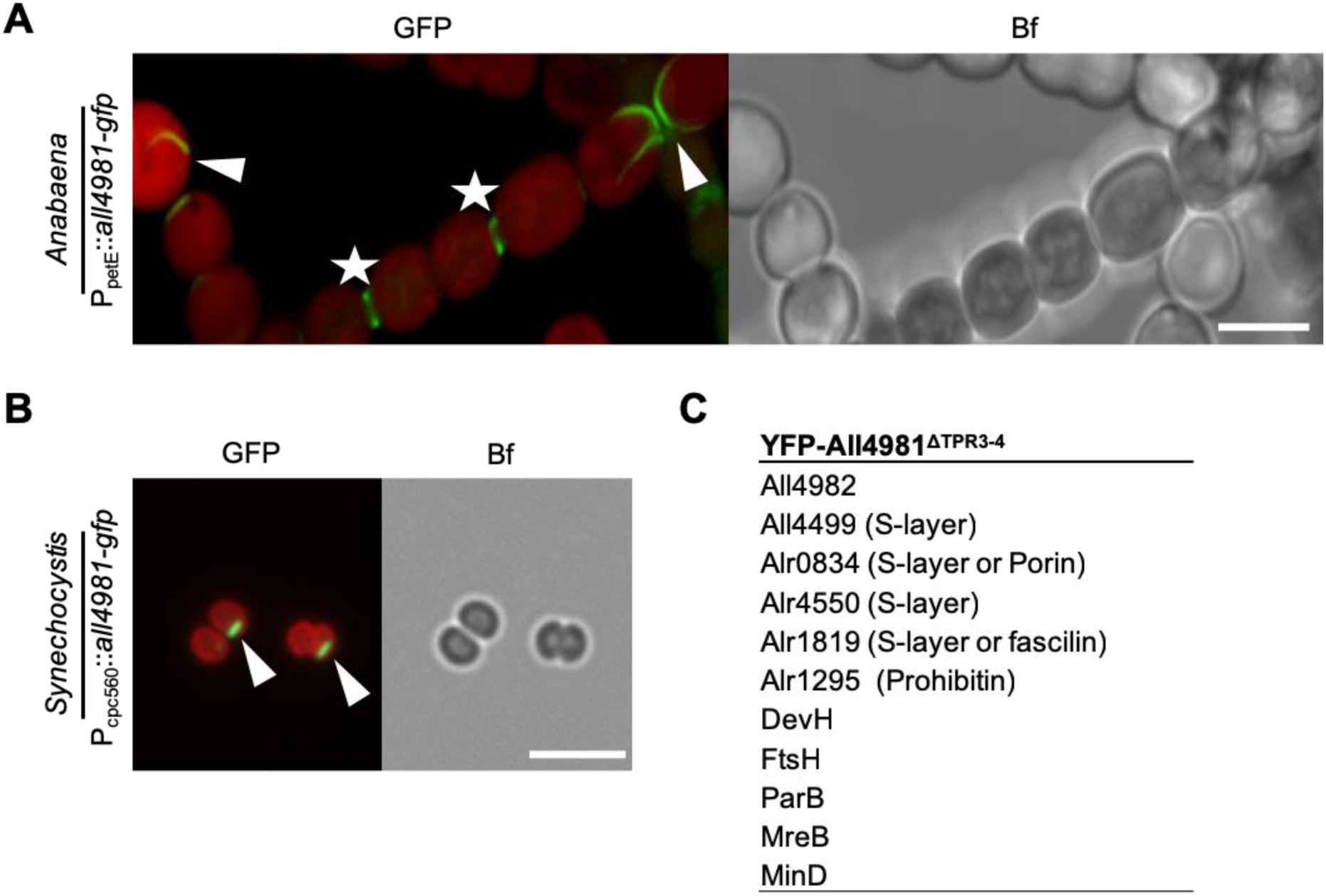
All4981 forms cell-traversing filaments in cyanobacteria. (**A,B**) GFP fluorescence and merged GFP fluorescence and chlorophyll autofluorescence (red) and bright field micrographs of (**A**) *Anabaena* and (**B**) *Synechocystis* cells expressing All4981-GFP. *Anabaena* cells were grown in BG11_0_ and *Synechocystis* cells were grown in BG11. (**A**): Maximum intensity projections of a Z-stack. White triangles indicate selected filaments traversing through the cells. White arrows point to spindle-like YFP-All4981 filaments. White stars mark septal formations between two neighboring cells. Scale bars: 5 µm. (**C**) Excerpt of interacting proteins of interest from mass spectrometry analysis of anti-GFP co-immunoprecipitations of *Anabaena* cells expressing YFP-All4981^ΔTPR3-4^ from P_petE_.

### *Synechococcus* CCRPs are involved in cytokinesis and colony integrity

The results of the *in vivo* localization of a functional Syc2039-GFP fusion protein (Supplementary Fig. 5E,F) contrasted the ambiguous *in vitro* polymerization pattern (Fig. 2). Filaments were readily observed in different cyanobacterial hosts, indicating that in Syc2039 self-polymerization is independent of the host (Fig. 6A). Notably, however, Syc2039 formed different structures in each host. In *Anabaena*, filaments were long, curved and intertwined; in *Synechocystis* filaments appeared as spindle-like structures and in *Synechococcus* filaments were long, sometimes helical and often aligned with or in close proximity to the cell envelope (Fig. 6A). A similar helical or cell periphery-aligned localization pattern was also observed in *E. coli* (Supplementary Fig. 1E). In *Synechocystis* and *Synechococcus* Syc1139-GFP localized as spots at the cell periphery, while in *E. coli* it seemingly coated the entire cell envelope (Fig. 6A, Supplementary Fig. 1E). Syc1139-GFP failed to be expressed in *Anabaena*, suggesting that (over-)expression of this protein has a negative impact on that organism. Using double homologous gene replacement, we generated a *Synechococcus* Δ*syc2039* mutant strain and a non-segregated *Synechococcus* Δ*syc1139* mutant strain (Supplementary Fig. 5A-C). The non-segregated nature of the Δ*syc1139* mutant suggests that this gene performs an essential cellular function and cannot be fully deleted. Colony integrity of the Δ*syc2039* mutant was unaltered while the Δ*syc1139* mutant was characterized by apparent changes in colony morphology (Fig. 6B), which were lost upon growth on non-selective plates (Supplementary Fig. 5D). Additionally, both mutants presented an impairment in liquid culture growth: the Δ*syc2039* mutant grew in standard BG11 medium but failed to grow upon addition of several osmotic stressors, whereas the Δ*syc1139* mutant failed to grow in liquid culture entirely (Fig. 6C). Spot assays confirmed a decreased viability of the Δ*syc1139* mutant and showed that it is highly sensitive to Proteinase K but unaffected by lysozyme (Supplementary Fig. 6A). These cell wall defects, together with the *in vitro* cell shape-like filamentation pattern suggest that Syc1139 might form a protective and protease-resistant proteinaceous layer below the cytoplasmic membrane. This possibility would also be in concert with the distorted colony morphology of the non-segregated Δ*syc1139* mutant strain. The Δ*syc2039* mutant was unaffected by cell wall and membrane destabilizers (Supplementary Fig. 6B). To investigate the role of these proteins in cell division, we stained intracellular DNA with DAPI and localization of FtsZ was detected by immunofluorescence in *Synechococcus* WT and both mutant strains. A proportion of Δ*syc2039* mutant cells exhibited a segregated DNA distribution either to both cell poles or to just one pole (Fig. 6D). Furthermore, some cells of both mutants lacked any discernible intracellular DNA or perceptible chlorophyll signal and were elongated compared to the WT (Fig. 6D,E). The WT phenotype of the Δ*syc2039* mutant could be rescued by insertion of P_trc_::*syc2039-gfp* or P_syc2039_::*syc2039* into the neutral NS1 (Bustos and Golden, 1992) locus (Supplementary Fig. 5E,F). Although both mutant cells were elongated compared to WT cells (Fig. 6E), the intracellular localization of FtsZ was unaffected (Supplementary Fig. 6C). And despite the defect in cytokinesis, the Δ*syc2039* mutant strain revealed similar liquid culture growth properties as the WT (Supplementary Fig. 6D). Taken together, Syc2039 forms abundant filamentous networks *in vivo* and is involved in cytokinesis or cell cycle control. We could further show that *syc1139* is an essential gene important for cytokinesis, cellular integrity and colony formation, implicating structural functions.

**Fig. 6:**
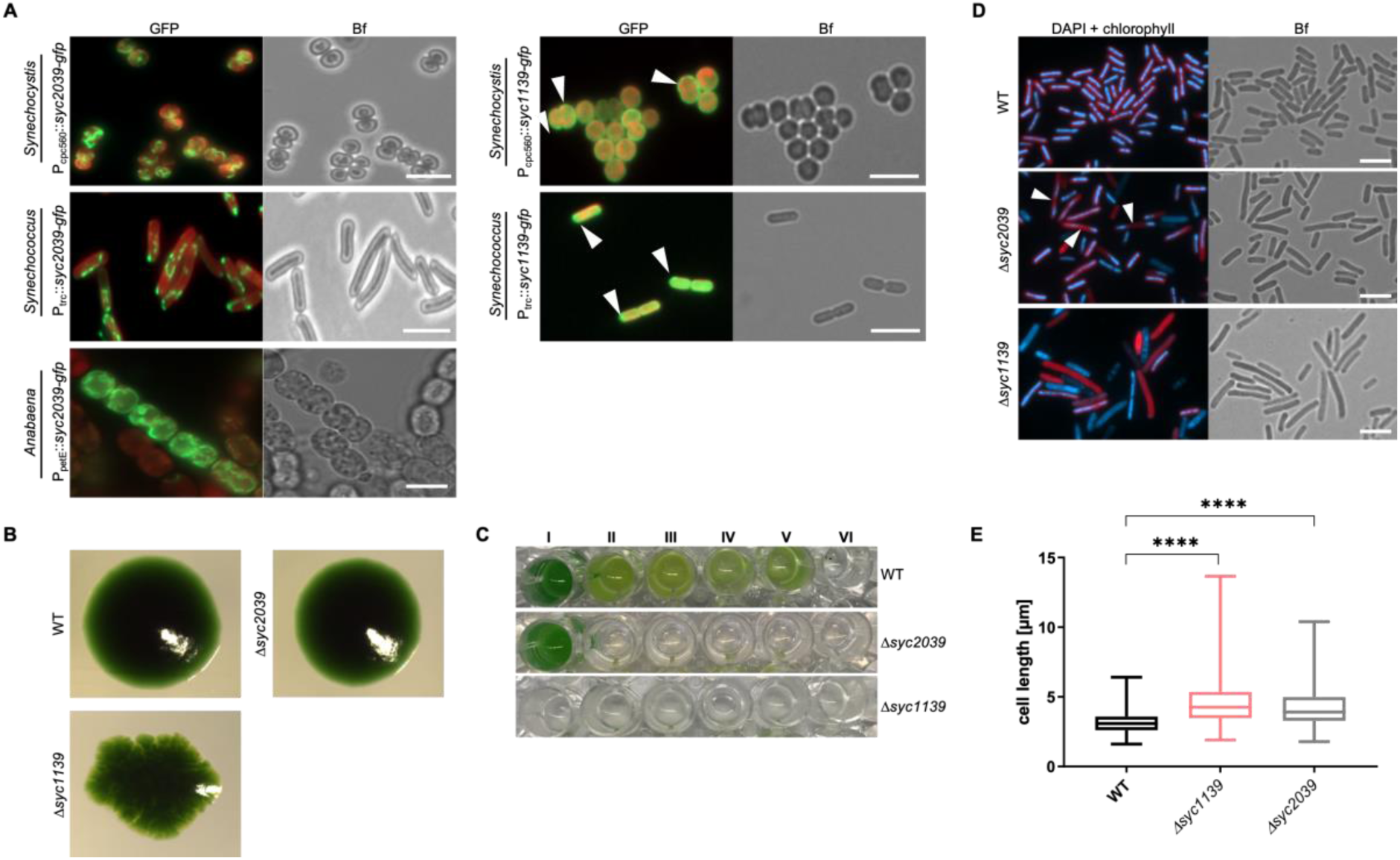
*Synechococcus* CCRPs affect cytokinesis and cellular integrity. (**A**) Merged GFP fluorescence and chlorophyll autofluorescence (red) and bright field micrographs of *Synechocystis*, *Synechococcus* and *Anabaena* cells expressing Syc2039-GFP or Syc1139-GFP from P_trc_. *Synechocystis* cells were grown in BG11, *Anabaena* cells were grown in BG11_0_ supplemented with 0.25 µM CuSO_4_ for 1 day, and *Synechococcus* cells were grown on BG11 plates supplemented with 0.01 mM (Syc2039) or 1 mM (Syc1139) IPTG. Micrographs of *Synechococcus* and *Anabaena* cells expressing Syc2039-GFP are maximum intensity projections of a Z-stack. White triangles indicate Syc1139-GFP spots. Attempts to translationally fuse a YFP-tag to the N-terminus of Syc2039 were unsuccessful, possibly due to the transmembrane domain predicted to the Syc2039 N-terminus (Supplementary Table 1). (**B**) Colony formation of *Synechococcus* WT and mutant strains on BG11 plates. (**C**) Cell viability of *Synechococcus* WT and mutant strains grown in (I) BG11 or BG11 supplemented with (II) 5 mM glucose, (III) 200 mM glucose, (IV) 2 mM NH4Cl, (V) 200 mM maltose or (VI) 500 mM NaCl. (**D**) Merged DAPI fluorescence and chlorophyll autofluorescence (red) and bright field micrographs of *Synechococcus* WT and mutant strains grown on BG11 plates and stained with 10 μg ml^−1^ DAPI. White triangles indicate non-dividing cells revealing inhomogeneous DNA placement. (**E**) Cell length of *Synechococcus* WT (n=648), non-segregated Δ*syc1139* (n=417) and Δ*syc2039* (n=711) mutant cells. Values indicated with * are significantly different from the WT. ****: P<0.0001 (one-way ANOVA, using Turkey’s multiple comparison test). Scale bars: 5 µm.

## Discussion

Earlier studies suggested that there is likely a broad spectrum of coiled-coil-rich and rod-domain containing proteins with IF-like function in prokaryotes (Bagchi *et al.*, 2008). And indeed, reports on such proteins followed with the discovery of Scy (in *Streptomyces coelicolor*) and several CCRPs from *Helicobacter pylori* (Waidner *et al.*, 2009; Walshaw, Gillespie and Kelemen, 2010; Specht *et al.*, 2011; Holmes *et al.*, 2013). Here we further investigated the presence and function of CCRPs with filament-forming IF-like properties in prokaryotes, by predicting and evaluating CCPRs in cyanobacteria. Our *in vitro* polymerization assay allowed for a rapid detection of protein filaments *in vitro* using fluorescence microscopy thus bypassing the need to investigate filament-formation by laborious electron microscopy procedures. The observed protein filament lengths were in the range of previously described *in vitro* filaments of FtsZ (Camberg, Hoskins and Wickner, 2009) and of the human prion protein in its amyloid form (Bocharova *et al.*, 2005) that were obtained by a similar experimental procedure.

Our results show that Fm7001 assembles into polymers *in vitro* upon renaturation from urea as well as *in vivo*, and that this protein has an impact on cellular and trichome morphology, thereby fulfilling major IF criteria (Köster *et al.*, 2015; Kelemen, 2017). Consequently, we propose that Fm7001 constitutes a novel filament-forming CCRP specific to multicellular, cell-differentiating and branching cyanobacteria. The floating Fm7001 polymer sheet in high molar urea (i.e. 4.5 M urea) indicates an exceptionally high self-association capacity of Fm7001. In comparison, the eukaryotic IF protein Vimentin exists only as tetramers in 5 M urea (Herrmann *et al.*, 1996). *In vivo* localization experiments revealed an essential role of the Fm7001 C-terminus for filamentation, which is a common observation for known prokaryotic filament-forming proteins, including MreB (Swulius and Jensen, 2012), Crescentin (Ausmees, Kuhn and Jacobs-Wagner, 2003) as well as eukaryotic IF proteins (Geisler and Weber, 1982; Weber and Geisler, 1982; Traub and Vorgias, 1983; Nakamura *et al.*, 1993; Herrmann *et al.*, 1996). Additionally, the assigned structural similarities of Fm7001 with the acetyl-CoA-carboxylase may provide further support for the theory that filament-forming proteins originated from metabolic enzymes that obtained polymerization features (Ingerson-Mahar and Gitai, 2012). Notwithstanding, the metabolic activity of Fm7001 was not evaluated in our study hence its presumed enzymatic activity remains to be tested. Additionally, so far, no sufficient genome modification systems exist for *Fischerella* (Stucken *et al.*, 2012; Stucken, Koch and Dagan, 2013), as such a precise analysis of the function of Fm7001 is currently not possible.

Several prokaryotic tubulin-like and actin-like cytoskeletal proteins, such as ParM and TubZ, are known to be encoded on plasmids or on bacteriophages (Hurme *et al.*, 1994; Wagstaff and Löwe, 2018). In *Synechocystis*, *slr7083* is encoded on the large toxin-antitoxin defense plasmid (pSYSA) (Kopfmann and Hess, 2013), thus it adds another protein to the list of those filament-forming CCRPs carried by an autonomously replicating genetic element. Preliminarily we suspected that Slr7083 has a role in plasmid-segregation similar to ParM. However, Slr7083 showed no indications of dynamic properties, which would be indispensable for a plasmid segregation mechanism. Furthermore, unlike ParM (Carballido-Lopez, 2006), Slr7083 did not localize in a spindle-like pattern *in vivo* and was only expressed at later growth phases, which is contradictory to a possible involvement in the cell cycle. In contrast, the polymers formed by Slr7083 *in vitro* and *in vivo* rather suggest that it could form a proteinaceous layer below the cytoplasmic membrane. Notably, Slr7083 *in vitro* structures resemble the nuclear lamina formed by nuclear lamins and FilP lace-like *in vitro* filaments (Stuurman, Heins and Aebi, 1998; Bagchi *et al.*, 2008; Fuchino *et al.*, 2013). It is thus conceivable that Slr7083 has a role in cellular stiffness as well as rigidity and mediates mechanical cell stabilization. However, restriction of transcription to only a comparably short period of the culture growth phase challenges the idea of a cell-stabilizing function for Slr7083. In contrast, cell motility in *Synechocystis* seems to be partially regulated by Slr7083, reminiscent of the role of the actin cytoskeleton in eukaryotes.

The role of Slr7083 in cell motility is possibly mediated by means of its interaction with Slr1301, which has already previously been shown to be essential for twitching motility in *Synechocystis* (Bhaya *et al.*, 2001). So far it is unknown how photoreceptors transduce the perceived light stimuli to the motility apparatus in *Synechocystis* ultimately resulting in phototactic movements (Schuergers, Mullineaux and Wilde, 2017). It is tenable to hypothesize that Slr1301 might constitute the missing link between the two systems, possibly in combination with Slr7083. This hypothesis is supported by the physical interaction of Slr1301 with PilB and the *in vivo* localization of Slr1301 that is similar to that observed for PilB (Schuergers *et al.*, 2015). A comparable complex was observed in *Pseudomonas aeruginosa*, where FimL (a proposed scaffolding protein with a weakly predicted coiled-coil) was shown to connect the chemosensory receptor system to the type IV pili apparatus, regulating the chemotactic and virulence pathways (Inclan *et al.*, 2016). In eukaryotes, cellular motility is strongly dependent on cytoskeletal proteins (Cappuccinelli, 1980), thus it is possible that filament-forming proteins are also key factors for cell locomotion in prokaryotes. Although IFs do not directly participate in cell motility in eukaryotes (Lodish *et al.*, 2000), an adaptation of filament-forming CCRPs in prokaryotes for this task is conceivable. Bactofilins constitute a separate class of prokaryotic-specific polymerizing proteins and were proposed to be involved in coordinated motility in *C. crescentus* (Kühn *et al.*, 2010). Additionally, the filament-forming CCRP AglZ from *Myxococcus xanthus* was previously shown to govern gliding motility together with a multi-protein complex that also involves the MreB cytoskeleton (Yang *et al.*, 2004; Nan *et al.*, 2010). The interaction of Slr1301 with twitching motility proteins was prevailed in the non-motile *Synechocystis* strain, hinting for additional beneficial functions of this interaction besides motility. Notably, we previously reported filament-forming properties for Alr0931 (CypS), which is a homolog of Slr1301 in *Anabaena* (Springstein *et al.*, 2019). While CypS polymerizes into filaments *in vitro*, Slr1301 does not, which could indicate a specific adaptation of CypS to filament-formation in multicellular cyanobacteria. Despite their different cellular functions and *in vitro* polymerization properties, the homologous proteins Slr1301, Syc1139 and CypS retained the ability to cross-interact (Supplementary Fig. 3A). Further studies may focus on identifying the protein domains that mediate this interaction, likely residing within the highly conserved amino acid sequence region in this homologous protein family (Supplementary Fig. 3B). These regions are likely important for an interaction with species-specific proteins that lead to their species-specific cellular function.

TPR proteins are known to mediate protein-protein interactions and can assemble into multimers, but their ability to polymerize into filaments has not been described so far (Blatch and Lässle, 1999). Nonetheless, All4981 polymerizes *in vitro* and *in vivo* in all tested hosts. Additionally, it forms extracellular filaments and is presumably an essential protein in *Anabaena*. These observations suggest that All4981 is a *bona fide* prokaryotic filament-forming TPR protein. The association of All4981 with MreB, FtsZ-regulators, the S-layer and SepJ indicates that it might function as a bridge that connects the shape-determinants outside of the cell wall and inside of the cytoplasmic membrane to the sites of cell-cell connections (i.e. septal junctions). A function of All4981 in *Anabaena* cell and filament shape is also supported by its interaction with the *Anabaena* filament and cell shape stabilizing proteins LfiA and LfiB (Springstein *et al.*, 2019).

Considering the presence of an N-terminal transmembrane domain and the lack of clear *in vitro* filaments, it is unlikely that Syc2039 constitutes a genuine filament-forming protein. Nonetheless, the highly abundant filamentous network formed in all tested bacterial hosts suggests that Syc2039 is associated with cytoskeletal structures. Specifically, the elongated phenotype and the disturbed cytokinesis in the *Synechococcus* Δ*syc2039* mutant and the non-segregated Δ*syc1139* mutant suggest an association with the FtsZ-driven elongasome. Direct interaction with FtsZ or MreB could not be shown, as such, future studies will likely attempt to unravel the presumed connection of the *Synechococcus* CCRPs to those two major cytoskeletal systems. Notably, besides its cytokinetic defect, the Δ*syc2039* mutant showed growth characteristics like the WT, suggesting that feedback mechanisms between cytokinesis and cell division are disturbed in the Δ*syc2039* mutant.

Our results reveal two novel filament-forming CCRPs - Fm7001 and All4981 – from different cyanobacterial subsections and morphotypes (Fig. 7). Our study thus extends the spectrum of known filament-forming CCRPs in prokaryotes and expands the set of functional properties associated with IF-like proteins in prokaryotes. Notably, as indicated by Bagchi *et al.* (2008), we demonstrate that the sole observation of coiled-coil-rich regions within a protein sequence cannot be regarded as a sole predictor of protein polymerization, hence identification of novel filament-forming proteins requires additional *in vitro* and *in vivo* assays. The cyanobacterial CCRPs we report here, like other bacterial CCRPs (Ausmees, Kuhn and Jacobs-Wagner, 2003; Bagchi *et al.*, 2008; Waidner *et al.*, 2009; Fiuza *et al.*, 2010; Specht *et al.*, 2011; Holmes *et al.*, 2013) and eukaryotic IFs (Alberts *et al.*, 2014) are essential cellular components (All4981), are important for cell shape determination (Fm7001, Syc1139 and Syc2039), mediate cellular motility (Slr7083 and Slr1301), DNA segregation (Syc1139 and Syc2039) and colony integrity (Syc1139). Our study thus strengthens the perception that like eukaryotes, prokaryotes require organized internal complexes and even microcompartments to maintain cell shape, size and proper cell function and highlights the usefulness of polymerized proteinaceous structures for cellular processes. Remarkably, some of the identified CCRPs were highly conserved among all cyanobacterial morphotypes, suggesting that their function is conserved. Future studies are required in order to evaluate the functional conservation of homologous proteins in different cyanobacterial species and morphotypes. On the other hand, Syc2039 and Slr7083 are highly strain specific, possibly performing a function that is adapted to the very needs of their hosts. Similarly to the eukaryotic cytolinker proteins (Leung, Green and Liem, 2002; Wiche, Osmanagic-Myers and Castañón, 2015), cyanobacterial CCRPs were often associated with other cytoskeletal systems (MreB, FtsZ and other filament-forming CCRPs) and sites of cell-cell connections (i.e. SepJ), which demonstrates the necessity for those structures to be in a constant interplay even in comparably small cells. The discovery of filament-forming CCRPs with different levels of conservation in various cyanobacterial morphotypes thus opens up new avenues of research on their contribution to cyanobacterial morphological diversity.

**Fig. 7:**
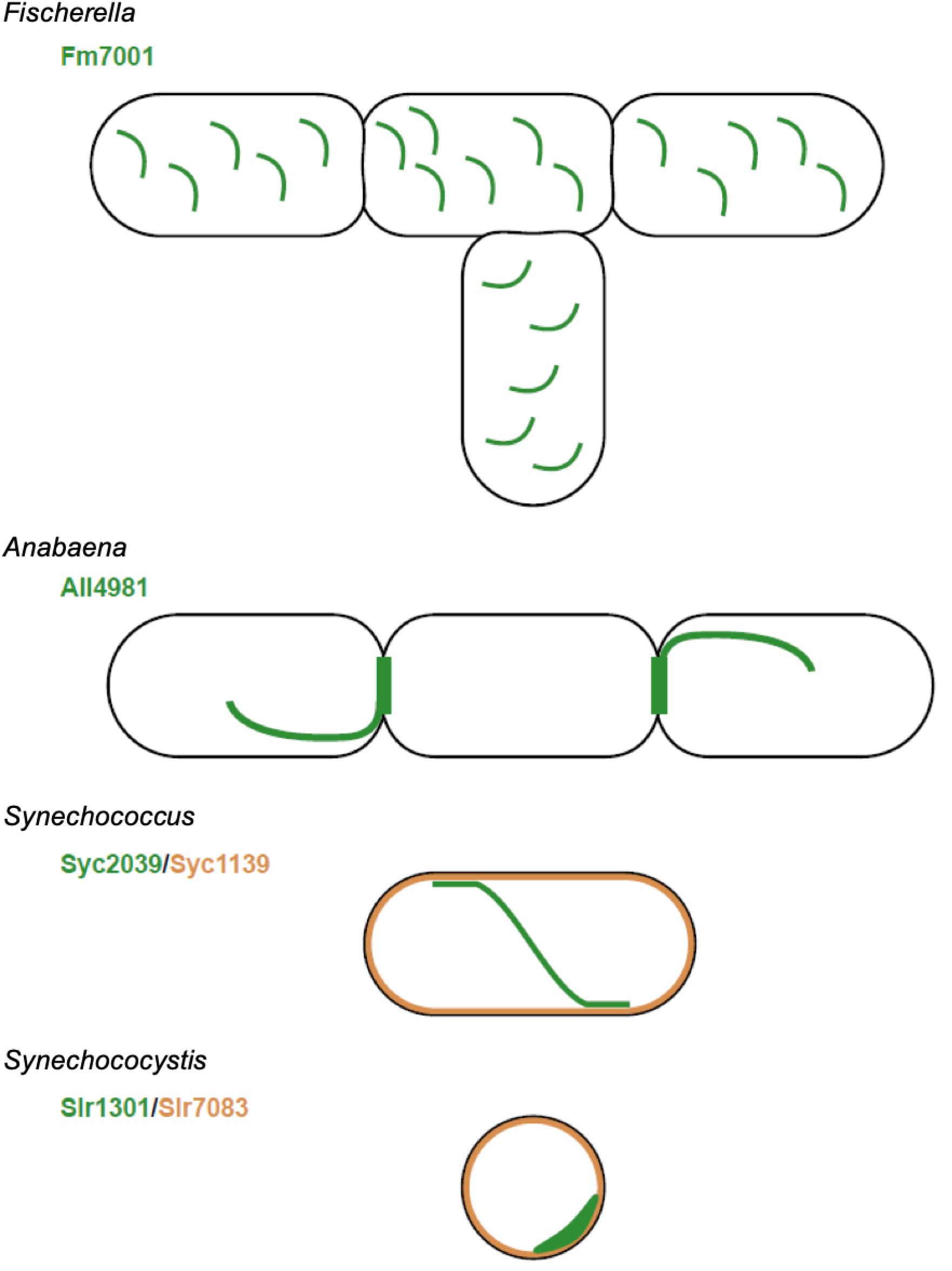
Cyanobacterial CCRP systems. Schematic models for the *in vivo* localization of cyanobacterial CCRPs in their respective hosts. Fm7001 forms filamentous strings in *Fischerella*. In *Anabaena*, All4981 assembles into pole-arising filaments that traverse through the cell or forms septal-localized bridge-like formations. Syc2039, either independently of other *Synechococcus* proteins, or in direct cooperation with other filamentous proteins, forms long and sometimes helical strings that are often aligned with or in close proximity to the cell periphery. In *Synechococcus*, Syc1139 likely forms a protective proteinaceous layer below the cytoplasmic membrane. In *Synechocystis*, Slr1301 forms crescent-like structures while Slr7083 seemingly underlies the cytoplasmic membrane. Both localization types were also observed for PilB, suggesting a cooperative function.

## Material and Methods

### Data and CCRP prediction

The cyanobacteria protein families were constructed from completely sequenced genomes available in RefSeq database (O’Leary *et al.*, 2015) (ver. May 2016; Supplementary File 2). For the construction of protein families, at the first stage, all protein sequences annotated in the genomes were blasted all-against-all using stand-alone BLAST (Altschul *et al.*, 1990) (V. 2.2.26). Protein sequence pairs that were found as reciprocal best BLAST hits (rBBHs; Tatusov, Koonin and Lipman, 1997) with a threshold of E-value ≤ 1×10^−5^ were further compared by global alignment using needle (EMBOSS package, V. 6.6.0.0; (Rice, Longden and Bleasby, 2000). Sequence pairs having ≥30% identical amino acids were clustered into protein families using the Markov clustering algorithm (MCL) (Enright, Van Dongen and Ouzounis, 2002) (ver. 12-135) with the default parameters. For the CCRPs prediction, 1,535 protein sequences containing non-standard amino acids were discarded. Coiled-coil regions in protein sequences were predicted using PEPCOIL (EMBOSS package, V. 6.6.0.0; (Rice, Longden and Bleasby, 2000). The algorithm was executed with a window size of 21 and the threshold for amino acids in coiled-coil conformation was set to ≥ 80 amino acid residues similarly as described by Bagchi *et al.* (2008). Statistical tests were performed with MatLab©. For the comparison of CCRPs proportion, the compared groups included: 1) SynProCya group, 2) unicellular cyanobacteria, 3) unicellular cyanobacteria that divide in more than one plane, and 4) multicellular cyanobacteria. Identification of conserved amino acid domains within cyanobacterial CCRP homologs (CypS (Alr0931), Slr1301 and Syc1139) was done using MULTALIGN (Corpet, 1988).

Protein candidates were further manually examined with online available bioinformatic tools (NCBI Conserved Domain (CD) Search (Marchler-Bauer *et al.*, 2016), TMHMM Server (Krogh *et al.*, 2001) (V. 2.0), PSIPRED (McGuffin, Bryson and Jones, 2000), PSORTb (Yu *et al.*, 2010) (ver. 3.0), I-TASSER (Zhang, 2009). CCRPs exhibiting similar predictions to known IF and IF-like proteins like CreS, FilP, vimentin, desmin or keratin were selected, and proteins predicted to be involved in other cellular processes were excluded.

### Bacterial strains and growth conditions

*Fischerella*, *Anabaena* and *Synechocystis* were obtained from the Pasteur Culture Collection (PCC) of cyanobacteria (France). *Synechococcus* was a gift from Martin Hagemann (University Rostock). Glucose-tolerant motile *Synechocystis* PCC-M substrain was a gift from Annegret Wilde (University Freiburg). Cells were grown photoautotropically in BG11 or without combined nitrogen (BG11_0_) at a 16h/8h light/dark regime (*Fischerella*) or at constant light (*Anabaena*, *Synechococcus* and *Synechocystis*) with a light intensity of 20 µmol m^−2^ s^−1^. When appropriate, 50 µg ml^−1^ kanamycin (Km), 2.5 µg ml^−1^ spectinomycin (Sp), 2.5 µg ml^−1^ streptomycin (Sm) or 30 µg ml^−1^ neomycin (Nm) was added. Non-segregated Δ*syc1139* cells were always grown in the presence of Km. *E. coli* strains DH5α, DH5αMCR, XL1-blue and HB101 were used for cloning and conjugation by triparental mating. BTH101 was used for BACTH assays and BL21 (DE3) was used for expression of His- and GFP-tagged proteins in *E. coli*. All *E. coli* strains (Supplementary Table 2) were grown in LB medium containing the appropriate antibiotics at standard concentrations.

### Plasmid and strain construction

All plasmids employed in this study were either generated by using standard restriction enzyme-base cloning procedures or using Gibson assembly (Gibson *et al.*, 2009). A detailed description of the cloning strategies for the respective plasmids is available upon request from the authors. All primers, plasmids and strains employed or generated in this study are listed in Supplementary Tables 2-5. GFP, YFP and eCFP protein tags were used as reporter proteins and His_6_ tag was used for protein affinity purification. For gene replacement mutants, homologous flanks for double homologous recombination comprised 1000 bp upstream and downstream of the gene of interest. Mutant strains harboring gene replacements with antibiotic resistance cassettes (*nptII* or *CS.3*; Beck *et al.*, 1982; Sandvang, 1999) were verified by colony PCR testing for absence of gene of interest using primers #129/#130 for Δ*slr7083*, primers #168/#169 for Δ*slr1301*, primers #146/#147 for Δ*syc2039* or primers #161/#162 for Δ*syc1139*. We also attempted to generate gene replacement mutants for *all4981* and *fm7001* but remained unsuccessful.

### Transformation of cyanobacteria

Transformation of *Synechococcus* was achieved by natural transformation as described by Ivleva *et al.* (2005) and transformation of *Synechocystis* was accomplished by natural transformation as described by Vermaas *et al.* (2002) or by conjugation as described by Ungerer and Pakrasi (2016). *Anabaena* and *Fischerella* were transformed by conjugation as described by Ungerer and Pakrasi (2016) or Stucken *et al.* (2012), respectively. Ex-conjugant colonies from *Synechococcus* and *Synechocystis* carrying gene replacements were re-streaked three to four times and absence of genes of interest was verified by colony PCR. Transformation of sonicated (fragmented) and NaCl-treated *Fischerella* cells followed by the conjugational method described by Ungerer and Pakrasi (2016) was also feasible for *Fischerella*, albeit with a lower transformation frequency.

### Phenotypic characterization of the mutant strains

Defects in cell viability were evaluated by spot assays adapted from Dörrich *et al.* (2014). Wild type and mutant strains from liquid cultures or BG11 plates were adjusted to an OD_750_ of about 0.4 in liquid BG11 liquid. Next, 5 µl of cells were spotted in triplicates onto BG11 plates or BG11 plates supplemented with Proteinase K or lysozyme at indicated concentrations in 10-fold serial dilutions and incubated under standard growth conditions until no further colonies arose in the highest dilution.

Growth defects were assessed with growth curves. For this, cells were grown in liquid BG11 medium, washed three times by centrifugation (6500 × *g*, RT, 3 min) in BG11, adjusted to an OD_750_ of 0.1 and then grown in triplicates or quadruples at standard growth conditions in 15 ml culture volumes. OD_750_ values were recorded every 24 h.

Cell length of *Synechococcus* WT, mutant strains and mutant complementation strains was measured using the line tool from the imaging software Fiji.

Cell wall integrity defects were evaluated by testing the influence of osmotic factors on cell growth. *Synechococcus* WT and mutant strains were grown on BG11 agar plates, transferred to BG11 liquid medium and grown under standard growth conditions with or without 5 mM glucose, 200 mM glucose, 2 mM NH_4_Cl, 200 mM maltose or 500 mM NaCl.

To evaluate the motility of *Synechocystis* and PCC-M WT and mutant strains, three single colonies of the respective strain were streaked on a line on a BG11 growth plate. Growth plates were then placed into the standard culture incubator for 10 d with with illumination limited from one direction.

### Protein purification and *in vitro* filamentation assays

C-terminally His_6_-tagged proteins were expressed and subsequently purified under denaturing conditions using Ni-NTA affinity columns as previously described by Springstein *et al.* (2019). For expression of MBP-Fm7001-His_6_, DH5α cells carrying pMAL-c2x-Fm7001-His_6_ were grown and induced accordingly but in the presence of 0.2% glucose. Purified proteins were dialyzed overnight against polymerization buffer (PLB: 50 mM PIPES, 100 mM KCl, pH 7.0; HLB: 25 mM HEPES, 150 mM NaCl, pH 7.4) at 18 °C and 180 rpm with three bath changes using a Slide-A-Lyzer™ MINI Dialysis Device (10K MWCO, 0.5 ml or 2 ml; Thermo Fischer Scientific). Purified proteins were stained with 0.005 mg NHS-Fluorescein (Thermo Fischer Scientific) per 1 ml protein dialysate and *in vitro* filamentation was analyzed by epifluorescence microscopy.

For Fm7001-His_6_, proteins were slowly dialyzed against 2 mM Tris-HCl, 4.5 M urea, pH 7.5 (18°C, 200 rpm) decreasing 0.5 M urea every 2 h (from 6 M to 4.5 M urea). The resulting floating filamentous web was then analyzed by bright field microscopy.

Syc2039-His_6_ failed to be expressed in *E. coli* BL21 (DE3). To bypass this, Syc2039-GFP-His, under the control of an IPTG-inducible P_trc_, was inserted into a neutral locus of *Synechococcus*. Cells were grown to an OD_750_ of 0.8 and protein expression was induced with 0.05 mM IPTG for 3 d. Induced cells were harvested and washed with PBS by centrifugation (4800 × *g*, 4 °C, 10 min) and stored at −80 °C. Protein purification, dialysis and labeling was then performed as described above with the exception that BG11 growth medium was used as dialysate.

### Co-immunoprecipitation

For co-immunoprecipitations of fluorescently tagged CCRP candidates, cyanobacterial strains expressing YFP-All4981 or Slr1301-YFP were grown in BG11 or BG11_0_ liquid medium. Co-immunoprecipitation was performed using the μMACS GFP isolation kit (Miltenyl Biotec) as previously described by Springstein *et al.* (2019) using PBS-N (PBS supplemented with 1% NP-40) or HSLB (50 mM NaH_2_PO_4_, 500 mM NaCl, 1% NP-40, pH 7.4) lysis buffers supplemented with a protease inhibitor cocktail (cOmplete™, EDTA-free Protease Inhibitor Cocktail, Sigma-Aldrich). Proteins were identified by mass spectrometry as previously described by Springstein *et al.* (2019) for YFP-All4981 or by Kahnt *et al.* (2007) for Slr1301-YFP.

### Immunofluorescence

The localization of FtsZ in *Synechococcus* WT and mutant strains was evaluated by immunofluorescence using a modified protocol from Heinz *et al.* (2016). In contrast, cells were lysed in 50 mM Tris-HCl pH 7.4, 10 mM EDTA and 0.2 mg ml^−1^ lysozyme for 30 min at 37 °C and samples were blocked in 1x Roti®-ImmunoBlock (Carl Roth) in PBS supplemented with 0.05% Tween 20. Samples were incubated with rabbit anti-FtsZ primary antibody (Agrisera; raised against *Anabaena* FtsZ; 1:250 diluted) in blocking buffer followed by incubation with 7.5 µg ml^−1^ Alexa Fluor 488-conjugated goat anti-rabbit IgG (H+L) secondary antibody (Thermo Fischer Scientific) in blocking buffer. Before microscopy, cells were stained with 10 µg ml^−1^ DAPI (final concentration) in PBS.

### Brightfield and fluorescence microscopy analysis

Bacterial strains grown in liquid culture were either directly applied to a microscope slide or previously immobilized on a 2% low-melting agarose in PBS agarose pad and air dried before microscopic analysis. Epifluorescence microscopy was performed using an Axio Imager.M2 light microscope (Carl Zeiss) equipped with Plan-Apochromat 63x/1.40 Oil M27 objective and the AxioCam MR R3 imaging device (Carl Zeiss). GFP, Alexa Fluor 488, eCFP and YFP fluorescence was visualized using filter set 38 (Carl Zeiss; excitation: 470/40 nm band pass (BP) filter; emission: 525/50 nm BP). Chlorophyll auto-fluorescence was recorded using filter set 15 (Carl Zeiss; excitation: 546/12 nm BP; emission: 590 nm long pass). When applicable, cells were previously incubated in the dark at RT for about 5 min with 10 µg ml^−1^ DAPI in PBS to stain intracellular DNA. For visualization of DAPI fluorescence filter set 49 (Carl Zeiss; excitation: G 365 nm; emission: 455/50 nm) was employed. *E. coli* BL21 (DE3) cells expressing C-terminally GFP-tagged protein candidates were grown over night in LB and then diluted 1:40 in the same medium the following day. Cells were grown for 2 h at 37 °C, briefly acclimated to 20 °C for 10 min and induced with 0.05 mM IPTG at 20 °C. Protein localization of GFP/YFP-tagged proteins was then observed after indicated time points of cells immobilized on an agarose pad.

### Statistical analysis

Beta-galactosidase values were measured in triplicates from three independent colonies and significant differences compared to WT were determined by a one-way ANOVA using Dunnett’s multiple comparison test. For statistical evaluation of *Synechococcus* WT and mutant cell length, a one-way ANOVA using Turkey’s multiple comparison test was used. Significance levels are the same as for the beta-galactosidase assay. Statistical tests were performed with the GraphPad Prims 8.0.0 software. Significance levels are indicated by stars (*) and correspond to: *: P < 0.05, **: P < 0.01, ***: P < 0.001, ****: P < 0.0001.

### RNA isolation and RT-PCR

Total RNA was isolated from 10 ml culture using either the Direct-zol™ RNA MiniPrep Kit (Zymo Research; *Synechocystis*, *Synechococcus* and *Anabaena*) according to the manufacturer’s instructions or the Plant RNA Reagent (Thermo Fischer Scientific; *Anabaena*, *Fischerella* and *Synechocystis*). For RNA isolation using the Plant RNA Reagent, a modified protocol was employed. To this end, cells were pelleted by centrifugation (4800 × *g*, 10 min, 4 °C) and the supernatant was discarded. The pellet was resuspended in 0.5 ml of Plant RNA Reagent und lysed in a Precellys® 24 homogenizer (Bertin) with 3 strokes at 6500 rpm for 30 s in 2 ml soil grinding (SK38) or tough microorganism (VK05) lysis tubes (Bertin). RNA was then isolated according to the manufacturer’s instructions. Isolated RNA was treated with DNA-free™ Kit (2 units rDNAs/reaction; Thermo Fischer Scientific) and 1 µg (*Fischerella*, *Synechocystis* and *Synechococcus*) or 200 ng (*Anabaena*) RNA was reverse transcribed using the Maxima™ H Minus cDNA Synthesis Master Mix (with dsDNase; Thermo Fischer Scientific, for *Fischerella*, *Synechocystis* and *Synechococcus*) or the qScript™ cDNA Synthesis Kit (Quanta Biosciences, for *Anabaena*). RT-PCR of cDNA samples for *fm7001*, *ftsZ*, *slr7083*, *rnpB*, *slr1301*, *syc2039*, *syc1139*, *all4981*, *all4981*+*all4982* and *all4981*+*all4983* was done using primer pairs #1/#2, #3/#4, #5/#6, #7/#8, #9/#10, #11/#12, #13/#14, #15/#16, #17/#15 and #18/#15, respectively.

### Bacterial two hybrid assays

In this study, the BACTH system (Euromedex) was employed. Gene candidates were cloned into the expression vectors pKNT25, pKT25, pUT18 and pUT18C by GIBSON assembly, thereby generating C and N-terminal translational fusions to the T25 or T18 subunit. Chemically competent *E. coli* BTH101 (Δ*cya*) cells were co-transformed with 5 ng of the indicated plasmids, plated onto LB plates supplemented with 200 µg ml^−1^ X-gal, 0.5 mM IPTG, Amp, Km and grown at 30 °C for 24-36 h. Interactions were quantified by beta-galactosidase assays from three colonies for each combination according to the protocol described by Euromedex or in a 96 well format according to Karimova, Davi and Ladant (2012). For this aim, cultures were either grown over night at 30 °C or for 2 d at 20 °C in LB Amp, Km, 0.5 mM IPTG and interaction strength of the investigated proteins was by quantified by beta-galactosidase-mediated hydrolyzation of ONPG (ortho-Nitrophenyl-β-galactoside), which is then recorded in Miller units (Miller, 1992).

## Acknowledgements

We thank Katrin Schumann, Myriam Barz, Lisa Stuckenschneider, Lisa-Marie Philipp and Marius Lasse Theune for their assistance in the experimental work. Furthermore, we thank Martin Thanbichler and Daniela Kiekebusch (both from Philipps University, Marburg, Germany) for their support with mass spectrometry analysis. The study was supported by the German science foundation (DFG) (Grant No. STU513/2-1 awarded to KS).

## Author contribution

BLS and KS designed the study. BLS established and performed the experimental work with contributions from JW. CW and TD performed the comparative genomics analysis. AOH analyzed protein samples by mass spectrometry. BLS, TD and KS drafted the manuscript with contributions from all coauthors.

## Supplementary Information

**Supplementary Fig. 1:**
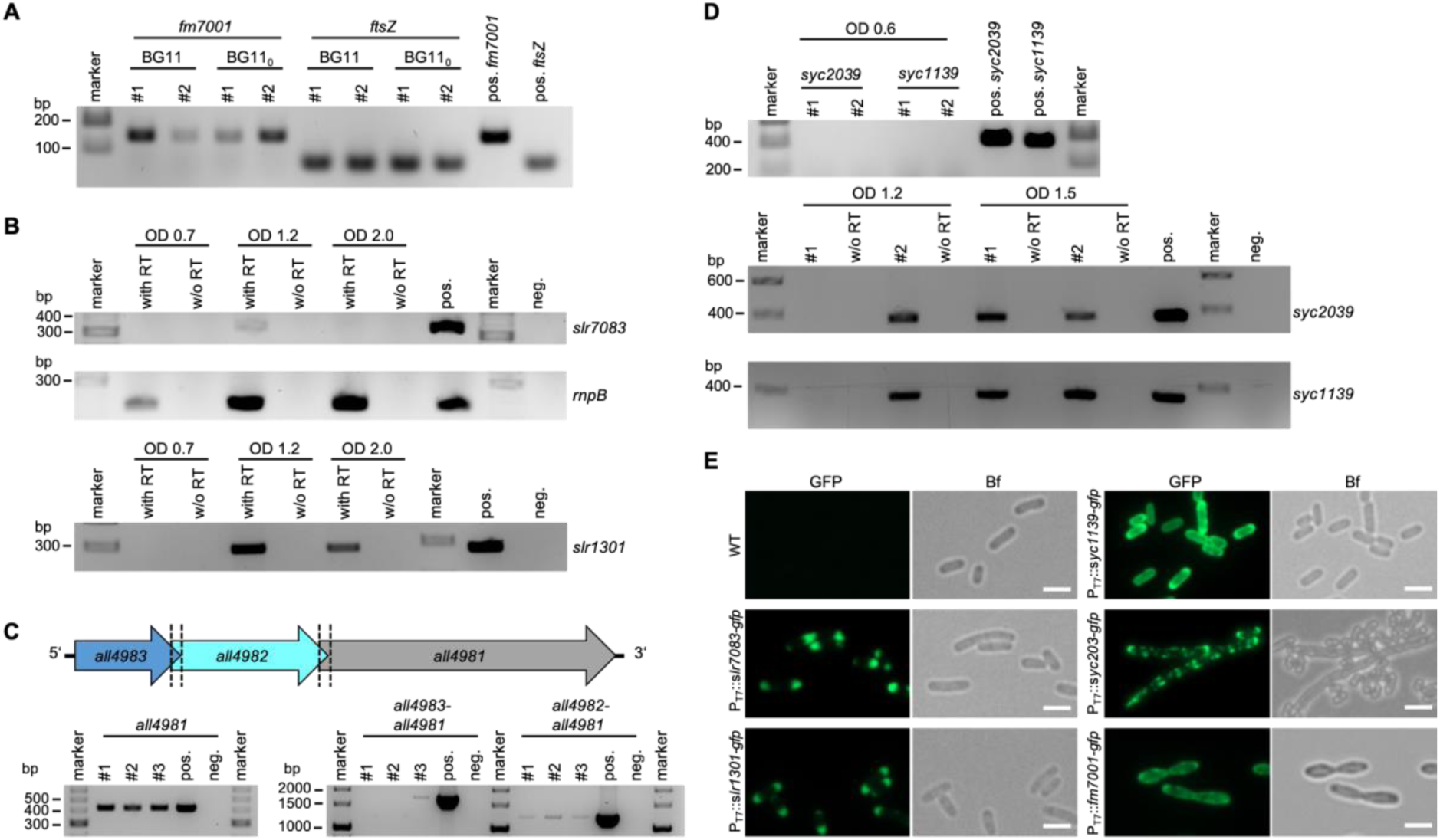
Expression of candidate CCRPs and heterologous expression in *E. coli*. (**A**) RT-PCR of reverse transcribed whole RNA from young *Fischerella* WT cultures grown in BG11 or BG110 from two independent biological replicates. Gene transcripts were verified using internal *fm7001* gene primers (#1/#2) or internal *ftsZ* gene primers (#3/#4) as a control. (**B**) RT-PCR of reverse transcribed whole RNA from *Synechocystis* WT (OD_750_ 0.7, 1.2 or 2.0) grown in BG11 using internal *slr7083* gene primers (#5/#6) or internal *slr1301* gene primers (#9/#10). Internal *rnpB* gene primers (#7/#8) were included as a control. (**C**, top) Schematic representation of the genomic context of *all4981*. The 3’ end of *all4983* overlaps with 4 bp with the 5’ region of *all4982*, which has the same overlap with the 5’ end of *all4981*. Both overlaps are comprised of the same four nucleotides (ATGA). (**C**, bottom) RT-PCR of reverse transcribed whole RNA from *Anabaena* WT cultures grown in BG11 (OD_750_ 1.8) from three independent biological replicates. *all4981* gene transcript was internal gene primers (#15/#16). For operon structure of *all4983-all4981* or *all4982-all4981*, primer pairs #17/#16 or #18/#16 were used, respectively. Only one replicate showed a common transcript for an *all4983-all4981* operon, which is likely is the result of the long fragment (about 1800 bp). The employed cDNA synthesis kit is optimized for fragments up to 1000 bp, thus making longer reverse transcriptions unlikely. (**D**) RT-PCR of reverse transcribed whole RNA from *Synechococcus* WT (OD_750_ 0.6, 1.2 or 1.5) grown in BG11 from two independent biological replicates. Gene transcripts were verified using internal *syc2039* gene primers (#11/#12) and internal *syc1139* gene primers (#13/#14). (**B**,**D**) RNA was either reverse transcribed in the reaction buffer containing reverse transcriptase (with RT) or without reverse transcriptase (w/o RT) as a control for residual genomic DNA contamination. (**A-D**) Genomic DNA of the respective species was included as positive control for the different reactions. (**E**) GFP fluorescence and bright field micrographs of *E. coli* BL21 (DE3) cells expressing Slr7083-GFP, Slr1301-GFP, Syc1139-GFP, Syc2039-GFP or Fm7001-GFP. Cells were grown at 20 °C or (Fm7001-GFP) 16 °C and protein expression was induced with 0.05 mM IPTG for 24 h. Scale bars: 2.5 µm.

**Supplementary Fig. 2:**
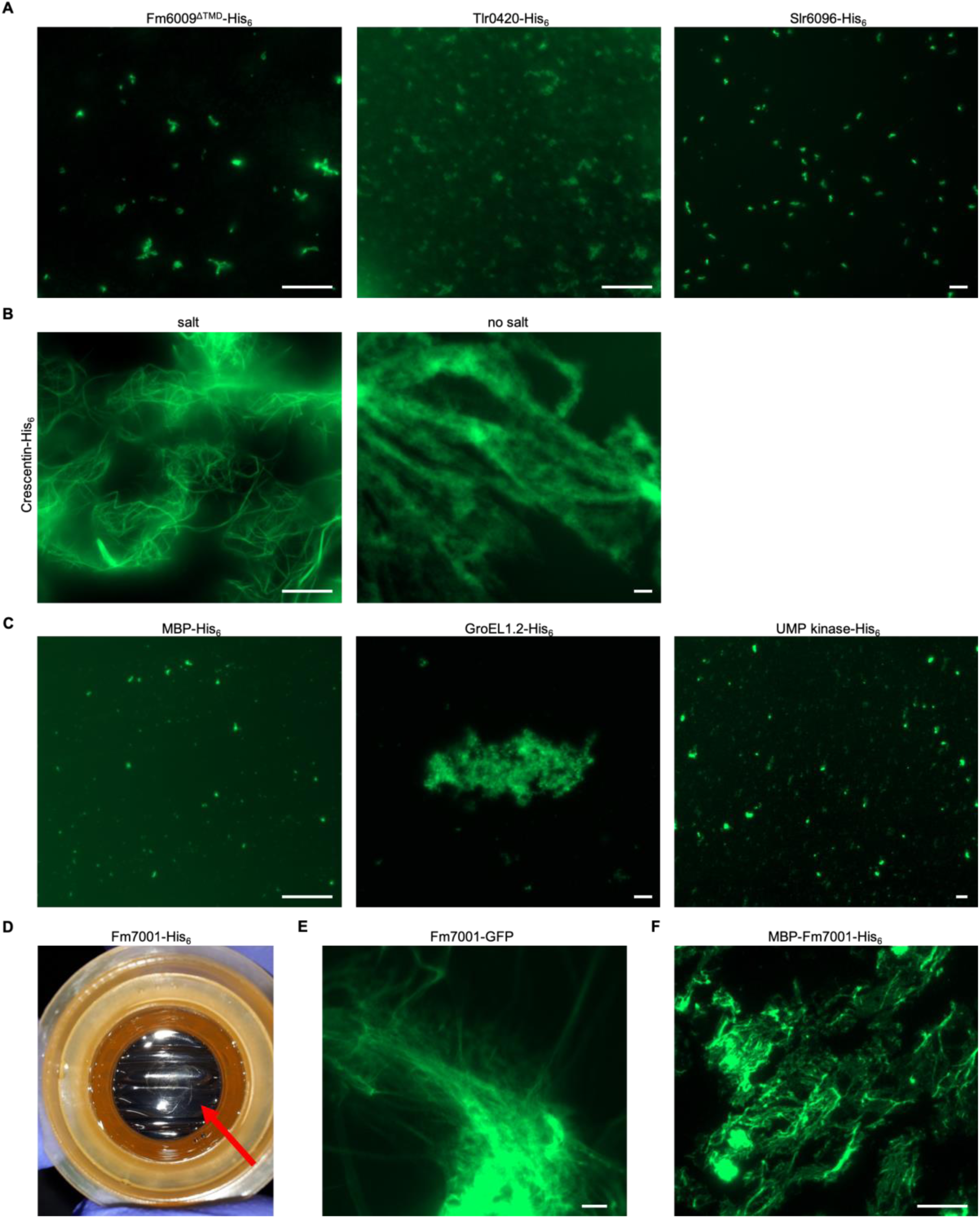
*In vitro* polymerization is dependent on monovalent ions. (A-C) NHS-fluorescein fluorescence micrographs of *in vitro* structures formed by purified and renatured Fm6009^ΔTMD^-His_6_ (lacking the transmembrane domain, i.e. the first 91 aa), Tlr0420-His_6_ or Slr6096-His_6_ (1 mg ml^−1^ each), Crescentin-His_6_ (0.7 mg ml^−1^), MBP-His_6_ (1 mg ml^−1^), GroEL1.2 (0.7 mg ml^−1^) or UMP kinase (0.5 mg ml^−1^), in HLB or Crescentin-His_6_ (0.7 mg ml^−1^) renatured in 25 mM Hepes, pH 7.4. Note: Crescentin-His_6_ *in vitro* polymerization into smooth filaments is strictly dependent of the presence of salt in the renaturation buffer as Crescentin-His_6_ without salt assembles into filamentous aggregates only. (**A**) Proteins were dialyzed in a stepwise urea-decreasing manner and stained with an excess of NHS-Fluorescein. (**D**) Bright field micrograph of a sheet-like flat object floating on top of the dialysate (red arrow) formed upon dialysis of Fm7001-His_6_ (0.7 mg ml^−1^) into 2 mM Tris-HCl, 4.5 M urea, pH 7.5. (**E,F**) Epifluorescence micrographs of filamentous structures formed by (**E**) denatured cell-free extracts of *E. coli* BL21 (DE3) expressing Fm7001-GFP (0.7 mg ml^−1^ whole protein) dialyzed into 2 mM Tris-HCl, 3 M urea, pH 7.5 or by (**F**) natively purified MBP-Fm7001-His_6_ (0.8 mg ml^−1^) stained with NHS-fluorescein in HLB. Scale bars: 10 µm.

**Supplementary Fig. 3:**
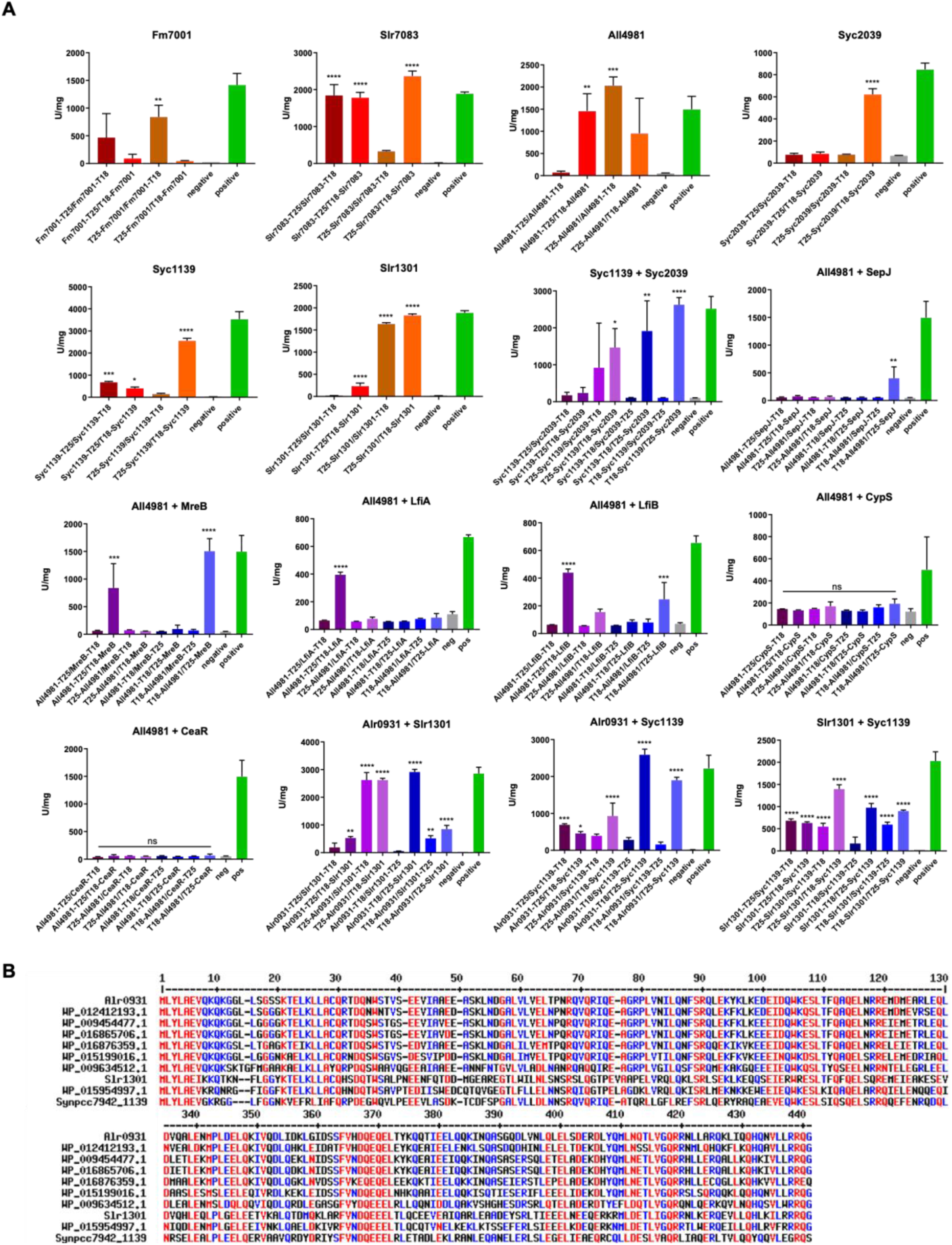
BACTH analysis of cyanobacterial CCRPs. (**A**) Beta-galactosidase assays (BACTH) of *E. coli* BTH101 cells co-expressing indicated T25 and T18 translational fusions of all possible pair-wise combinations from three independent colonies grown for 1 d at 30 °C or 2 d at 20 °C. Quantity values are given in Miller Units per milligram LacZ of the mean results from three independent colonies. Negative: N-terminal T25 fusion construct of the respective protein co-transformed with empty pUT18C. Positive: Zip/Zip control. Error bars indicate standard deviations (n=3). Values indicated with * are significantly different from the negative control. *: P < 0.05, **: P<0.01, ***: P<0.001, ****: P<0.0001 (one-way ANOVA using Dunnett’s multiple comparison test). (**B**) Multiple sequence alignment of selected cyanobacterial homologous CCRPs using MULTALIGN (Corpet, 1988). Alr0931 (termed CypS; *Anabaena*), Slr1301 (*Synechocystis*) and Synpcc7942_1139 (*Synechococcus*) are identified by their designated cyanobase locus tag. Other proteins are given as NCBI accession numbers. WP_012412193.1 (*Nostoc punctiforme* PCC 73102), WP_009454477.1 (*Fischerella thermalis* PCC 7521), WP_016865706.1 (*Fischerella*), WP_016876359.1 (*C. fritschii* PCC 9212), WP_015199016.1 (*Calothrix* sp. PCC 6303), WP_009634512.1 (*Synechocystis* sp. PCC 7509), and WP_015954997.1 (*Cyanothece* sp. PCC 7424). Amino acids from 1-130 and 334-441 are depicted. Red highlighted amino acid residues are conserved among all listed species, blue amino acids are mostly conserved, and black amino acids are not conserved. Characteristic for this group of conserved cyanobacterial CCRPs is a highly conserved N-terminus with a M-L-Y-L-A-E-V sequence motif present in nearly all homologs, followed by a moderately conserved N-terminal region of the first 120 amino acids. Two other highly conserved domains are present in this group, one located around the centre of the proteins (between the 340^th^ and 370^th^ amino acid), and another one shortly thereafter between the 400^th^ and 420^th^ amino acid.

**Supplementary Fig. 4:**
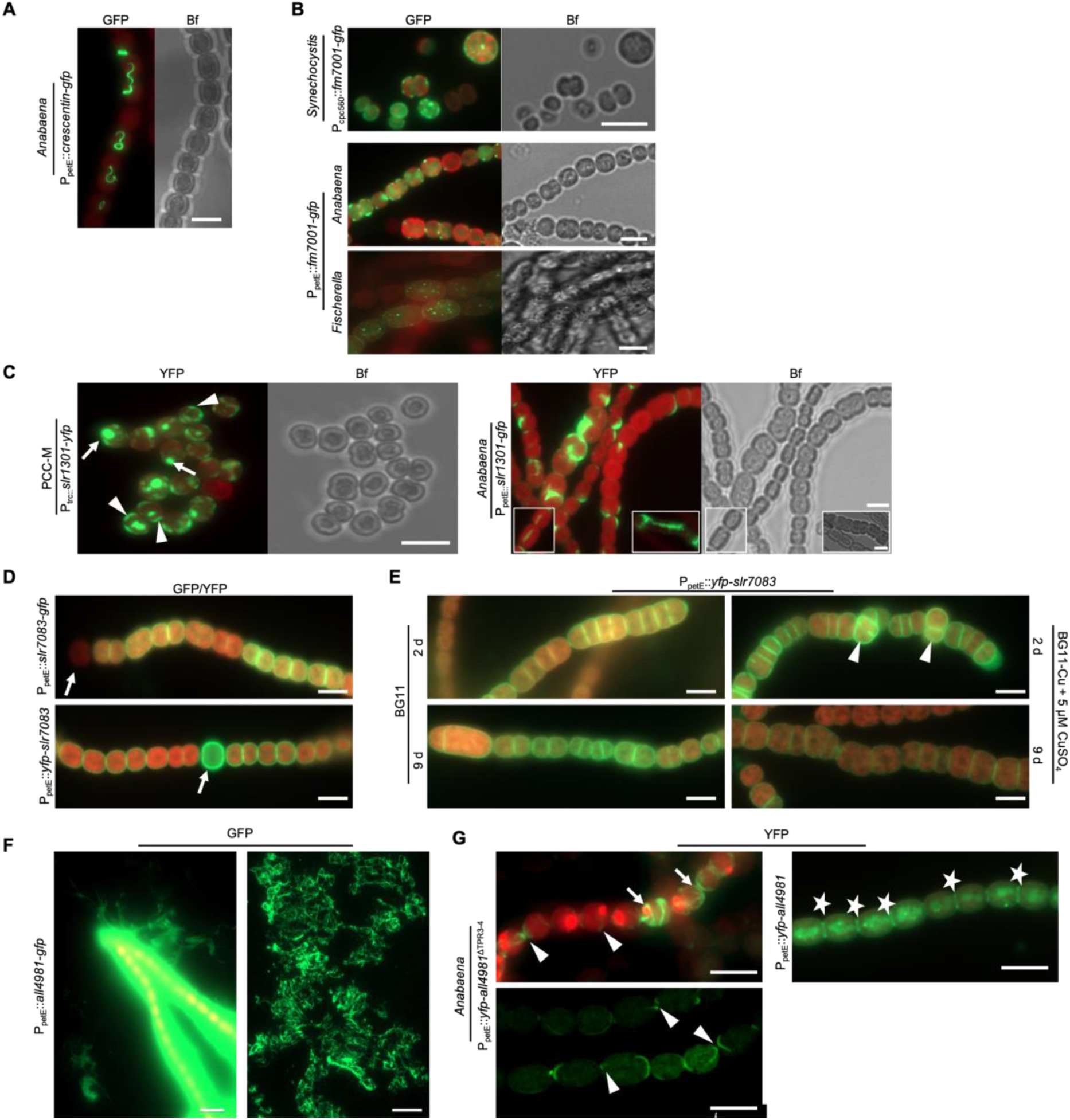
Expression of candidate CCRPs in different cyanobacterial species. GFP fluorescence, chlorophyll autofluorescence (red) and bright field micrographs of *Synechocystis*, *Anabaena* or *Fischerella* cells expressing (**A**) Crescentin-GFP, (**B**) Fm7001-GFP, (**C**) Slr1301-GFP, (**D**) Slr7083-GFP, (**E**) YFP-Slr7083, (**F**) All4981-GFP or (**G**) YFP-All4981^ΔTPR3-4^ or YFP-All4981 from P_petE_, P_trc_, or P_cpc560_. (**B**) Unlike N-terminal fusion with a YFP tag, no *in vivo* filaments can be observed upon C-terminal fusion of Fm7001 with a GFP-tag in any tested cyanobacterium. (**C**) Besides intracellular filaments (figure inlays), Slr1303 accumulated at the periphery within *Anabaena* cells and induced a partial swollen cell phenotype. White triangles mark crescent-like localizations. White arrows show Slr1301-YFP accumulations. (**D)** *Anabaena* cells were grown on BG110 plates. White arrows indicate heterocysts. (**E**) *Anabaena* cells expressing YFP-Slr7083 from P_petE_ grown in liquid BG11 or liquid BG11 without copper and induced with 5 µM CuSO_4_ for 2 and 9 d. White triangles point to multiseriate *Anabaena* trichome growth upon protein overexpression. (**F**) *Anabaena* cells expressing All4981-GFP from P_petE_ grown in BG11 supplemented with 0.5 µM CuSO_4_ for 2 d. Extended period of overexpression of All4981-GFP led to cell rupture. Protein filaments released from the *Anabaena* trichome are shown in the left image while the right image shows extracellular (*ex vivo*) filaments observed in the growth medium. (**G**) *Anabaena* cells were grown in BG110 supplemented with 0.5 µM CuSO_4_. White triangles indicate selected filaments traversing through the cells. White arrows point to spindle-like YFP-All4981^ΔTPR3-4^ filaments. White stars mark septal localizations. Scale bars: (**A-D**,**G**) 5 µm, (**F**, left) 10 µm or (**F**, right) 20 µm.

**Supplementary Fig. 5:**
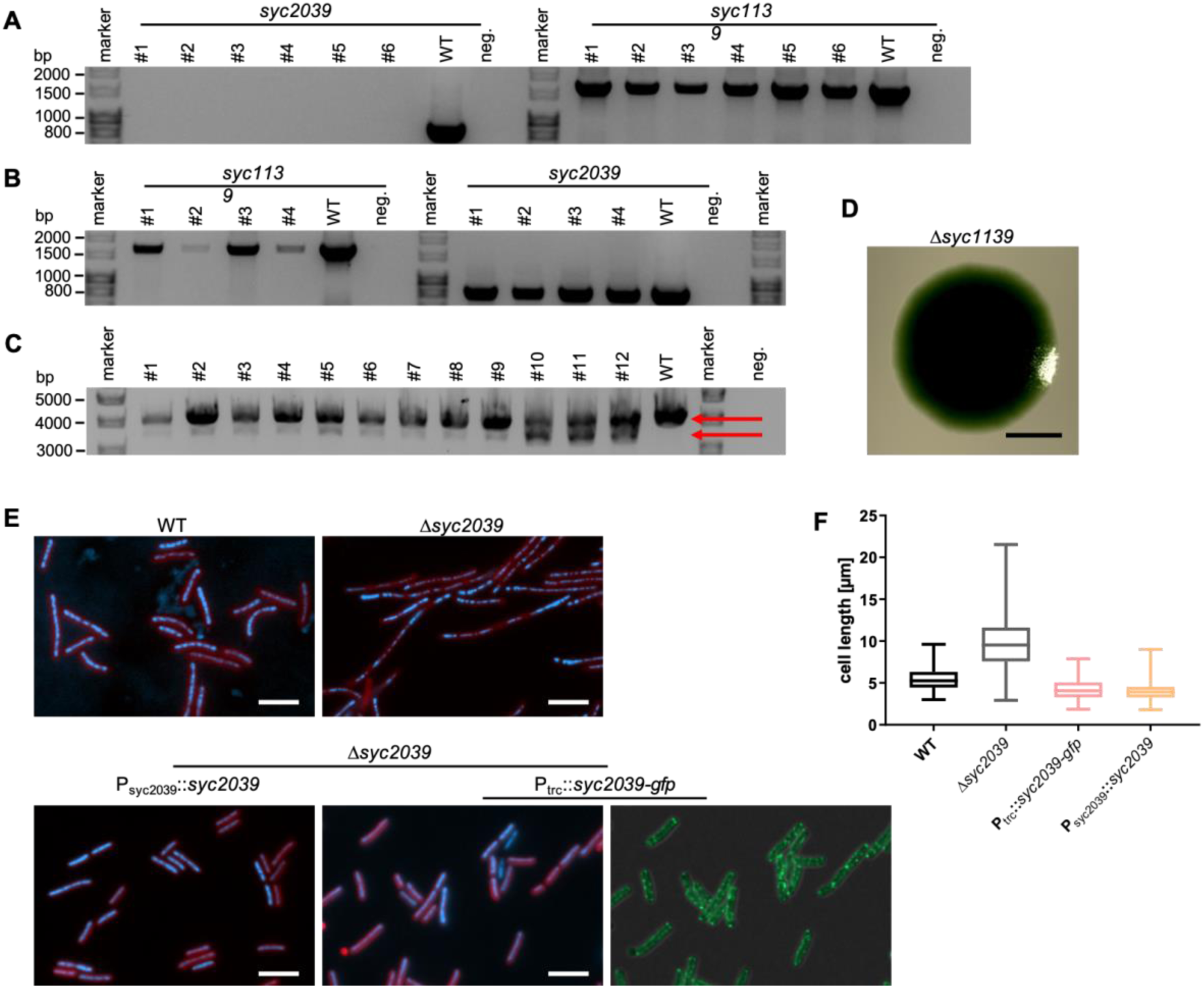
Verification of *Synechococcus* CCRP mutants. (**A**) Colony PCR of six Δ*syc2039* mutant clones using *syc2039* gene primers (#149/#147) and *syc1139* gene primers (#161/#162) as a control. (**B**) Colony PCRs of four non-segregated Δ*syc1139* mutant clones using *syc1139* gene primers (#174/#175) or *syc2039 gene* primers (#159/#160) as a control. (**C**) Colony PCR of twelve non-segregated Δ*syc1139* mutant clones using primers encompassing the homologous flanking regions used for homologous recombination (#238/#239). Upper red arrow indicates WT allele PCR product. Lower red arrow indicates Δ*syc1139* mutant PCR product. As a positive control, *Synechococcus* genomic DNA was included. (**D**) Growth of Δ*syc1139* mutant on non-selective plates leads to a reversal to WT phenotype. (**E**) Merged DAPI fluorescence and chlorophyll autofluorescence (red) and merged GFP fluorescence and bright field micrographs of *Synechococcus* WT, Δ*syc2039* mutant and Δ*syc2039* mutant complemented with P_syc2039_::*syc2039* or P_trc_::*syc2039-gfp* inserted into the neutral NS1 locus. Cells were grown in BG11 or BG11 supplemented with 0.001 mM IPTG (for strain carrying P_trc_::*syc2039-gfp*) and stained with 10 μg ml^−1^ DAPI. (**F**) Cell length of *Synechococcus* WT (n=505), Δ*syc2039* mutant (n=517), Δ*syc2039* mutant carrying P_trc_::*syc2039-gfp* (n=547) and Δ*syc2039* mutant carrying P_syc2039_::*syc2039* (n=529) cells.

**Supplementary Fig. 6:**
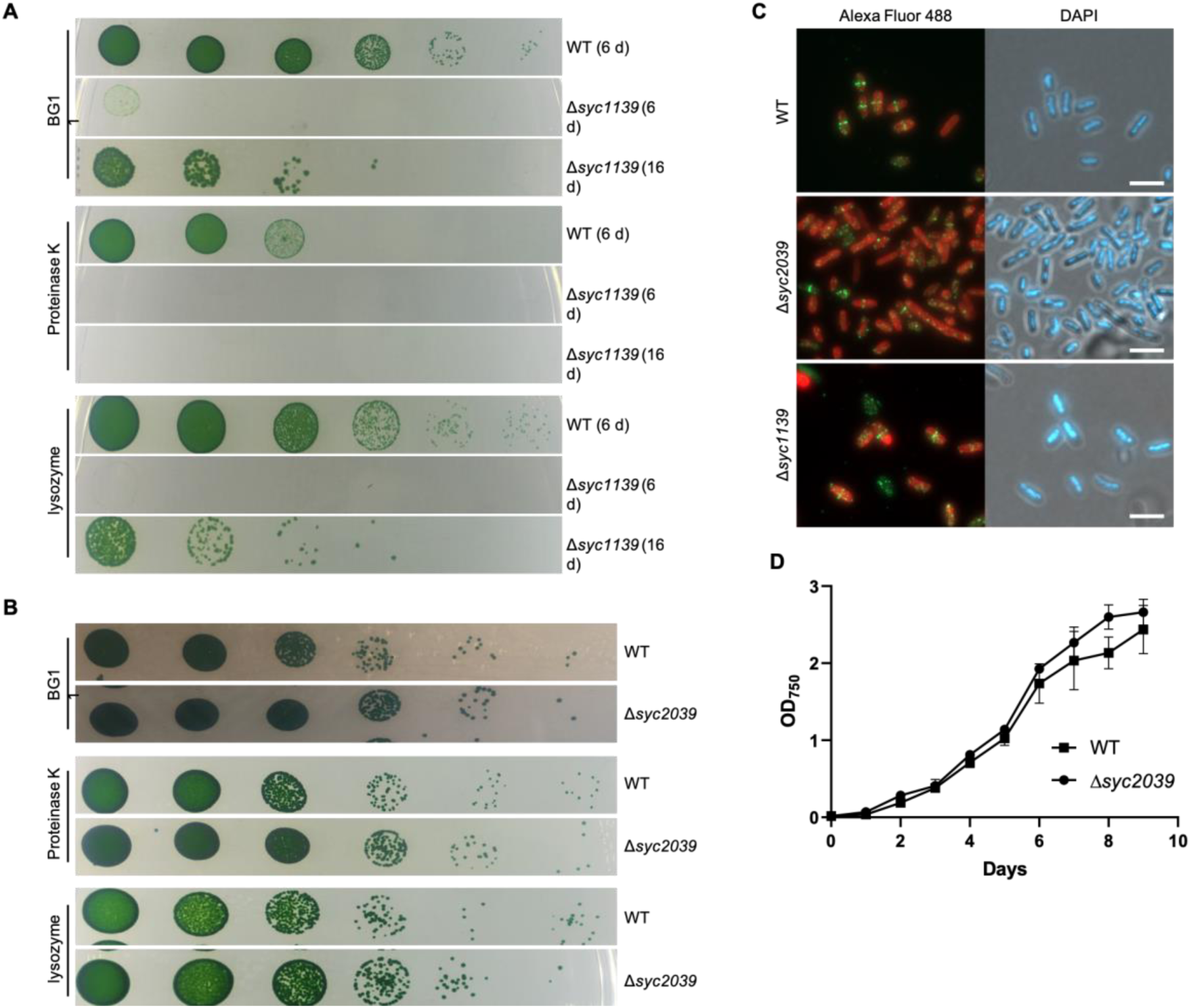
Phenotypic characterization of *Synechococcus* mutant strains. (**A**) *Synechococcus* WT (upper lane, after 6 days) and non-segregated Δ*syc1139* mutant (middle lane: after 6 days and lower lane after 16 days) strains were grown on BG11 plates or BG11 plates supplemented with 50 µg ml^−1^ Km. Cells were resuspended in BG11, adjusted to an OD_750_ of 0.4 and spotted in triplicates of serial 10-fold dilutions on BG11 plates or BG11 plates supplemented with 100 µg ml^−1^ Lysozyme or 50 µg ml^−1^ Proteinase K. Cells were grown until no further colonies arose in the highest dilution. (**B**) *Synechococcus* WT and Δ*syc2039* mutant strains were grown in liquid culture at standard growth conditions until an OD_750_ of about 2.0, diluted in BG11 to an OD_750_ of 0.4 and spotted in triplicates of serial 10-fold dilutions on BG11 plates or BG11 plates supplemented with 100 µg ml^−1^ Lysozyme or 50 µg ml^−1^ Proteinase K. Cells were grown until no further colonies arose in the highest dilution. (**C**) Merged Alexa Flour-488 fluorescence and chlorophyll autofluorescence (red) and merged bright field and DAPI fluorescence micrographs of *Synechococcus* WT, Δ*syc2039* or non-segregated Δ*syc1139* mutant strains grown on BG11 plates and subjected to immunofluorescence staining using an anti-FtsZ primary antibody (Agrisera, raised against *Anabaena* FtsZ) and an Alexa Fluor-488 coated secondary antibody. Cells were mounted in Prolong Diamond antifade mountant with DAPI (Thermo Fischer Scientific). Scale bars: 5 µm. (**D**) Growth curve of *Synechocystis* WT and Δ*syc2039* mutant strain. Cells were grown in BG11, adjusted to an OD_750_ of 0.1 and then grown in triplicates at standard growth conditions. OD_750_ values were recorded once a day for 9 d. Error bars show the standard deviation (n=3).

**Supplementary Table 1:** Properties of cyanobacterial CCRPs.

| Gene /Locus tag | Genus | Subsection | Homologs distribution | Predicted proteins of similar structure (I-TASSER) | Homolog similarities | Conserved domains | Others |
| --- | --- | --- | --- | --- | --- | --- | --- |
| *crescentin* | *C. crescentus* | n/a | n/a | Cytoplasmic domain of bacterial cell division protein EzrA |  | SMC, CCDC158 | IF-like CCRP |
| *filP* | *S. coelicolor* | n/a | n/a | Dynein tail; α-Actinin; Tropomyosin | | DUF3552, SMC, RNase_Y | IF-like CCRP |
| *desmin* | *Homo sapiens* | n/a | n/a | PI4KIIIa lipid kinase |  | Filament (pfam00038), SMC, Spc7, MscS_TM | IF protein |
| *vimentin* | *Homo sapiens* | n/a | n/a | PI4KIIIa lipid kinase |  | Filament (pfam00038), SMC, Spc7 | IF protein |
| *syc2039* | *Synechococcus* | I | I | Tropomyosin |  | SMC, MukB, CALCOCO1 | N-terminal TMD; only in *Synechococcus* sp. |
| *syc1139* | *Synechococcus* | I | I, II, III, IV, V | Cytoplasmic domain of bacterial cell division protein EzrA | 39% | SMC, MukB, Spc7 | Homolog to *slr1301* |
| *slr6096* | *Synechocystis* | I | I, III, IV | Cytoplasmic domain of bacterial cell division protein EzrA |  | SMC |  |
| *slr7083* | *Synechocystis* | I | I | Plectin |  | SMC, MscS_TM | Encoded on pSYSA plasmid, only in *Synechocystis* sp. |
| *slr1301* | *Synechocystis* | I | I, II, III, IV, V | Cytoplasmic domain of bacterial cell division protein EzrA | 39% | SMC, SbcC, APG6, DUF3552 | Homolog to *syc1139* |
| *tlr0420* | *BP-1* | I | I, III | Plectin |  | SMC, MscS_TM |  |
| *fm7001* | *Fischerella* | V | IV, V | α-catenin or vinculin; similarity to acyl-CoA dehydrogenase | 63% | Acetyl-CoA carboxylase carboxyl transferase (PLN0322) | Highly expressed; 3' end 9 bp overlap to *fm7000* |
| *fm6009* | *Fischerella* | V | V | Structure of β-catenin and HTC-4 | | COG0610 | |
| *all4981* | *Anabaena* | IV | III, IV, V | TTC7B/Hyccin Complex, Clathrin | 47% | TPR | 5' with a 4 bp overlap to *all4982* |

The first column indicates the respective gene name or locus tags of each protein candidate. The second and third column indicate the respective subsection of the corresponding cyanobacterial genus. Column four lists the subsections that contain homologous proteins to the respective CCRP. Column five indicates structural similarities of the candidate to proteins in the Protein Data Bank (PDB) based on I-TASSER (Zhang, 2009; Yang and Zhang, 2015). The sixth column lists predicted sub-domains of protein candidates identified by BLAST Conserved Domain Search. Column seven names other features of interest. Abbreviations: (TMH) Transmembrane helix; (DUF) Domain of unknown function; (CCDC158) Coiled-coil domain-containing protein 158; (SMC) Structural maintenance of chromosomes; (MukB) The hinge domain of chromosome partition protein MukB; (APG6) Autophagy protein Apg6, (SbcC) DNA repair exonuclease SbcCD ATPase; (CALCOCO1) Calcium binding and coiled-coil domain; (Spc7) Spc7 kinetochore protein; (Filament) Intermediate filament protein; (TPR): Tetratricopeptide repeat; (PLN0322) Acetyl-CoA carboxylase carboxyl transferase; (COG0610) Type I site-specific restriction-modification system. *Anabaena* CCRPs CypS, LfiB, CeaR, Alr4393 and All4935 (Springstein *et al.*, 2019) also revealed structural similarities to EzrA. n/a: not applicable.

**Supplementary Table 2:** Bacterial strains.

| Strain | Genotype | Resistance | Reference |
| --- | --- | --- | --- |
| *E. coli* XL1 blue | *endA1 gyrA96(nal<sup>R</sup>) thi-1 recA1 relA1 lac glnV44 F[::Tn10 proAB<sup>+</sup> lacI<sup>q</sup> Δ(lacZ)M15] hsdR17(rK<sup>-</sup> mK<sup>+</sup>)* | Tet | Stratagene |
| *E. coli* HB101 | *F<sup>-</sup> mcrB mrr hsdS20(r<sub>B</sub><sup>-</sup> m<sub>B</sub><sup>-</sup>) recA13 leuB6 ara-14 proA2 lacY1 galk2 xyl-5 mtl-1 rpsL20(Sm<sup>R</sup>) glnV44 λ<sup>-</sup>* | Sm | (Boyer and Roulland-Dessoix, 1969) |
| *E. coli* DH5α | *F<sup>-</sup> Φ80lacZΔM15 Δ(lacZYA-argF) U169 recA1 endA1 hsdR17 (rK<sup>-</sup>, mK<sup>+</sup>) phoA supE44 λ<sup>-</sup> thi-1 gyrA96 relA1* |  | (Meselson and Yuan, 1968) |
| *E. coli* DH5αMCR | *F<sup>-</sup> endA1 supE44 thi-1 λ<sup>-</sup> recA1 gyrA96 relA1 deoR Δ(lacZYA-argF)U169 Φ80d lacZΔM15 mcrA Δ(mrr hsdRMS mcrBC)* |  | (Grant *et al.*, 1990) |
| *E. coli* BL21 (DE3) | *F<sup>-</sup> ompT gal dcm lon hsdS<sub>B</sub>(r<sub>B</sub><sup>-</sup> m<sub>B</sub><sup>-</sup>) λ(DE3 [lacI lacUV5-T7p07 ind1 sam7 nin5]) [malB<sup>+</sup>]<sub>K-12</sub>(λ<sup>S</sup>)* |  | (Studier and Moffatt, 1986)s |
| *E. coli* BTH101 | *F<sup>-</sup>, cya-99, araD139, galE15, galk16, rpsL1 (Str<sup>r</sup>), hsdR2, mcrA, mcrB1* | Sm | Euromedex |
| *Fischerella muscicola* PCC 7414 | WT |  | PCC, France |
| *Synechocystis* sp. PCC 6803 | Glucose tolerant Kazusa substrain WT |  | PCC, France |
| BLS4 | Δ*slr1301::CS.3* | Sm,Sp | This study |
| BLS5 | Δ*slr7083::CS.3* | Sm,Sp | This study |
| *Synechocystis* sp. PCC-M 6803 | Glucose tolerant and motile Moscow PCC-M substrain WT |  | A gift from Annegret Wilde (University Freiburg) |
| BLS6 | Δ*slr1301::CS.3* | Sm,Sp | This study |
| BLS7 | Δ*slr7083::CS.3* | Sm,Sp | This study |
| *Synechococcus elongatus* PCC 7942 | WT |  | A gift from Martin Hagemann (University Rostock) |
| BLS8 | Non-segregated Δ*sync2039::nptII* | Km | This study |
| BLS9 | Δ*sync2039::nptII* | Km | This study |
| *Anabaena* sp. PCC 7120 | WT |  | PCC, France |

**Supplementary Table 3:**
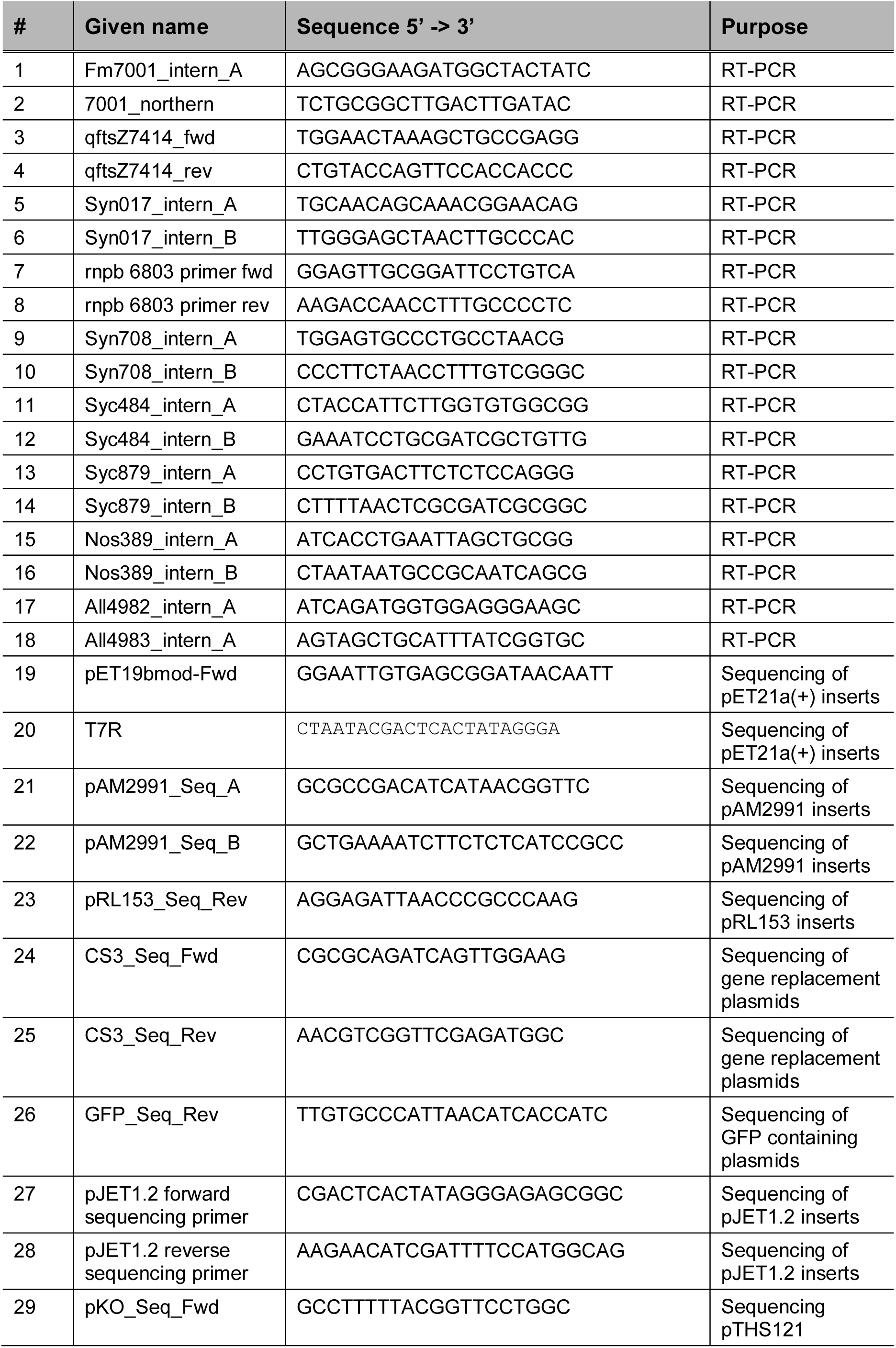
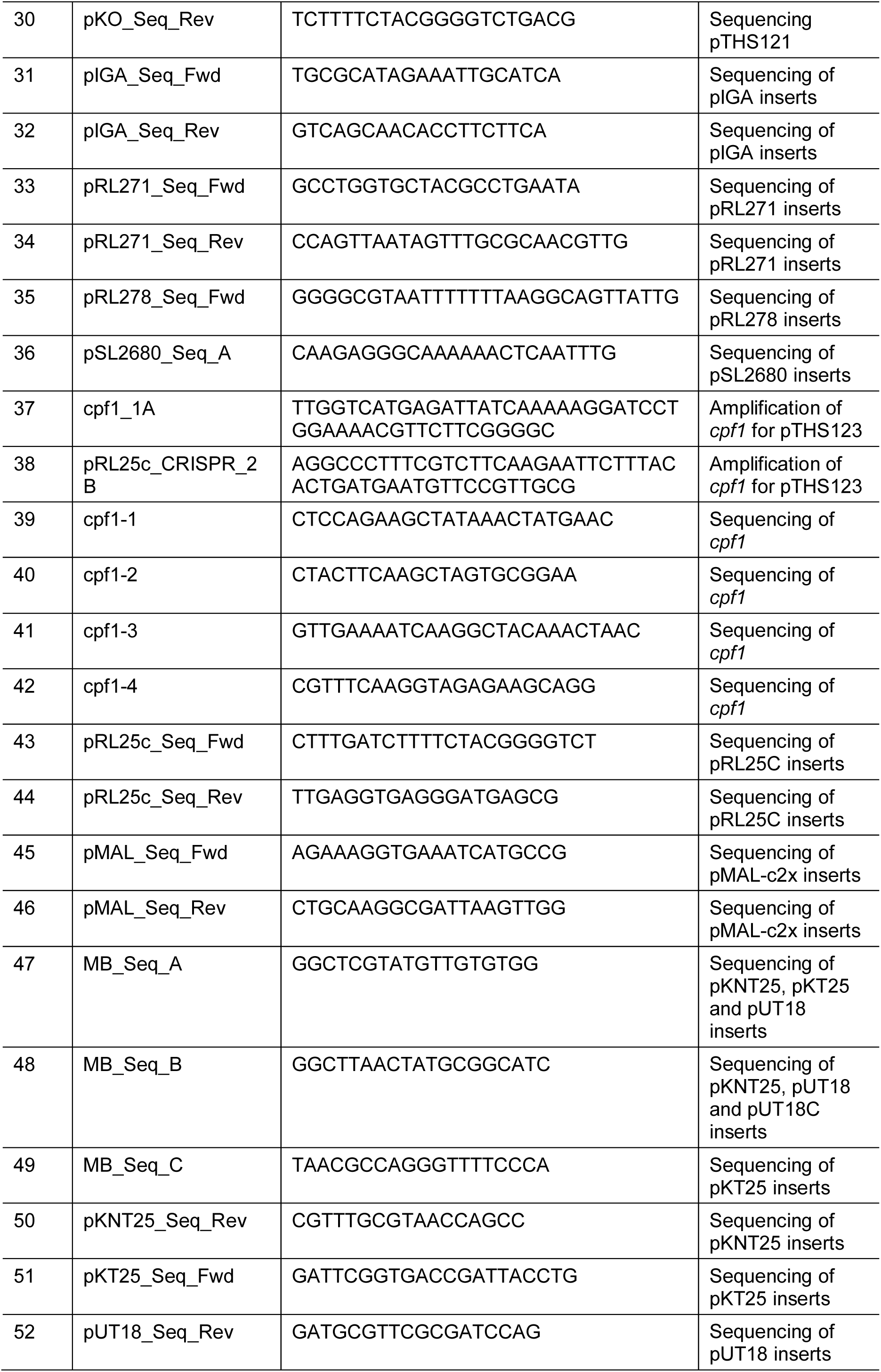
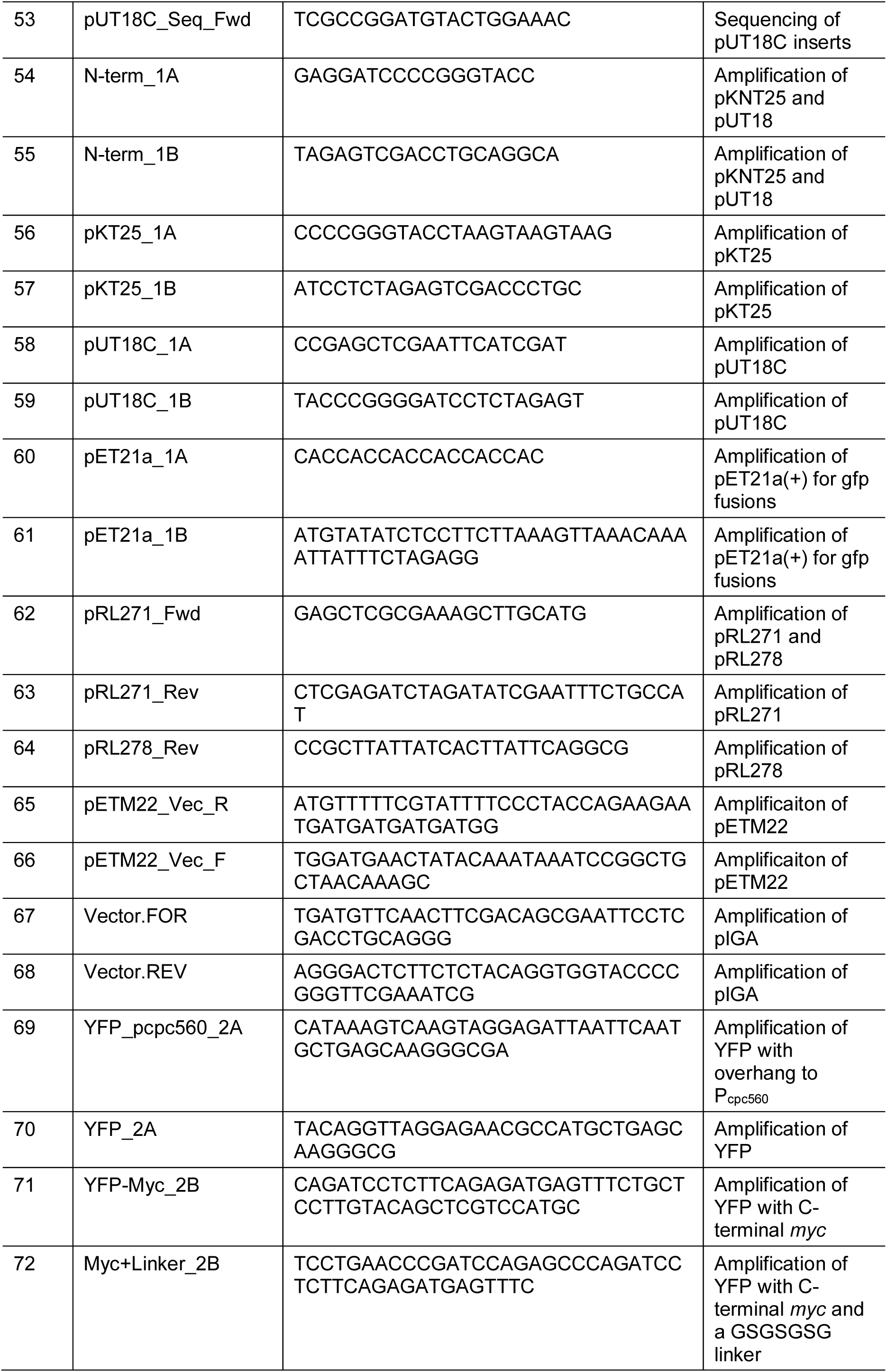
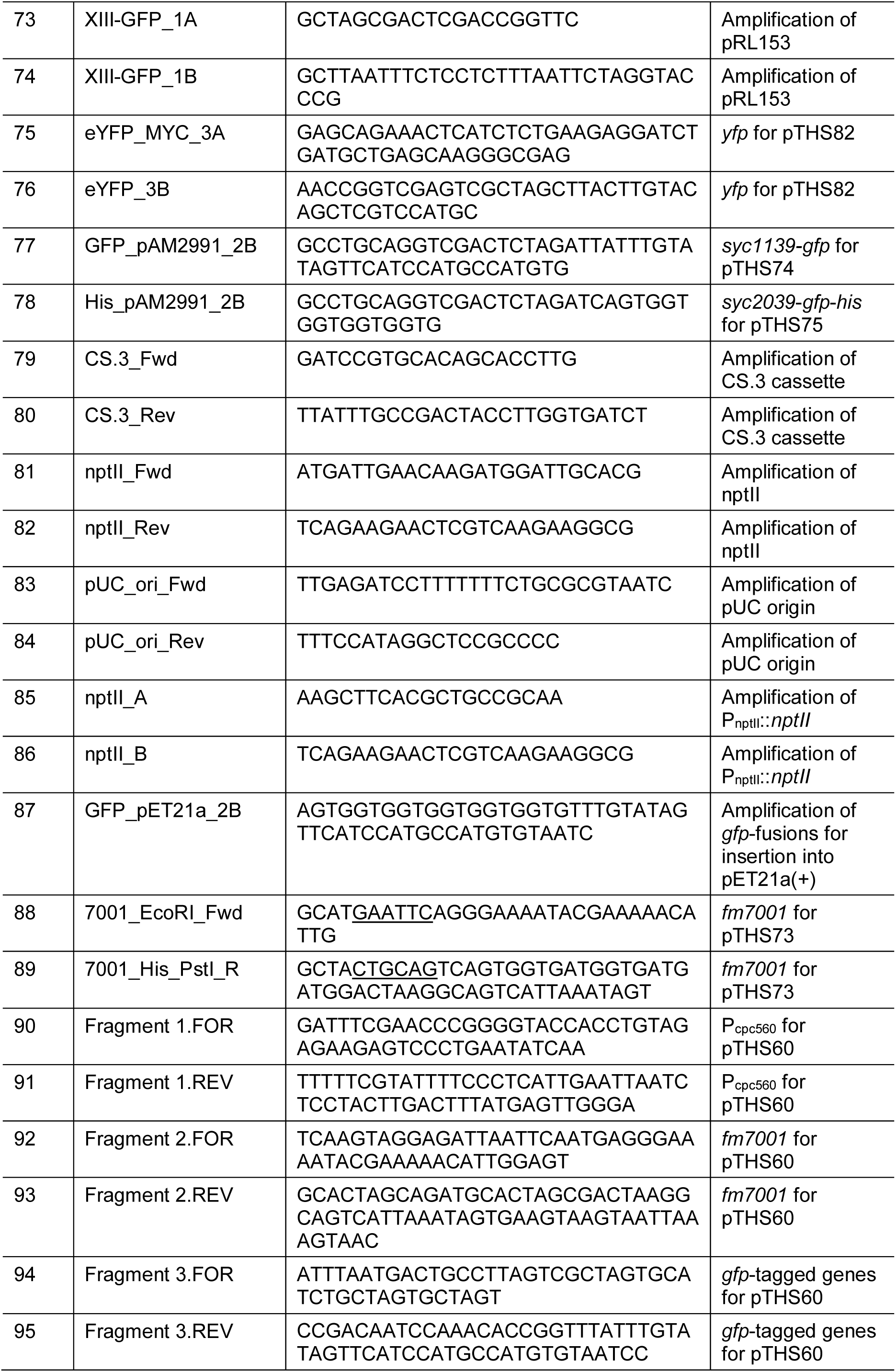
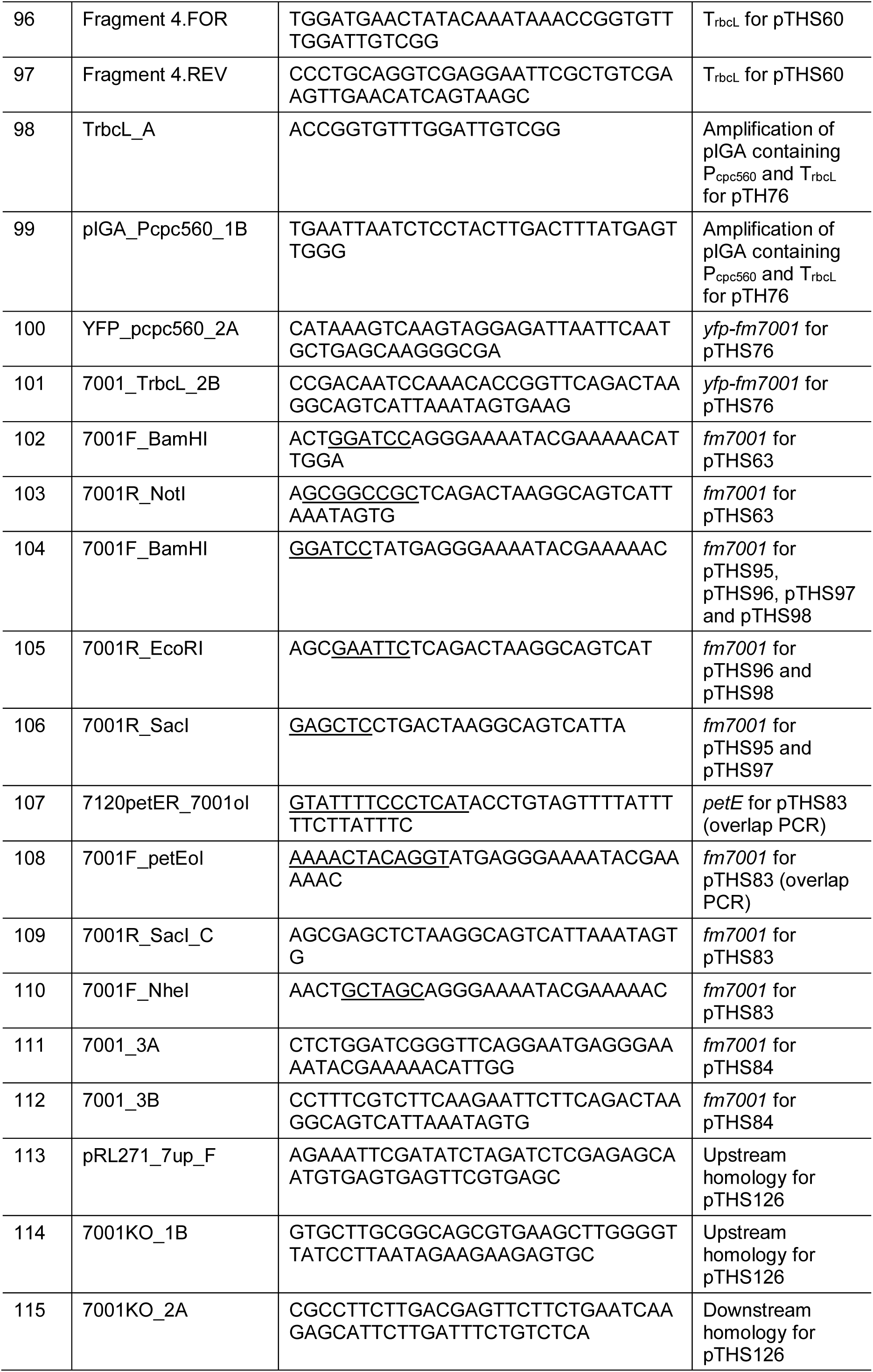
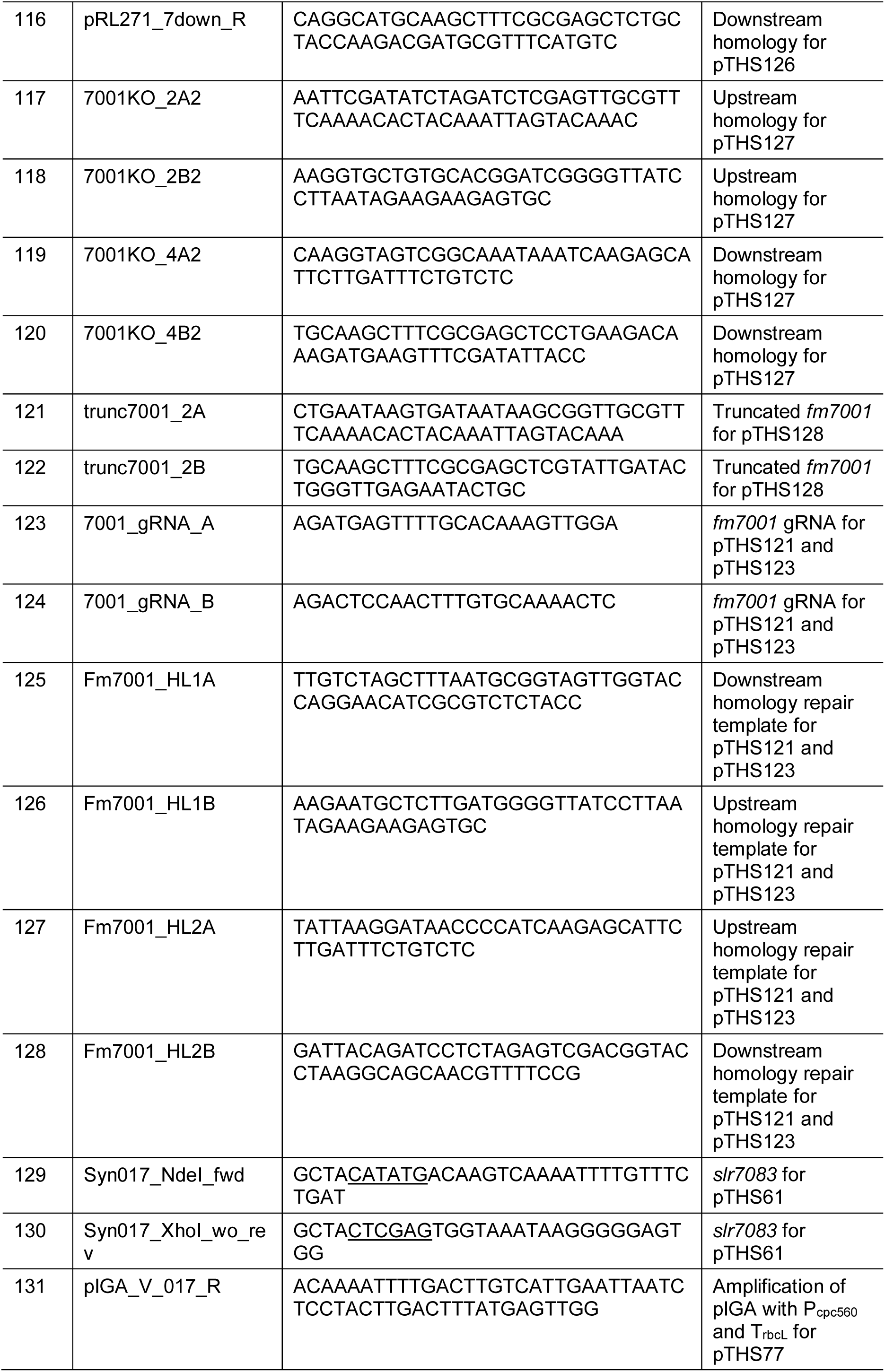
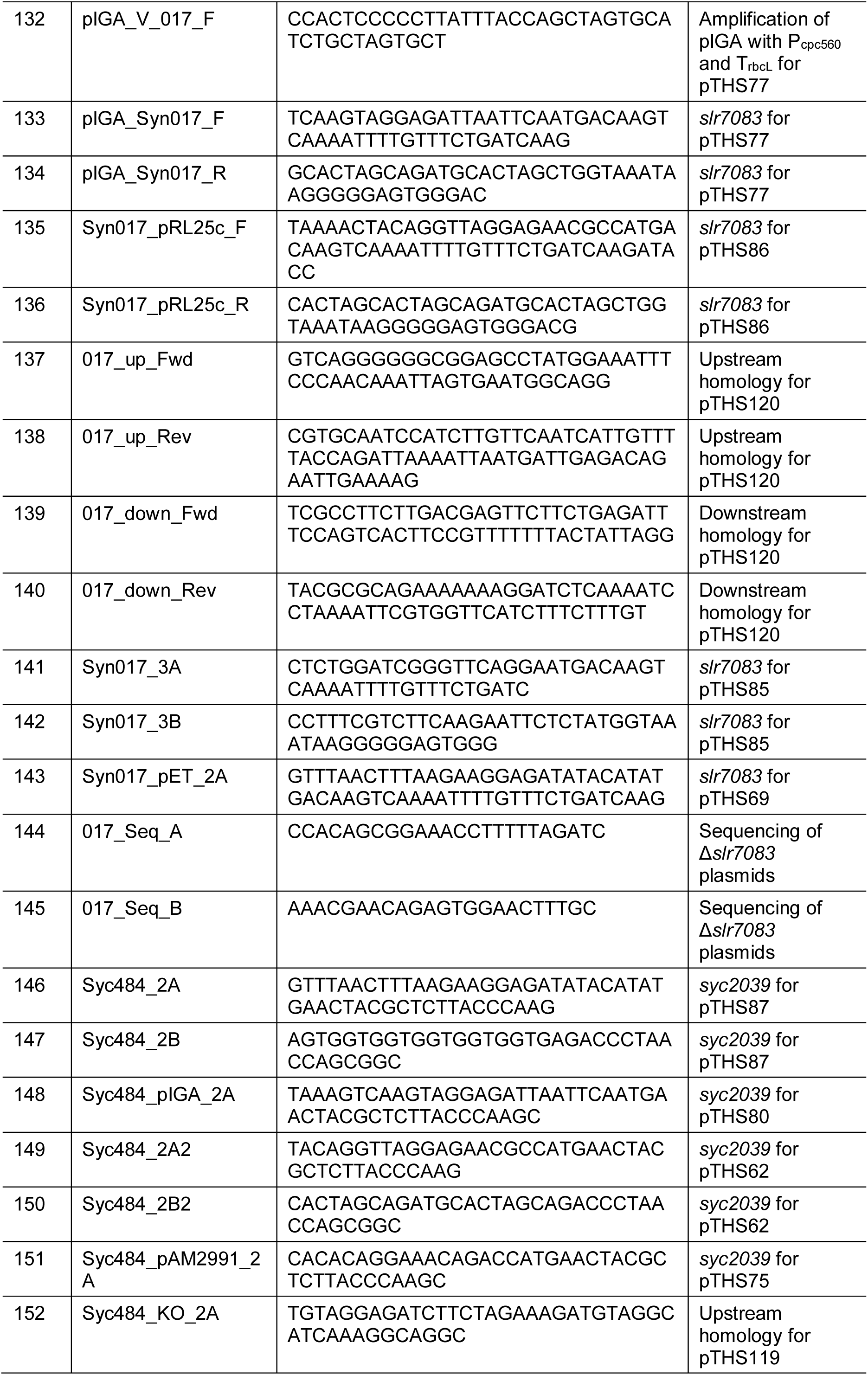
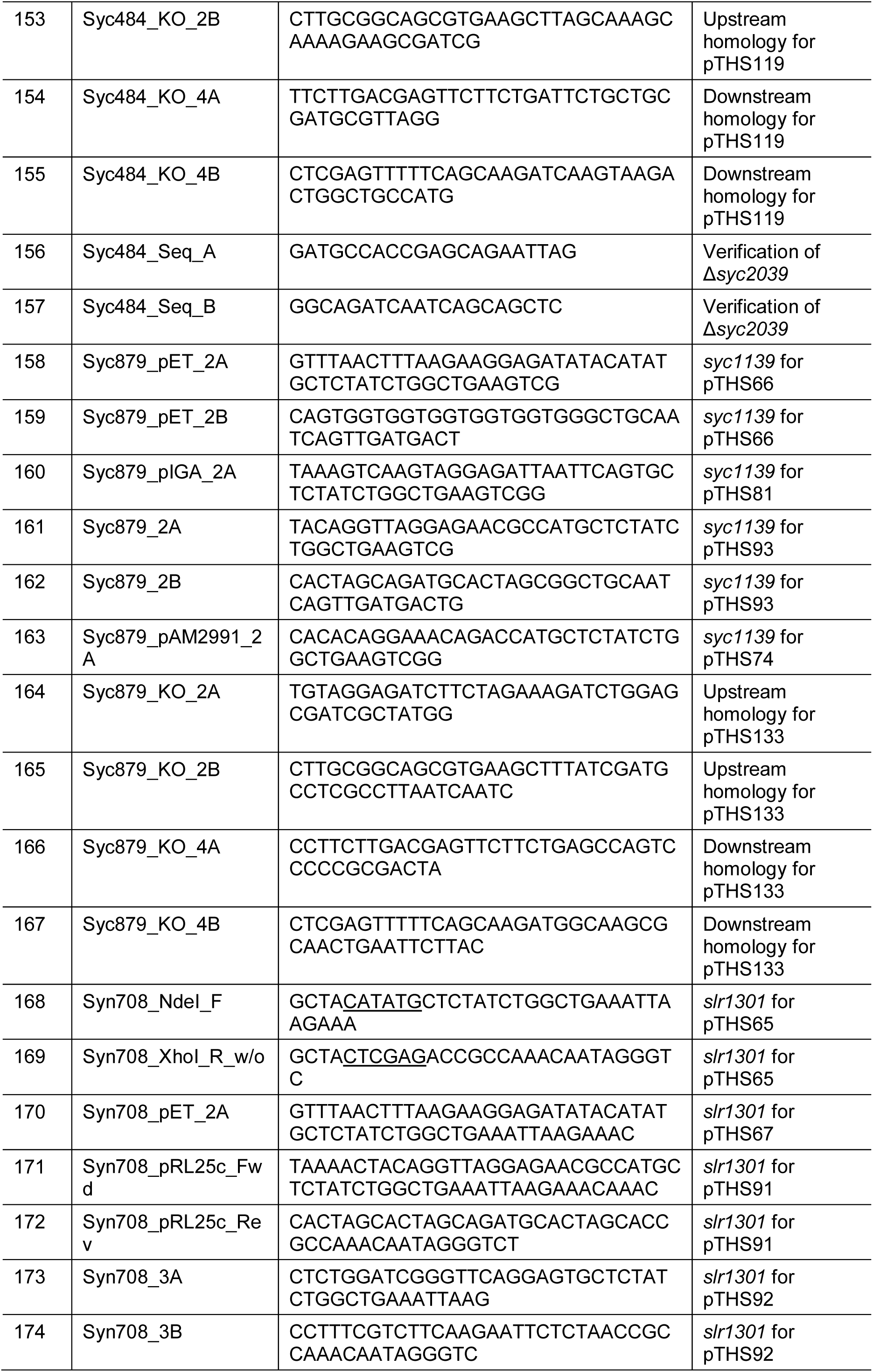
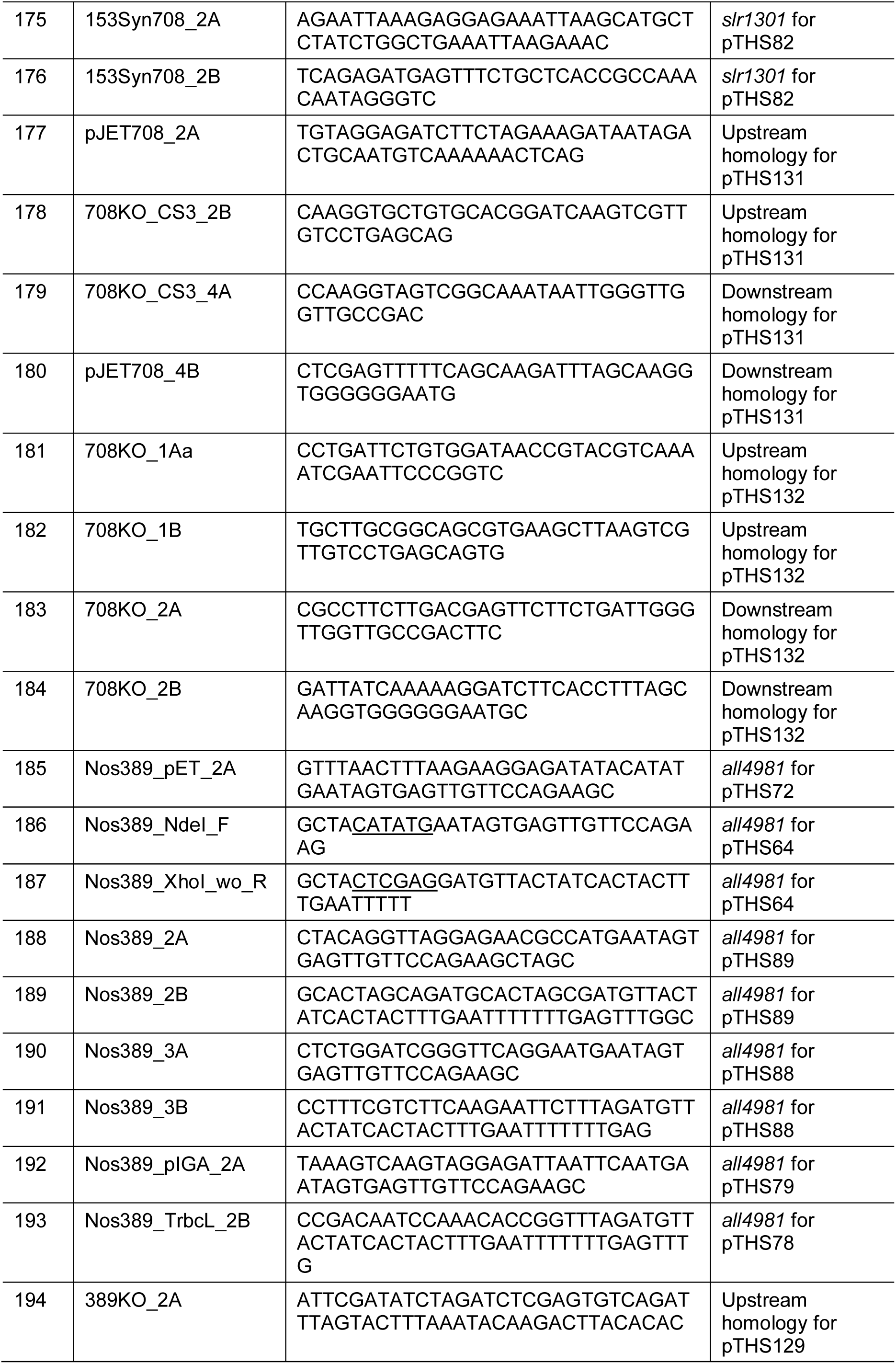
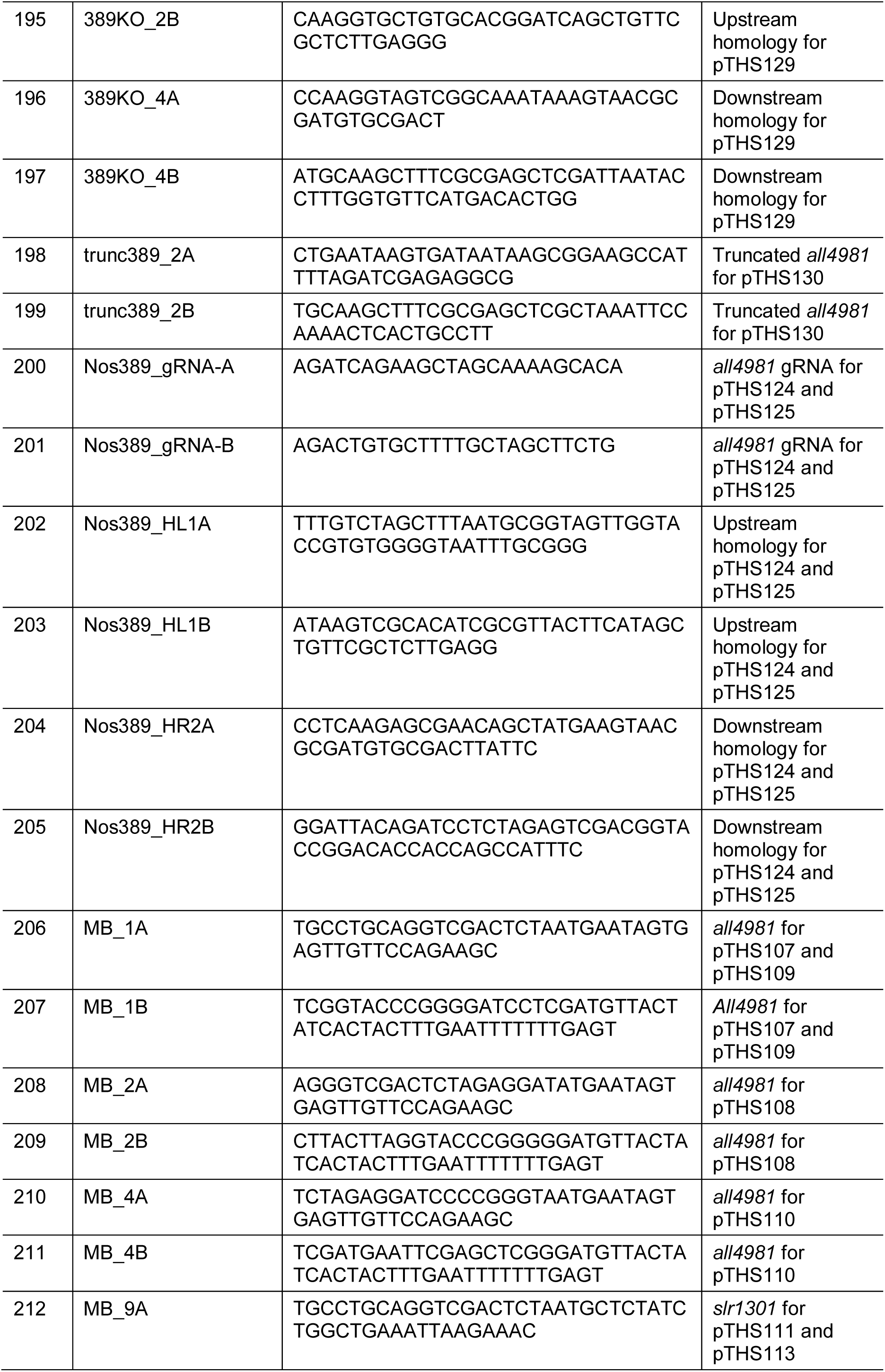
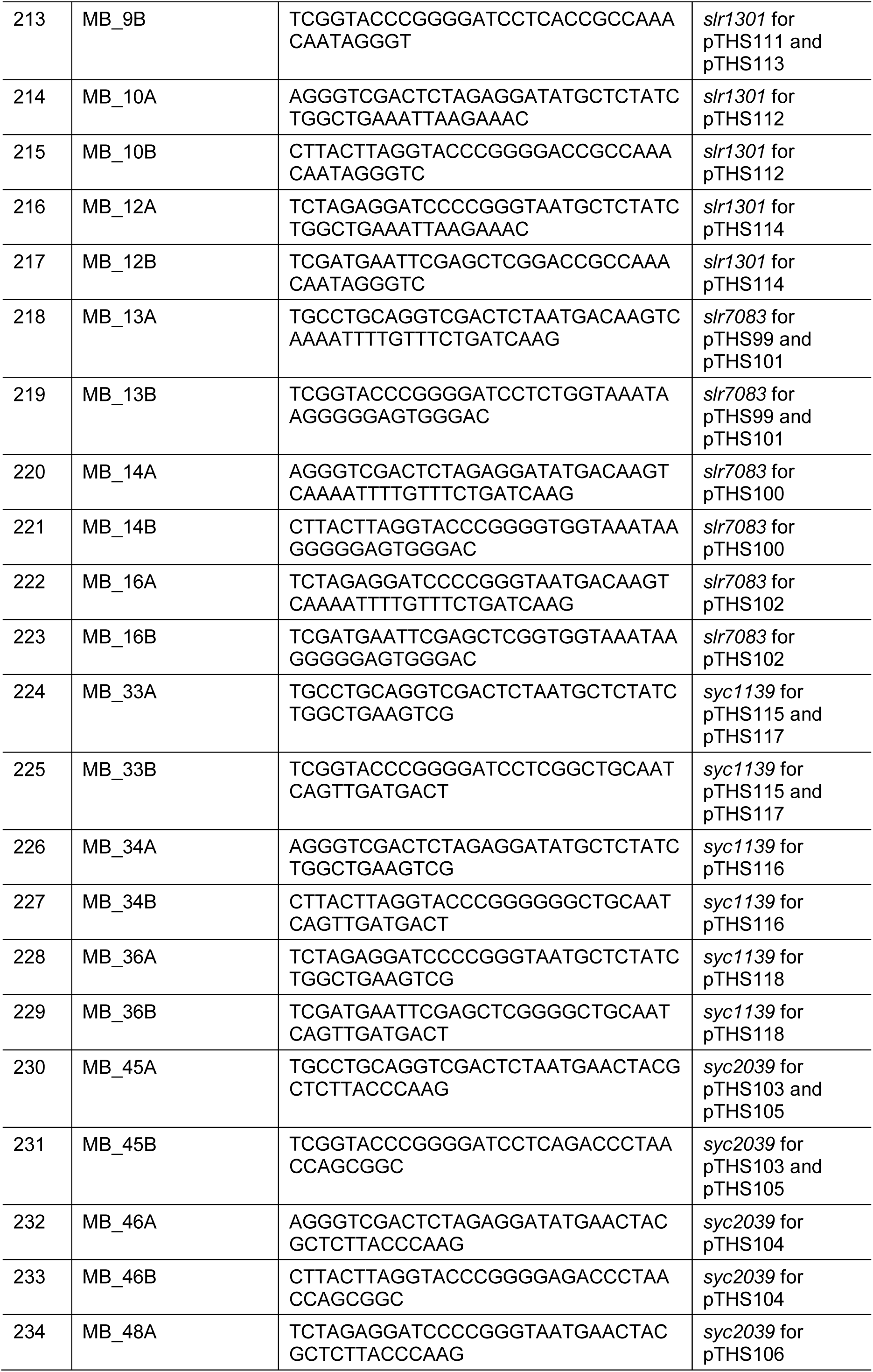
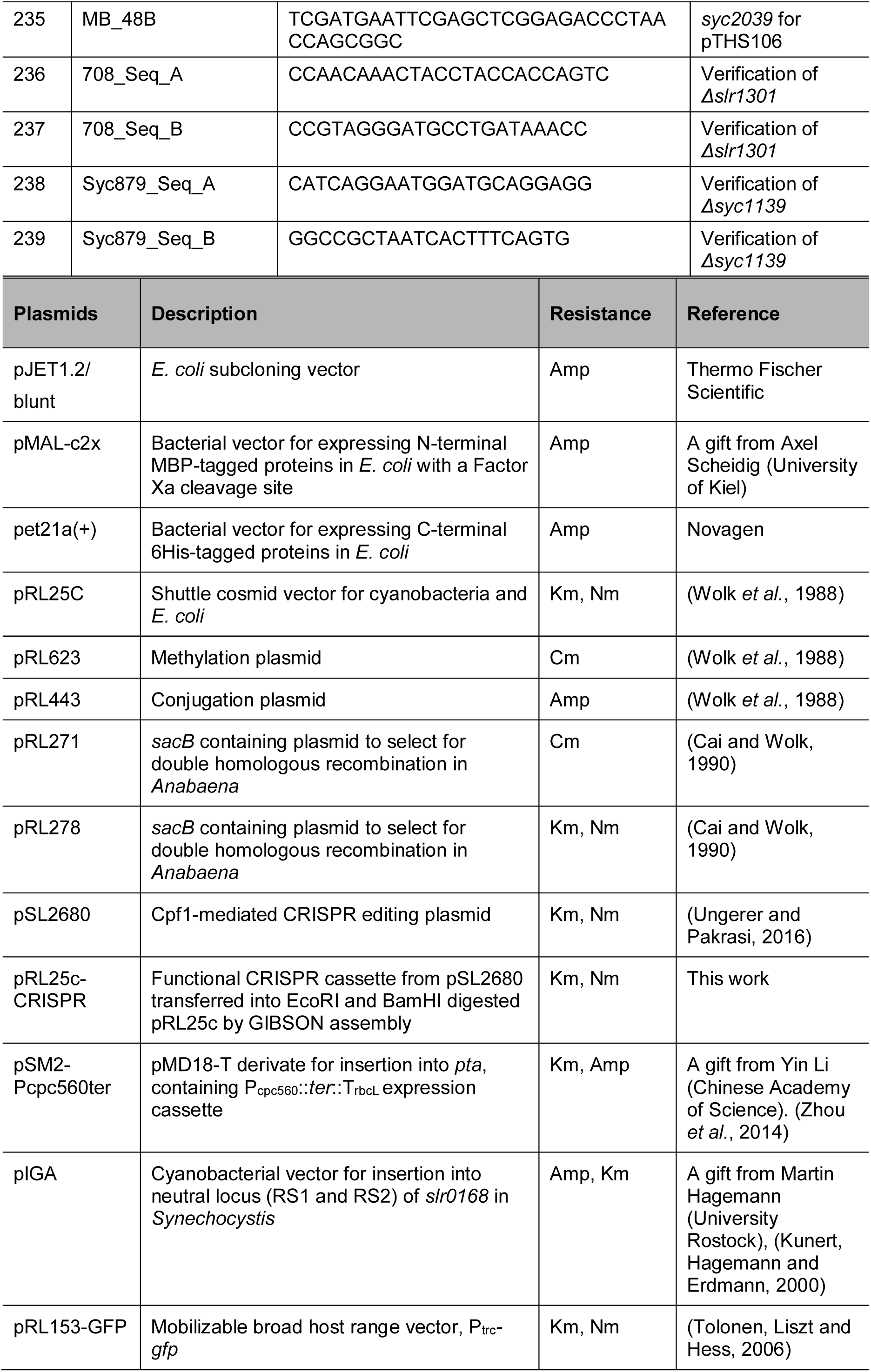
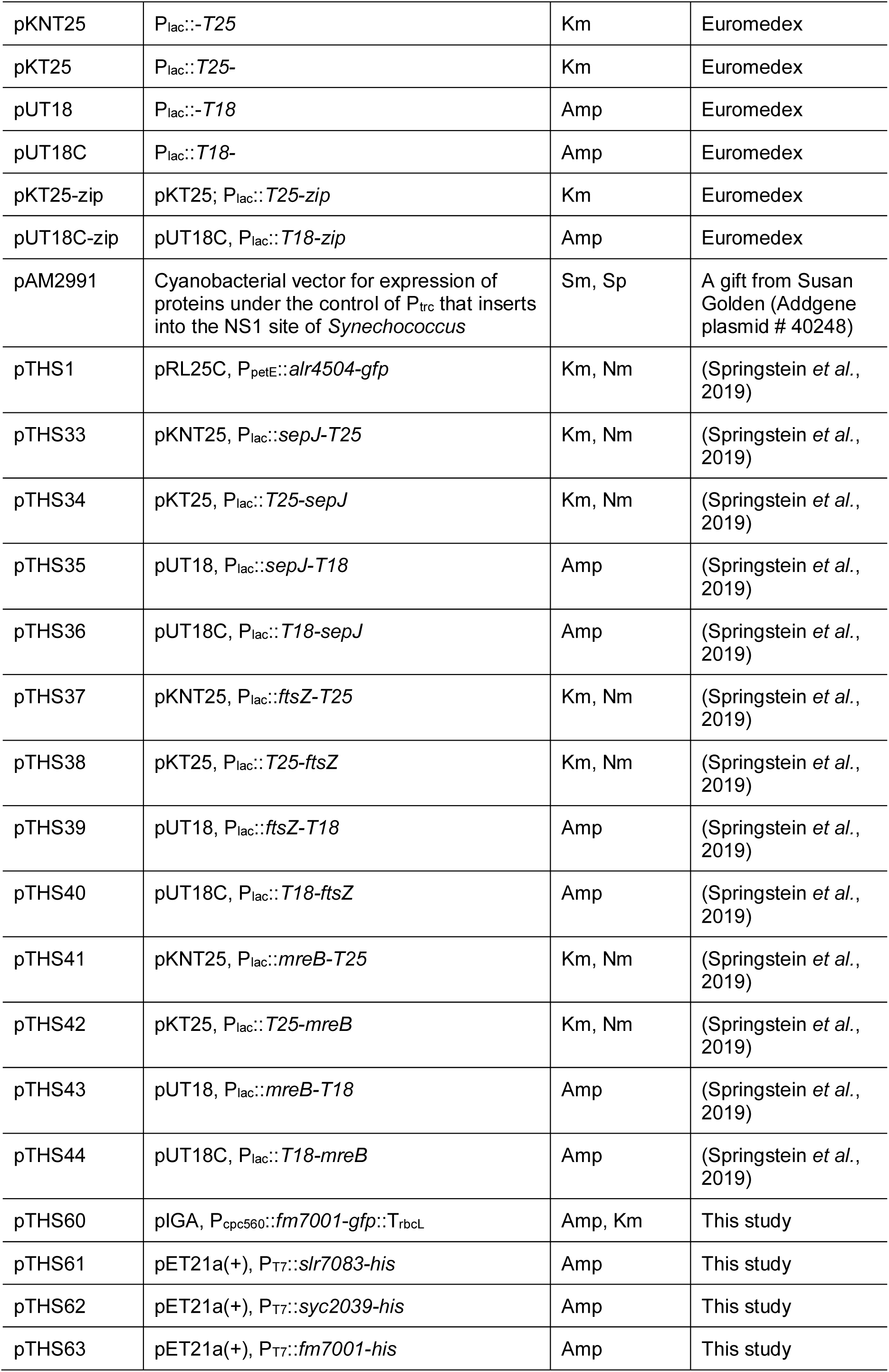
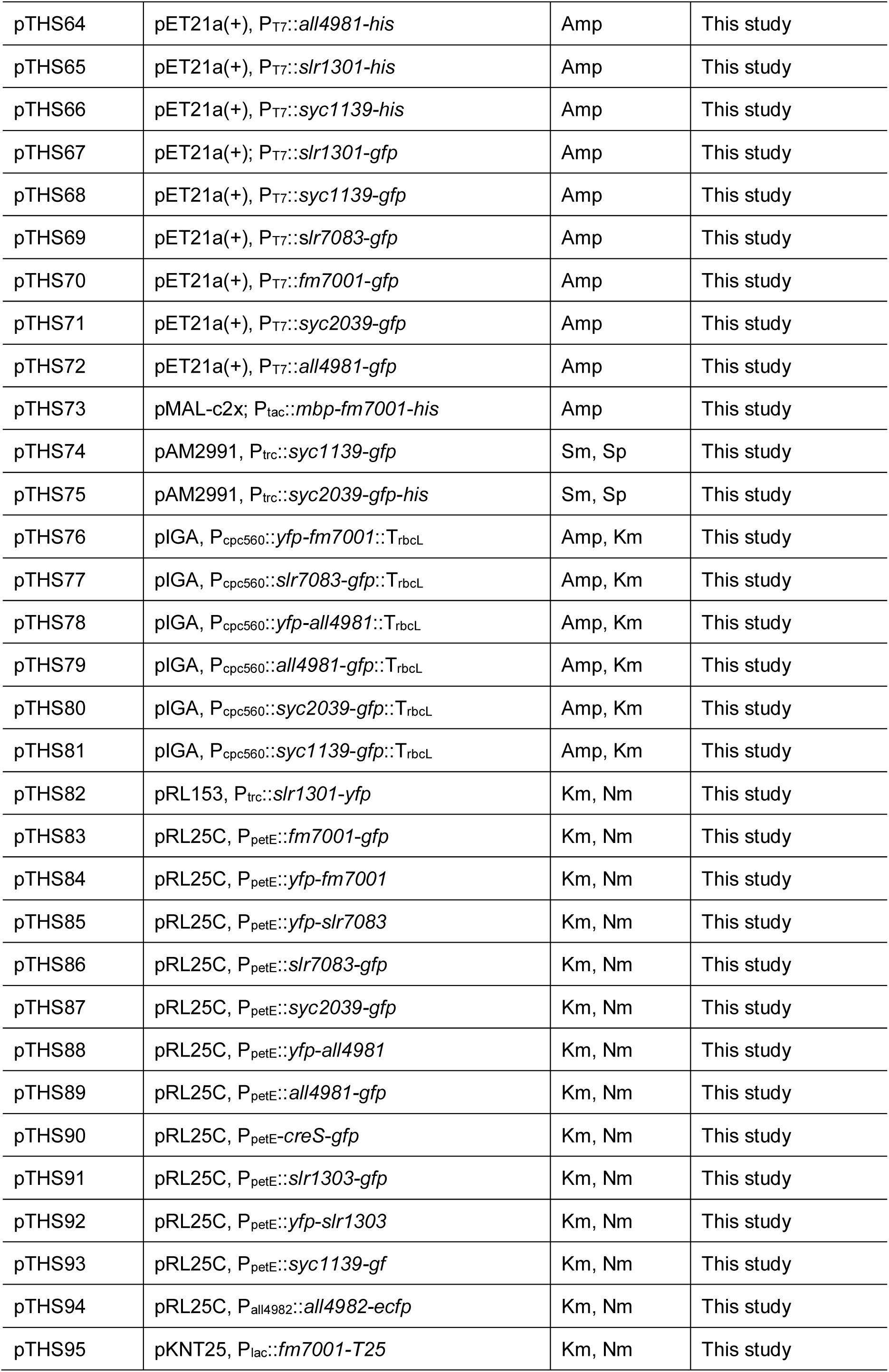
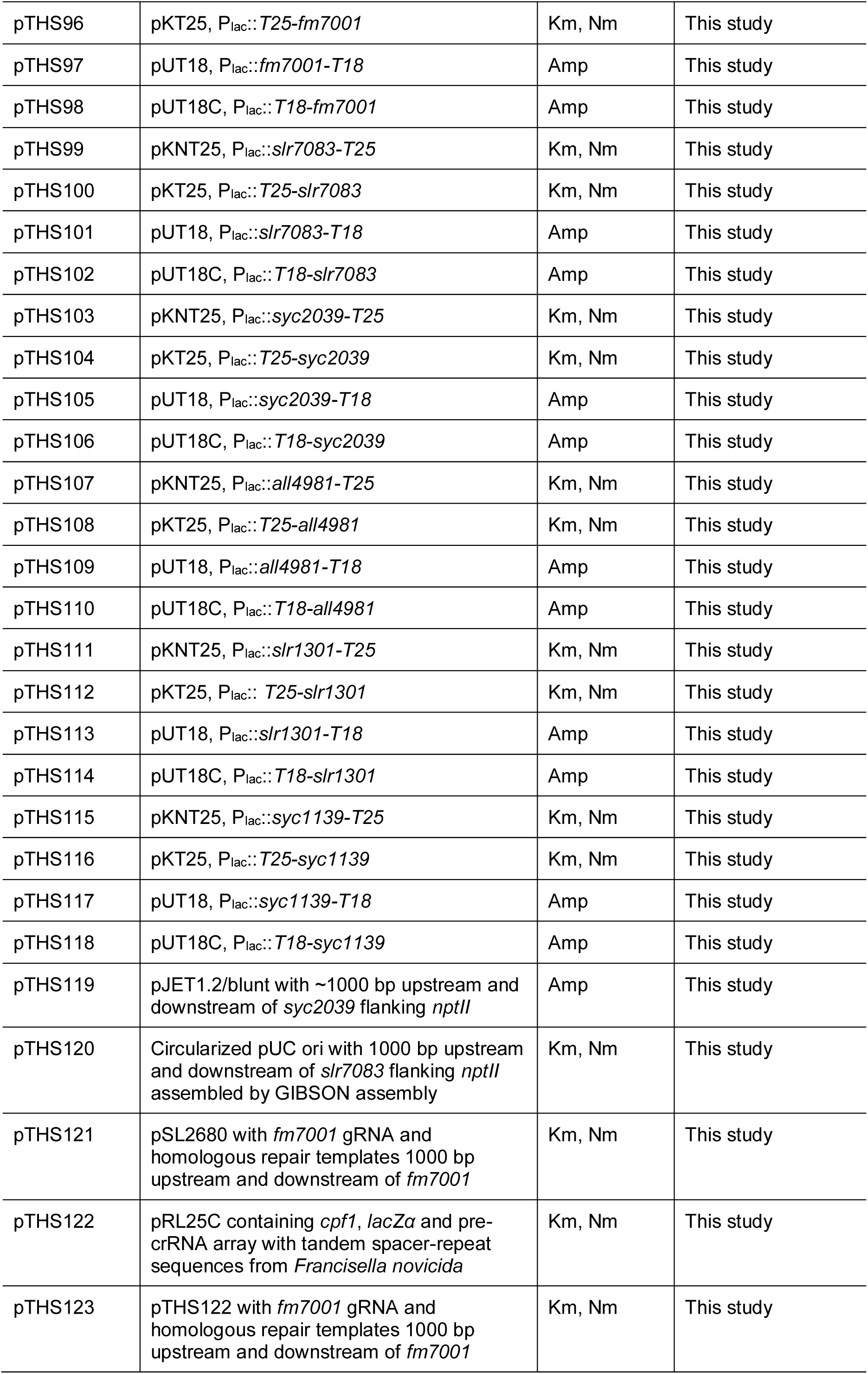
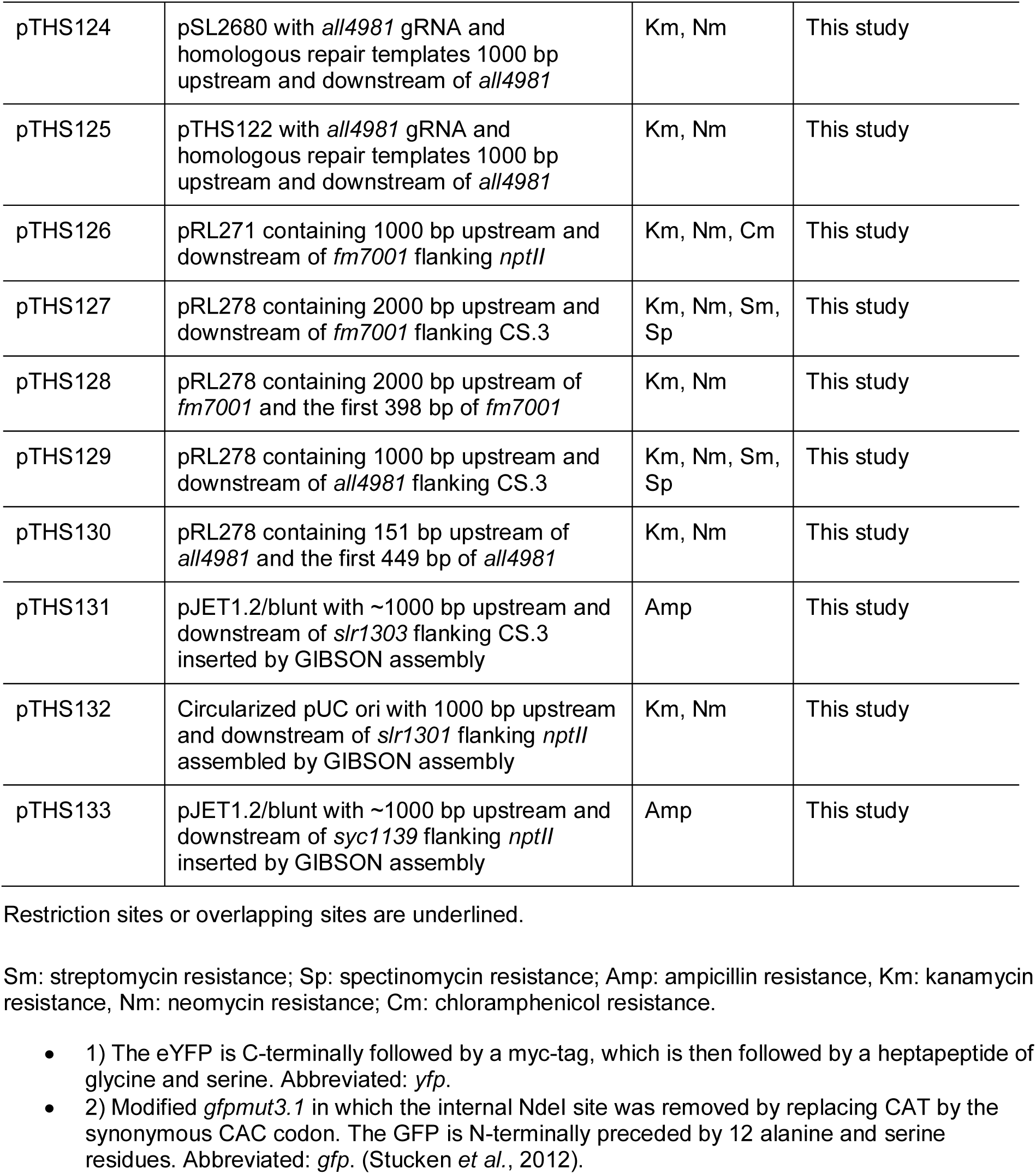
Oligonucleotides and Plasmids.

## References

Alberts, B. et al. (2014) Molecular Biology of the Cell. 6th editio. Garland Science.

Altschul, S. F. et al. (1990) ‘Basic local alignment search tool’, Journal of Molecular Biology. Academic Press, 215(3), pp. 403–410. doi: 10.1016/S0022-2836(05)80360-2.

Ausmees, N., Kuhn, J. R. and Jacobs-Wagner, C. (2003) ‘The bacterial cytoskeleton: An intermediate filament-like function in cell shape’, Cell, pp. 705–713. doi: 10.1016/S0092-8674(03)00935-8.

Bagchi, S. et al. (2008) ‘Intermediate filament-like proteins in bacteria and a cytoskeletal function in Streptomyces’, Molecular Microbiology, 70(4), pp. 1037–1050. doi: 10.1111/j.1365-2958.2008.06473.x.

Beck, E. et al. (1982) ‘Nucleotide sequence and exact localization of the neomycin phosphotransferase gene from transposon Tn5’, Gene, 19(3), pp. 327–336. doi: 10.1016/0378-1119(82)90023-3.

Bharat, T. A. M. et al. (2015) ‘Structures of actin-like ParM filaments show architecture of plasmid-segregating spindles’, Nature. Nature Publishing Group, a division of Macmillan Publishers Limited. All Rights Reserved., 523, p. 106. Available at: https://doi.org/10.1038/nature14356.

Bhaya, D. et al. (2001) ‘Novel Motility Mutants of Synechocystis Strain PCC 6803 Generated by In Vitro Transposon Mutagenesis Novel Motility Mutants of Synechocystis Strain PCC 6803 Generated by In Vitro Transposon Mutagenesis †’, Journal of Bacteriology, 183(20), pp. 1–5. doi: 10.1128/JB.183.20.6140.

Bi, E. and Lutkenhaus, J. (1991) ‘FtsZ ring structure associated with division in Escherichia coli’, Nature, 354(6349), pp. 161–164. doi: 10.1038/354161a0.

Blatch, G. L. and Lässle, M. (1999) ‘The tetratricopeptide repeat: A structural motif mediating protein-protein interactions’, BioEssays, 21(11), pp. 932–939. doi: 10.1002/(SICI)1521-1878(199911)21:11<932::AID-BIES5>3.0.CO;2-N.

Bocharova, O. V. et al. (2005) ‘In vitro conversion of full-length mammalian prion protein produces amyloid form with physical properties of PrPSc’, Journal of Molecular Biology, 346(2), pp. 645–659. doi: 10.1016/j.jmb.2004.11.068.

Boyer, H. and Roulland-Dessoix, D. (1969) ‘A complementation analysis of the restrcition and modification of DNA in Escherichia coli.’, J. Mol. Biol., 41, pp. 459–472.

Buikema, W. J. and Haselkorn, R. (2001) ‘Expression of the Anabaena hetR gene from a copper-regulated promoter leads to heterocyst differentiation under repressing conditions’, Proceedings of the National Academy of Sciences, 98(5), pp. 2729–2734. doi: 10.1073/pnas.051624898.

Bustos, S. A. and Golden, S. S. (1992) ‘Light-regulated expression of the psbD gene family in Synecbococcus sp. strain PCC 7942: evidence for the role of duplicated psbD genes in cyanobacteria’, Molecular and General Genetics MGG, 232(2), pp. 221–230. doi: 10.1007/BF00280000.

Cabeen, M. T. et al. (2009) ‘Bacterial cell curvature through mechanical control of cell growth’, The EMBO Journal, pp. 1208–1219. doi: 10.1038/emboj.2009.61.

Cai, Y. and Wolk, C. P. (1990) ‘Use of a conditionally lethal gene in Anabaena sp. strain PCC 7120 to select for double recombinats and to entrap insertion sequnces’, Journal of Bacteriology, 172(6), pp. 3138–3145.

Camberg, J. L., Hoskins, J. R. and Wickner, S. (2009) ‘ClpXP protease degrades the cytoskeletal protein, FtsZ, and modulates FtsZ polymer dynamics’, Proc Natl Acad Sci U S A, 106(26), pp. 10614–10619. doi: 10.1073/pnas.0904886106.

Cappuccinelli, P. (1980) ‘The movement of eukaryotic cells’, in *Motility of Living Cells*. Dordrecht: Springer Netherlands, pp. 59–74. doi: 10.1007/978-94-009-5812-8_4.

Carballido-Lopez, R. (2006) ‘The Bacterial Actin-Like Cytoskeleton’, Microbiology and Molecular Biology Reviews, 70(4), pp. 888–909. doi: 10.1128/MMBR.00014-06.

Charbon, G., Cabeen, M. T. and Jacobs-Wagner, C. (2009) ‘Bacterial intermediate filaments: In vivo assembly, organization, and dynamics of crescentin’, Genes and Development, pp. 1131–1144. doi: 10.1101/gad.1795509.

Corpet, F. (1988) ‘Multiple sequence alignment with hierarchical clustering.’, Nucleic acids research, 16(22), pp. 10881–90. doi: 10.1093/nar/16.22.10881.

Dörrich, A. K. et al. (2014) ‘Deletion of the Synechocystis sp. PCC 6803 kaiAB1C1 gene cluster causes impaired cell growth under light???dark conditions’, Microbiology (United Kingdom), 160(2014), pp. 2538–2550. doi: 10.1099/mic.0.081695-0.

England, P. et al. (2005) ‘The Scc Spirochetal Coiled-Coil Protein Forms Helix-Like Filaments and Binds to Nucleic Acids Generating Nucleoprotein Structures’, Journal of Bacteriology, 188(2), pp. 469–476. doi: 10.1128/jb.188.2.469-476.2006.

Enright, A. J., Van Dongen, S. and Ouzounis, C. A. (2002) ‘An efficient algorithm for large-scale detection of protein families’, Nucleic acids research. Oxford University Press, 30(7), pp. 1575–1584.

Fiuza, M. et al. (2010) ‘Phosphorylation of a novel cytoskeletal protein (RsmP) regulates rod-shaped morphology in Corynebacterium glutamicum’, Journal of Biological Chemistry, pp. 29387–29397. doi: 10.1074/jbc.M110.154427.

Flärdh, K. et al. (2012) ‘Regulation of apical growth and hyphal branching in Streptomyces’, Current Opinion in Microbiology, 15(6), pp. 737–743. doi: https://doi.org/10.1016/j.mib.2012.10.012.

Flores, E. et al. (2007) ‘Septum-localized protein required for filament integrity and diazotrophy in the heterocyst-forming cyanobacterium Anabaena sp. strain PCC 7120’, Journal of Bacteriology, 189(10), pp. 3884–3890. doi: 10.1128/JB.00085-07.

Fuchino, K. et al. (2013) ‘Dynamic gradients of an intermediate filament-like cytoskeleton are recruited by a polarity landmark during apical growth’, Proceedings of the National Academy of Sciences, pp. E1889–E1897. doi: 10.1073/pnas.1305358110.

Fuchs, E. and Weber, K. (1994) ‘INTERMEDIATE FILAMENTS: Structure, Dynamics, Function and Disease’, Annual Review of Biochemistry, 63, pp. 345–382.

Geisler, N. and Weber, K. (1982) ‘The amino acid sequence of chicken muscle desmin provides a common structural model for intermediate filament proteins’, The EMBO journal, 1(12), pp. 1649–1656. Available at: https://www.ncbi.nlm.nih.gov/pubmed/6202512.

Gibson, D. G. et al. (2009) ‘Enzymatic assembly of DNA molecules up to several hundred kilobases’, Nature Methods, 6(5), pp. 343–345. doi: 10.1038/nmeth.1318.

Grant, S. G. et al. (1990) ‘Differential plasmid rescue from transgenic mouse DNAs into Escherichia coli methylation-restriction mutants.’, Proceedings of the National Academy of Sciences, 87(12), pp. 4645–4649. doi: 10.1073/pnas.87.12.4645.

Heinz, S. et al. (2016) ‘Thylakoid Membrane Architecture in *Synechocystis* Depends on CurT, a Homolog of the Granal CURVATURE THYLAKOID1 Proteins’, The Plant Cell, 28(9), pp. 2238–2260. doi: 10.1105/tpc.16.00491.

Hempel, A. M. et al. (2012) ‘The Ser/Thr protein kinase AfsK regulates polar growth and hyphal branching in the filamentous bacteria Streptomyces’, Proceedings of the National Academy of Sciences of the United States of America. 2012/08/06. National Academy of Sciences, 109(35), pp. E2371–E2379. doi: 10.1073/pnas.1207409109.

Herrero, A., Stavans, J. and Flores, E. (2016) ‘The multicellular nature of filamentous heterocyst-forming cyanobacteria’, FEMS Microbiology Reviews, 40(6), pp. 831–854. doi: 10.1093/femsre/fuw029.

Herrmann, H. et al. (1996) ‘Structure and assembly properties of the intermediate filament protein vimentin: The role of its head, rod and tail domains’, Journal of Molecular Biology, 264(5), pp. 933–953. doi: 10.1006/jmbi.1996.0688.

Herrmann, H. and Aebi, U. (2004) ‘Intermediate Filaments: Molecular Structure, Assembly Mechanism, and Integration Into Functionally Distinct Intracellular Scaffolds’, Annual Review of Biochemistry, 73(1), pp. 749–789. doi: 10.1146/annurev.biochem.73.011303.073823.

Holmes, N. A. et al. (2013) ‘Coiled-coil protein Scy is a key component of a multiprotein assembly controlling polarized growth in Streptomyces’, Proceedings of the National Academy of Sciences, pp. E397–E406. doi: 10.1073/pnas.1210657110.

Hu, B. et al. (2007) ‘MreB is important for cell shape but not for chromosome segregation of the filamentous cyanobacterium Anabaena sp. PCC 7120’, Molecular Microbiology, 63(6), pp. 1640–1652. doi: 10.1111/j.1365-2958.2007.05618.x.

Huang, H.-H. et al. (2010) ‘Design and characterization of molecular tools for a Synthetic Biology approach towards developing cyanobacterial biotechnology’, Nucleic acids research. 2010/03/17. Oxford University Press, 38(8), pp. 2577–2593. doi: 10.1093/nar/gkq164.

Hurme, R. et al. (1994) ‘Intermediate filament-like network formed in vitro by a bacterial coiled coil protein’, Journal of Biological Chemistry, pp. 10675–10682.

Inclan, Y. F. et al. (2016) ‘A scaffold protein connects type IV pili with the Chp chemosensory system to mediate activation of virulence signaling in Pseudomonas aeruginosa’, Molecular Microbiology, 101(4), pp. 590–605. doi: 10.1111/mmi.13410.

Ingerson-Mahar, M. et al. (2010) ‘The metabolic enzyme CTP synthase forms cytoskeletal filaments’, Nature Cell Biology, pp. 739–746. doi: 10.1038/ncb2087.

Ingerson-Mahar, M. and Gitai, Z. (2012) ‘A growing family: the expanding universe of the bacterial cytoskeleton’, FEMS Microbiol Rev, 36(1), pp. 256–266. doi: 10.1111/j.1574-6976.2011.00316.x.

Ivleva, N. B. et al. (2005) ‘LdpA: A component of the circadian clock senses redox state of the cell’, EMBO Journal, 24(6), pp. 1202–1210. doi: 10.1038/sj.emboj.7600606.

Jain, I. H., Vijayan, V. and O’Shea, E. K. (2012) ‘Spatial ordering of chromosomes enhances the fidelity of chromosome partitioning in cyanobacteria.’, Proceedings of the National Academy of Sciences of the United States of America, 109(34), pp. 13638–43. doi: 10.1073/pnas.1211144109.

Jones, L. J. F., Carballido-López, R. and Errington, J. (2001) ‘Control of cell shape in bacteria: Helical, actin-like filaments in Bacillus subtilis’, Cell, 104(6), pp. 913–922. doi: 10.1016/S0092-8674(01)00287-2.

Kahnt, J. et al. (2007) ‘Post translational modifications in the active site region of methylcoenzyme M reductase from methanogenic and methanotrophic archaea’, The FEBS journal. Wiley Online Library, 274(18), pp. 4913–4921.

Karimova, G., Davi, M. and Ladant, D. (2012) ‘The β-lactam resistance protein Blr, a small membrane polypeptide, is a component of the Escherichia coli cell division machinery’, Journal of Bacteriology, 194(20), pp. 5576–5588. doi: 10.1128/JB.00774-12.

Kelemen, G. H. (2017) ‘Intermediate Filaments Supporting Cell Shape and Growth in Bacteria BT - Prokaryotic Cytoskeletons: Filamentous Protein Polymers Active in the Cytoplasm of Bacterial and Archaeal Cells’, in Löwe, J. and Amos, L. A. (eds). Cham: Springer International Publishing, pp. 161–211. doi: 10.1007/978-3-319-53047-5_6.

Koch, M. K., McHugh, C. A. and Hoiczyk, E. (2011) ‘BacM, an N-terminally processed bactofilin of Myxococcus xanthus, is crucial for proper cell shape’, Mol Microbiol., 80(4), pp. 1031–1051. doi: 10.1111/j.1365-2958.2011.07629.x.

Kopfmann, S. and Hess, W. R. (2013) ‘Toxin-antitoxin systems on the large defense plasmid pSYSA of synechocystis sp. pCC 6803’, Journal of Biological Chemistry, 288(10), pp. 7399–7409. doi: 10.1074/jbc.M112.434100.

Köster, S. et al. (2015) ‘Intermediate filament mechanics in vitro and in the cell: From coiled coils to filaments, fibers and networks’, Current Opinion in Cell Biology, 32, pp. 82–91. doi: 10.1016/j.ceb.2015.01.001.

Krogh, A. et al. (2001) ‘Predicting transmembrane protein topology with a hidden Markov model: application to complete genomes’, Journal of molecular biology. Elsevier, 305(3), pp. 567–580.

Kruse, T., Bork-Jensen, J. and Gerdes, K. (2005) ‘The morphogenetic MreBCD proteins of Escherichia coli form an essential membrane-bound complex’, Molecular Microbiology, 55(1), pp. 78–89. doi: 10.1111/j.1365-2958.2004.04367.x.

Kühn, J. et al. (2010) ‘Bactofilins, a ubiquitous class of cytoskeletal proteins mediating polar localization of a cell wall synthase in Caulobacter crescentus’, EMBO Journal, pp. 327–339. doi: 10.1038/emboj.2009.358.

Kunert, A., Hagemann, M. and Erdmann, N. (2000) ‘Construction of promoter probe vectors for Synechocystis sp. PCC 6803 using the light-emitting reporter systems Gfp and LuxAB’, Journal of Microbiological Methods, 41(3), pp. 185–194. doi: 10.1016/S0167-7012(00)00162-7.

Larsen, R. A. et al. (2007) ‘Treadmilling of a prokaryotic tubulin-like protein, TubZ, required for plasmid stability in Bacillus thuringiensis’, Genes and Development, 21(11), pp. 1340–1352. doi: 10.1101/gad.1546107.

Leung, C. L., Green, K. J. and Liem, R. K. H. (2002) ‘Plakins: A family of versatile cytolinker proteins’, Trends in Cell Biology, 12(1), pp. 37–45. doi: 10.1016/S0962-8924(01)02180-8.

Lin, L. and Thanbichler, M. (2013) ‘Nucleotide-independent cytoskeletal scaffolds in bacteria’, Cytoskeleton, 70(8), pp. 409–423. doi: 10.1002/cm.21126.

Lodish, H. et al. (2000) Molecular Cell Biology. 4th edn. New York: W. H. Freeman. Available at: https://www.ncbi.nlm.nih.gov/books/NBK21560/ (Accessed: 27 February 2018).

Lopes Pinto, F. et al. (2011) ‘FtsZ degradation in the cyanobacterium Anabaena sp. strain PCC 7120’, Journal of Plant Physiology. Elsevier GmbH., 168(16), pp. 1934–1942. doi: 10.1016/j.jplph.2011.05.023.

Löwe, J. and Amos, L. A. (2009) ‘Evolution of cytomotive filaments: The cytoskeleton from prokaryotes to eukaryotes’, International Journal of Biochemistry and Cell Biology, 41(2), pp. 323–329. doi: 10.1016/j.biocel.2008.08.010.

Lupas, A., Van Dyke, M. and Stock, J. (1991) ‘Predicting coiled coils from protein sequences’, Science, pp. 1162–1164. doi: 10.1126/science.252.5009.1162.

Marchler-Bauer, A. et al. (2016) ‘CDD/SPARCLE: functional classification of proteins via subfamily domain architectures’, Nucleic acids research. Oxford University Press, 45(D1), pp. D200–D203.

McGuffin, L. J., Bryson, K. and Jones, D. T. (2000) ‘The PSIPRED protein structure prediction server’, Bioinformatics. Oxford University Press, 16(4), pp. 404–405.

Meselson, M. and Yuan, R. (1968) ‘DNA restriction enzyme from E. coli.’, Nature, 217(5134), pp. 1110–4. Available at: http://www.ncbi.nlm.nih.gov/pubmed/4868368 (Accessed: 22 February 2018).

Miller, J. H. (1992) A Short Course in Bacterial Genetics – A Laboratory Manual and Handbook for Escherichia coli and Related Bacteria, Cold Spring Harbor Laboratory Press. Cold Spring Harbor. doi: 10.1002/jobm.3620330412.

Nakamura, Y. et al. (1993) ‘Acceleration of bovine neurofilament L assembly by deprivation of acidic tail domain’, European Journal of Biochemistry, 212(2), pp. 565–571. doi: 10.1111/j.1432-1033.1993.tb17694.x.

Nan, B. et al. (2010) ‘A multi-protein complex from Myxococcus xanthus required for bacterial gliding motility’, Molecular Microbiology, 76(6), pp. 1539–1554. doi: 10.1111/j.1365-2958.2010.07184.x.

Nayar, A. S. et al. (2007) ‘FraG is necessary for filament integrity and heterocyst maturation in the cyanobacterium Anabaena sp. strain PCC 7120’, Microbiology, 153(2), pp. 601–607. doi: 10.1099/mic.0.2006/002535-0.

O’Leary, N. A. et al. (2015) ‘Reference sequence (RefSeq) database at NCBI: current status, taxonomic expansion, and functional annotation’, Nucleic acids research. Oxford University Press, 44(D1), pp. D733–D745.

Rackham, O. J. L. et al. (2010) ‘The Evolution and Structure Prediction of Coiled Coils across All Genomes’, Journal of Molecular Biology. Academic Press, 403(3), pp. 480–493. doi: 10.1016/J.JMB.2010.08.032.

Ramos-León, F. et al. (2015) ‘Divisome-dependent subcellular localization of cell-cell joining protein SepJ in the filamentous cyanobacterium Anabaena’, Molecular Microbiology, 96(3), pp. 566–580. doi: 10.1111/mmi.12956.

Rice, P., Longden, I. and Bleasby, A. (2000) ‘EMBOSS: the European Molecular Biology Open Software Suite.’, Trends in genetics : TIG. Elsevier, 16(6), pp. 276–7. doi: 10.1016/S0168-9525(00)02024-2.

Rippka, R. et al. (1979) ‘Generic Assignments, Strain Histories and Properties of Pure Cultures of Cyanobacteria’, Microbiology, 111(1), pp. 1–61. doi: 10.1099/00221287-111-1-1.

Sandvang, D. (1999) ‘Novel streptomycin and spectinomycin resistance gene as a gene cassette within a class 1 integron isolated from Escherichia coli’, Antimicrobial Agents and Chemotherapy, 43(12), pp. 3036–3038. doi: 10.1128/AAC.44.2.475-475.2000.

Schuergers, N. et al. (2015) ‘PilB localization correlates with the direction of twitching motility in the cyanobacterium Synechocystis sp. PCC 6803’, Microbiology (Reading, England), 161(2015), pp. 960–966. doi: 10.1099/mic.0.000064.

Schuergers, N., Mullineaux, C. W. and Wilde, A. (2017) ‘Cyanobacteria in motion’, Current Opinion in Plant Biology. Elsevier Ltd, 37, pp. 109–115. doi: 10.1016/j.pbi.2017.03.018.

Shoeman, R. L. and Traub, P. (1993) ‘Assembly of Intermediate Filaments’, Bioessays, 15(9), pp. 605–611. doi: 10.1002/bies.950150906.

Specht, M. et al. (2011) ‘Helicobacter pylori Possesses Four Coiled-Coil-Rich Proteins That Form Extended Filamentous Structures and Control Cell Shape and Motility’, Journal of Bacteriology, 193(17), pp. 4523–4530. doi: 10.1128/JB.00231-11.

Springstein, B. L. et al. (2019) ‘A cytoskeletal network maintains filament shape in the multicellular cyanobacterium Anabaena sp. PCC 7120’, bioRxiv, p. 553073. doi: 10.1101/553073.

Stucken, K. et al. (2012) ‘Transformation and conjugal transfer of foreign genes into the filamentous multicellular cyanobacteria (subsection V) Fischerella and Chlorogloeopsis’, Current Microbiology, 65(5), pp. 552–560. doi: 10.1007/s00284-012-0193-5.

Stucken, K., Koch, R. and Dagan, T. (2013) ‘Cyanobacterial defense mechanisms against foreign DNA transfer and their impact on genetic engineering’, Biological Research, 46(4), pp. 373–382. doi: 10.4067/S0716-97602013000400009.

Studier, F. W. and Moffatt, B. A. (1986) ‘Use of bacteriophage T7 RNA polymerase to direct selective high-level expression of cloned genes’, Journal of Molecular Biology, 189(1), pp. 113–130. doi: 10.1016/0022-2836(86)90385-2.

Stuurman, N., Heins, S. and Aebi, U. (1998) ‘Nuclear lamins: Their structure, assembly, and interactions’, Journal of Structural Biology, 122(1–2), pp. 42–66. doi: 10.1006/jsbi.1998.3987.

Swulius, M. T. and Jensen, G. J. (2012) ‘The helical mreb cytoskeleton in Escherichia coli MC1000/pLE7 is an artifact of the N-terminal yellow fluorescent protein tag’, Journal of Bacteriology, 194(23), pp. 6382–6386. doi: 10.1128/JB.00505-12.

Tatusov, R. L., Koonin, E. V and Lipman, D. J. (1997) ‘A genomic perspective on protein families’, Science. American Association for the Advancement of Science, 278(5338), pp. 631–637.

Tolonen, A. C., Liszt, G. B. and Hess, W. R. (2006) ‘Genetic manipulation of Prochlorococcus strain MIT9313: green fluorescent protein expression from an RSF1010 plasmid and Tn5 transposition’, Applied and environmental microbiology. 2006/10/13. American Society for Microbiology, 72(12), pp. 7607–7613. doi: 10.1128/AEM.02034-06.

Traub, P. and Vorgias, C. E. (1983) ‘Involvement of the N-terminal polypeptide of vimentin in the formation of intermediate filaments’, Journal of Cell Science, 63(1), pp. 43 LP–67. Available at: http://jcs.biologists.org/content/63/1/43.abstract.

Ungerer, J. and Pakrasi, H. B. (2016) ‘Cpf1 Is A Versatile Tool for CRISPR Genome Editing Across Diverse Species of Cyanobacteria’, Scientific Reports. Nature Publishing Group, 6, pp. 1–9. doi: 10.1038/srep39681.

Vermaas, W. F. J. et al. (2002) ‘Transformation of the cyanobacterium Synechocystis sp. PCC 6803 as a tool for genetic mapping: optimization of efficiency’, FEMS Microbiology Letters, 206(2), pp. 215–219. doi: 10.1111/j.1574-6968.2002.tb11012.x.

Wagstaff, J. and Löwe, J. (2018) ‘Prokaryotic cytoskeletons: protein filaments organizing small cells’, Nature Reviews Microbiology. Nature Publishing Group. doi: 10.1038/nrmicro.2017.153.

Waidner, B. et al. (2009) ‘A novel system of cytoskeletal elements in the human pathogen Helicobacter pylori’, PLoS Pathogens. doi: 10.1371/journal.ppat.1000669.

Walshaw, J., Gillespie, M. D. and Kelemen, G. H. (2010) ‘A novel coiled-coil repeat variant in a class of bacterial cytoskeletal proteins’, Journal of Structural Biology, pp. 202–215. doi: 10.1016/j.jsb.2010.02.008.

Weber, K. and Geisler, N. (1982) ‘The structural relation between intermediate filament proteins in living cells and the alpha-keratins of sheep wool’, The EMBO journal, 1(10), pp. 1155–1160. Available at: https://www.ncbi.nlm.nih.gov/pubmed/6202505.

Weiss, G. L. et al. (2019) ‘Structure and Function of a Bacterial Gap Junction Analog’, Cell. Cell Press, 178(2), pp. 374–384.e15. doi: 10.1016/J.CELL.2019.05.055.

Weissenbach, J. et al. (2017) ‘Evolution of Chaperonin Gene Duplication in Stigonematalean Cyanobacteria (Subsection V)’, Genome Biology and Evolution, 9(1), pp. 241–252. doi: 10.1093/gbe/evw287.

Wiche, G., Osmanagic-Myers, S. and Castañón, M. J. (2015) ‘Networking and anchoring through plectin: A key to IF functionality and mechanotransduction’, Current Opinion in Cell Biology, 32, pp. 21–29. doi: 10.1016/j.ceb.2014.10.002.

Wickstead, B. and Gull, K. (2011) ‘The evolution of the cytoskeleton’, Journal of Cell Biology, 194(4), pp. 513–525. doi: 10.1083/jcb.201102065.

Wilk, L. et al. (2011) ‘Outer membrane continuity and septosome formation between vegetative cells in the filaments of Anabaena sp. PCC 7120’, Cellular Microbiology, 13(11), pp. 1744–1754. doi: 10.1111/j.1462-5822.2011.01655.x.

Wolk, C. P. et al. (1988) ‘Isolation and complementation of mutants of Anabaena sp. strain PCC 7120 unable to grow aerobically on dinitrogen.’, Journal of bacteriology, 170(3), pp. 1239–1244. doi: 10.1128/jb.170.3.1239-1244.1988.

Yang, J. and Zhang, Y. (2015) ‘I-TASSER server: New development for protein structure and function predictions’, Nucleic Acids Research, 43(W1), pp. W174–W181. doi: 10.1093/nar/gkv342.

Yang, R. et al. (2004) ‘AglZ Is a Filament-Forming Coiled-Coil Protein Required for Adventurous Gliding Motility of Myxococcus xanthus’, Journal of bacteriology, 186(18), pp. 6168–6178. doi: 10.1128/JB.186.18.6168.

Yu, N. Y. et al. (2010) ‘PSORTb 3.0: Improved protein subcellular localization prediction with refined localization subcategories and predictive capabilities for all prokaryotes’, Bioinformatics, 26(13), pp. 1608–1615. doi: 10.1093/bioinformatics/btq249.

Zhang, C.-C. C. et al. (1995) ‘Analysis of genes encoding the cell division protein FtsZ and a glutathione synthetase homologue in the cyanobacterium Anabaena sp. PCC 7120’, Research in Microbiology, 146(6), pp. 445–455. doi: 10.1016/0923-2508(96)80290-7.

Zhang, Y. (2009) ‘I-TASSER: Fully automated protein structure prediction in CASP8’, *Proteins: Structure*, Function and Bioinformatics, 77(SUPPL. 9), pp. 100–113. doi: 10.1002/prot.22588.

Zhou, J. et al. (2014) ‘Discovery of a super-strong promoter enables efficient production of heterologous proteins in cyanobacteria’, Scientific Reports, 4, pp. 1–6. doi: 10.1038/srep04500.

